# Mastodon: the Command Center for Large-Scale Lineage-Tracing Microscopy Datasets

**DOI:** 10.64898/2025.12.10.693416

**Authors:** Johannes Girstmair, Tobias Pietzsch, Vladimir Ulman, Stefan Hahmann, Matthias Arzt, Mette Handberg-Thorsager, Samuel Pantze, Ko Sugawara, Robert Haase, Jean-Yves Tinevez, Pavel Tomancak

## Abstract

Understanding development in living organisms requires following the divisions, movements, and fates of cells across developing systems. While advances in microscopy have enabled whole-embryo imaging at the cellular level, extracting and analyzing cell lineages from these massive datasets remains a significant computational challenge. We present Mastodon, a scalable, extensible software platform for manual, semi-automated, and automated cell tracking in large images. A purpose-built graph model supports responsive performance for datasets with millions of annotations, making Mastodon a future-proof platform for cell lineage analysis. Built as a Fiji plugin, Mastodon enables interactive visualization, editing, and analysis of complex lineage trees, seamlessly integrated with the raw image data. Comprehension of cell lineages in complex three-dimensional geometries is facilitated by interoperability with the powerful open-source render engine Blender. In three distinct developmental contexts, we demonstrate how Mastodon will accelerate biological insights by providing user-friendly navigation and explorative analysis in complex lineage datasets.

## Main

The unifying feature of multicellular organisms is their origin in a single-cell zygote. The zygote builds organismal form through an interplay of cell divisions, movements, interactions, differentiations, and deaths at least partially orchestrated by the genome. One of the valuable abstractions of this complex biological process has been the extraction and analysis of developmental lineages. In this context, the lineage describes a tree-like hierarchical relationship between cells as they progress through the developmental process, divide, and potentially die. Since the dawn of biology, researchers have meticulously extracted organisms’ lineage trees from direct observations or by labeling and following specific cells. This led to significant insights into cellular and morphological events of development, such as gastrulation and pattern formation (1,2). The most prominent example of lineage tree analysis has been the establishment of the complete organismal lineage in *C. elegans* from microscopic observations of unlabeled embryos (3,4).

In the modern era, the study of cell lineages by direct observation has been fueled by advances in cell labeling and microscopy. In particular, introducing Selective Plane Illumination Microscopy (SPIM) in combination with fluorescent cell marking (5–10) enabled researchers to record cellular-level developmental events at unprecedented scale, precision, and completeness. The sheer size of the microscopy recordings of embryos, organs, organoids, and non-model species dwarfs anything biologists needed to reckon with in the past (11–13). While microscopy technology has become relatively accessible and ubiquitous, the computational tools to extract the developmental lineages from the data are lagging. Segmenting cells, nuclei or subcellular structures in dense three-dimensional cellular fields and tracking the identified cellular objects over time is a complex image analysis problem, refractory even to modern machine learning techniques (14,15). Moreover, to gain biological insights, the resulting topology of an organismal lineage tree must be enriched with information about the cell’s identities, trajectories, cell cycle length, gene expression, and other cell properties. To realize the full potential of the foundational microscopy datasets, an extensible computational environment is needed to manage complex cell lineage data.

We addressed this challenge by developing Mastodon, a command center for cell lineage reconstruction and analysis from large-scale microscopy image data. Mastodon is built on the solid foundation of its predecessor tools, TrackMate (16) and MaMuT (17). It provides an environment for importing, extracting, curating, visualizing, analyzing, and exporting arbitrarily large cell lineage projects. At its core, it offers functionality to interactively navigate through the analogue microscopy and digital cell lineage domains jointly, at scale, and with high performance. It features open, interoperable, customizable, and extensible software architecture, enabling biologists to import existing lineages and programmers to develop custom lineage extraction and analysis tools. Mastodon bridges to a state-of-the-art visualization platform, Blender (https://www.blender.org/), that facilitates side-by-side comparison of cell lineage datasets in complex 3D geometries over time. At every analysis and visualization level, Mastodon gives users access to the raw microscopy data, providing important context for evaluating the plausibility of the computational lineage analysis hypothesis. Finally, Mastodon is distributed as a plugin for the popular image analysis platform Fiji (18), ensuring ease of installation, long-term stability, and continuous growth.

## Results

### Mastodon aims and features

Research fueled by modern bioimaging requires a robust scientific platform able to visualize, curate, and analyze tracking and lineage data derived from long-term time-lapse imaging. While the requirement is straightforward to articulate, meeting it presents a multitude of technological challenges. Due to the size of the images and tracking datasets, the visualization tools of such a platform need to be interactive and responsive, even when dealing with large data. It must also integrate well with the broader scientific software ecosystem and offer interoperability with the major tools used for cell tracking and associated image file format. It must also be convenient to use and fast to learn, to enable the wide scientific community with a powerful tool. Finally, it must be developed as an extensible platform, which can be reused and quickly augmented by the developer community with new capabilities for cell tracking and analysis. Mastodon is our best effort to address all these challenges. It operates as a command center for tracking (**Fig. 1**), leveraging technologies that we describe below, and that sets Mastodon apart among existing tools (**Extended Table 1**).

**Fig. 1.**
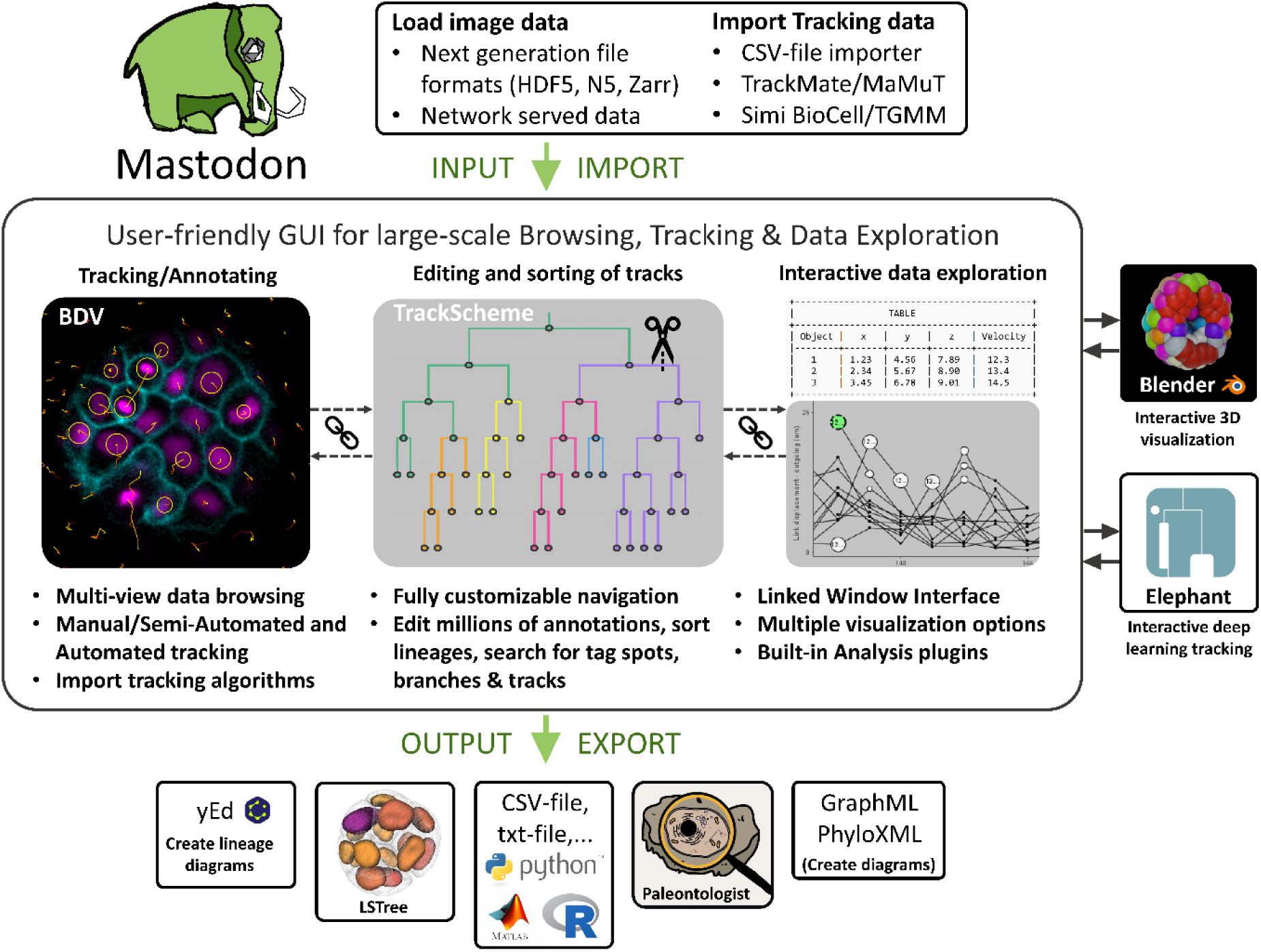
Schematic illustration of Mastodon capabilities. Mastodon is a comprehensive, open-source tracking software designed for managing large-scale tracking data in biological imaging datasets. The software has flexible import and output options. It features an intuitive and interactive GUI with interconnected views, offering a variety of visualisation and analysis tools. Users can generate, edit, and explore tracking data seamlessly across multiple interconnected windows, including BigDataViewer for image data visualisation, TrackScheme view for organising cell lineages, Table view for searching categorical or numerical annotations and Grapher view for plotting them. The software supports automated, semi-automated, and manual tracking methods. For enhanced 3D visualisations, Mastodon interoperates with Blender, providing interactive exploration and high-performance rendering of tracking data. The software’s plugin system allows for custom-made extensions, enabling users to add specific functionalities without modifying the core code (e.g. ELEPHANT).

To deal with very large images, Mastodon uses the BigDataViewer (BDV) technology (19). In addition, to handle large tracking data, we engineered a set of purpose-built libraries that ensure responsive user interfaces and a small memory footprint for very large lineage trees (**Supplementary Information**). These two technologies allow Mastodon to deal with the largest accessible tracking datasets, vastly exceeding predecessor platforms with classical graph data structures TrackMate (16) and MaMuT (17).

The tracking data is represented internally by a high performance data model with two levels of detail. Tracks are stored in a simple directed graph, where the vertices are objects - or ‘spots’ - at a specific time-point, and edges are ‘links’ that connect two spots across time. The spot stores the position, time and shape (encoded as 3D ellipsoids) of the object it represents (usually cells or nuclei). The links are oriented, following the arrow of time, and an event like a cell division amounts to two links emerging from the parent spot, each connecting to one of the two spots representing daughter cells. This data model can handle gaps in tracks (connecting two spots over missed detections) and fusion or division events with more than two spots. Additionally, this core graph can be collapsed to a lower level of granularity where a vertex represents what happened in the lineage between two consecutive cell divisions (**Supplementary Information**). This enables convenient inspection of the hierarchy of cell divisions across lineages as well as computation and storage of numerical features of cell cycles (**Fig. 1**). The data model can additionally store arbitrary categorical and numerical data, associated with spots and links.

From the user perspective, Mastodon takes the shape of a feature-complete, user-friendly scientific software. It offers several views of the tracking data. The main view - called BDV view - overlays the tracking data over the image. The TrackScheme view allows for visualizing tracking data in a hierarchical lineage display. The tracks are laid in columns, discarding spatial information. The spots in a track are arranged vertically in time and division events are represented by a fork in the view. Finally, the Table and Grapher views are designed to display analysis results in tabular form and as configurable 2D plots respectively. Importantly, all the views can be synchronized. The user can trigger a navigation event on an object in one view (for example by clicking on a nucleus in the image and creating a spot), which will bring all the synchronized views to focus on the object. A highlight and focus indicator provides visual cues in all views of the object that is under the mouse or has the keyboard focus. When the tracking data is manually edited, the resulting changes in the views are animated, which facilitates orientation in dense tracking data.

To fulfill the mission of lineage comprehension, Mastodon offers an interactive framework for tracking data analysis. Feature analyzers can be defined for any tracking data types (e.g. spots, links, branches). The default analyzers implemented in Mastodon allow computing the mean intensity in an object, a cell cycle curation, the instantaneous velocity, local cell density, etc. Additionally, the spots and links can be associated with one to many user-defined tags that can represent user- or computationally-defined information such as names and fates of cells.

Importantly, we built Mastodon with several features to help users navigate inside large data. The numerical features described above can be used to decorate the views. Color schemes can be customized based on a feature value and used in any of the views. This helps discover patterns in the tracking data. Finally, we introduce the notion of view context. The TrackScheme or Grapher view can be made to display only the portion of tracking data currently visible in a BDV view. This approach enables the examination of lineage details specific to a segment of the sample in isolation.

Mastodon is interoperable and can be used with hierarchical file formats suitable for large images (e.g., HDF5 (19) , N5 (<u>GitHub - saalfeldlab/n5: Not HDF5</u>), Zarr (20)), stored either locally or remotely. The tracking data can be imported from or exported to several file formats (CSV, TrackMate(16), MaMuT (17), Simi BioCell, TGMM (21), GraphML (22)) selected to cover a wide range of use cases.

Tracking or tracking curation can be performed manually thanks to convenient and fast tools to create, edit and delete spots and link spots (**online documentation, part A**). Manual annotation can be further accelerated by semi-automated tracking, where a configurable algorithm tracks a selected cell forward or backward in time. Mastodon also implements several established algorithms for fully automated tracking (23–25) that can be used across the entire image or within a specific subsection. The manual, semi-automated and automated tracking approaches can be used interchangeably (**Supplementary Video 1**). One of the distinguishing features of Mastodon is that almost everything is customizable. For instance, users can change the key bindings of various operations and the look and feel of the views. This customizability allows users to optimise the tedious and time-consuming task of creating and curating large-scale lineage datasets.

Extracting lineages from multidimensional images is a complex task, subject to active research and development. To maximize the benefit to the scientific community, we have made Mastodon extensible, allowing it to serve as a foundation for further innovations and developments by others. Any of the features described above can be added as Mastodon plugins: new views, new analyzers, new tracking and segmentation algorithms, new input / output filters, new miscellaneous components, etc. This open and extensible software architecture enables Mastodon to serve as an integrated environment for the development of impactful bioimage analysis tools. For example, using the Appose Java–Python communication framework (**see online documentation, part D**), we implemented detectors based on StarDist, Cellpose3, and Cellpose4 and a linker combining Trackastra (26). In addition, ELEPHANT enables incremental deep-learning–based tracking within Mastodon (27).

In this paper, we demonstrate Mastodon’s flexibility by introducing a new view that relies on the powerful open-source visualization and rendering engine, Blender. This new view unlocks Blender’s extensive 3D visualization capabilities and high-performance rendering capabilities for cell tracking in biology. Several instances of Mastodon, showing lineages of different specimens, can be linked to a single Blender scene and facilitate comparison between lineages.

Taken together, Mastodon is an integrated software environment focused on biological lineage data extraction and analysis. It is customizable, easy-to-use, interoperable and extensible. Mastodon is not tailored to any specific biological problem, but rather designed as a flexible toolbox for lineage extraction and analysis in various contexts. Its reliance on modern software libraries tailored for big image data make it performant on large scale lineage datasets. In the next section, we establish the limits of Mastodon using simulated tracking data.

### Benchmarking Mastodon performance on synthetic and biological data

To test the limits of Mastodon’s scalability, we developed a *Simulator* framework (see **Online Methods** for details) that generates lineage trees of arbitrary size. We generated three types of simulations: (i) cubic volumes of simulated spots with regular lineage topology (**Fig. 2a and Extended Data Video 1**), (ii) “gastrulation-like” simulations where growth is constrained by external boundaries arriving at embryo-like appearance (**Fig. 2b and Extended Data Video 2**), and (iii) “spherical cell populations” respecting basic cell volume exclusion rules (**Fig. 2c and Extended Data Video 3**). In all cases, the lineage can be made highly regular, with an equal number of time points between cell divisions, or the cell cycle length can be increasingly random, generating more realistic-looking cell lineages (**Fig. 2d,d’,d’’**).

**Fig. 2.**
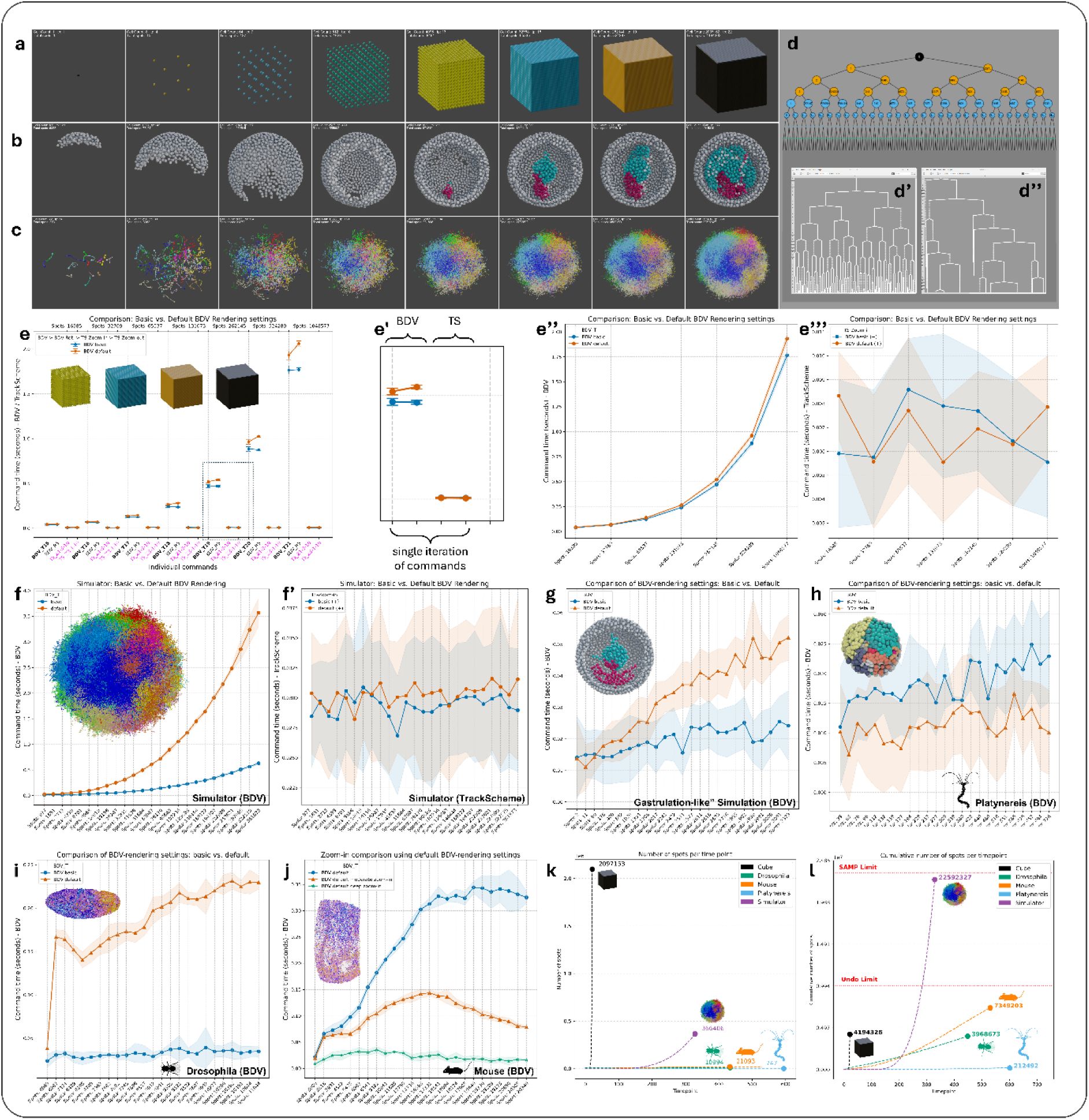
Benchmarking Mastodon performance on synthetic and real biological annotation trees. **(a)** Cubic volume simulation. Cells grow in an entirely regular, cubic arrangement, dividing synchronously to produce an error-free and uniformly expanding cell population over time. **(b)** Gastrulation-like simulation. Cells expand within a confined volume, generating an embryo-shaped aggregate that mimics early developmental patterns. **(c)** ‘Spherical ball’ simulation. Cells divide within a spherical boundary, respecting volume exclusion rules, creating a growing sphere over successive time points. **(d)** Highly regular vs. randomized cell cycles. Synthetic lineages can be generated with strict, synchronized division times (highly regular, see **d** and **d’**), or with varying cell-cycle lengths, resulting in more realistic and heterogeneous trees (randomized, compare **d’’**). **(e)** Performance testing and tasks. Plots display standardized interactions carried out in Mastodon to measure responsiveness during data visualization and data interaction. **(e’)** Close-up of a single iteration of input commands illustrating the performance of BigDataViewer and TrackScheme windows. The command sequence includes loading a new frame containing 262,145 spots (command = BDV_T), and then rotating all spots 360° in five frames (command = BDV_R5). Also shown are measurements from the TrackScheme window, where the responsiveness of a 10-frame zoom-in command (TS_Z1-3-10) and a corresponding 10-frame zoom-out command (TS_Z3-1-10) is evaluated. **(f–j)** Various performance benchmark tests on synthetic and real biological datasets were conducted to evaluate the BigDataViewer command BDV_Tn, which loads new frames containing varying numbers of spots under both basic and default rendering settings. Panels **f′–g′** also report performance metrics for the associated TrackScheme window. **(k–l)** Comparison of artificial annotation data (Cube and Simulator) with real biological datasets (*Platynereis*, *Drosophila*, and Mouse). **(k)** shows the maximum number of spots per time point, dominated by the Cube dataset with over 2 million spots, that can still be visualized in 3d using our Mastodon-Blender bridge whereas **(l)** presents the total cumulative number of spots per time point, with the Simulator dataset showing the highest counts. The first red dotted line shows Mastodon’s limit on continuously adding spots in a single session (for example, when using the simulator), which stems from an effectively infinite number of undo operations (the Undo limit). The second red dotted line indicates the maximum number of spots Mastodon can handle overall (23,342,213 spots), determined by its single-array memory pool (SAMP) approach to memory management.

We used the *Simulator* to generate large lineage tree files that emulate real biological cell lineages. These lineage tree files were loaded to Mastodon and used to measure the responsiveness of the framework. For quantitative responsiveness measurements, we developed a dedicated plugin that records the exact timing between predefined steps of Mastodon GUI interactions. For the benchmark, we defined a standardized series of steps that Mastodon can execute as commands. These include moving to a bookmark position (BDV_T*n*), rotating the dataset in BDV over *n* frames (BDV_R*n*), and performing zoom-in and zoom-out operations in TrackScheme across a range defined by bookmarks (TS_Z), performed in *n* steps (details provided in the Supplementary Information). The benchmark was applied to the cubic volume dataset of exponentially growing size (**Fig. 2e-e’’’**) using a consumer-grade notebook (**ThinkPad X1 Carbon Gen 9**), which served as our reference system (see **Supplementary Information** for details). As expected, drawing operations on cubic volumes with a larger number of annotations took longer to perform in BDV (**Fig. 2e,e’**). The differences between durations of individual steps on the same-sized cubic volumes were negligible (the inclusion of extra spots into the field of view causes a slight increase for rotations). Therefore, as the representative benchmark for BDV and TrackScheme view, we plot the “move to position” and “zoom-in” timing commands, respectively (**Fig 2e’’,e’’’**). For comprehensive plots showing all commands across all benchmarked datasets, see **Extended Data Figures 1-12**.

The performance of Mastodon depends heavily on the amount of details displayed for each annotation. Therefore, to benchmark the BDV view, we compared two options: (i) *basic,* displaying only annotation spot centers, and (ii) *default,* displaying full ellipsoid representations of annotations along with tracks in a user-limited temporal context (see **Online Methods**). We used the “spherical ball” *Simulator* dataset, containing 22.6 million generated annotations, to compare the behaviour of these two display modes. For the default rendering settings, we observed approximately 1 sec of latency at around 100.000 annotations in a single time point (**Fig. 2f**), which is acceptable from the user’s perspective. At 361,000 annotations, the latency increases to more than 3.5 seconds, which compromises the viewing experience. By contrast, under basic rendering settings, performance remained well below one second of latency across the entire dataset (**Fig. 2f**). The largest currently available biological lineage dataset of mouse embryonic development (28) has a maximum of 21.093 cells in one time-point, which is way below the threshold where Mastodon starts to lag. Moreover, from the perspective of a user tracking cells in such data, it makes little sense to display all annotations simultaneously. The user rather zooms in to focus on a fraction of the data. **Fig. 2j** demonstrates on the mouse embryo lineage dataset that zooming into the BDV view significantly improves performance, even under the default rendering settings. This indicates that the drawing performance of Mastodon can be optimized by adjusting display settings and the zoom level.

For the TrackScheme view, we recorded constant performance at roughly 300 milliseconds per frame across the entire spherical ball *Simulator* dataset, indicating that lag is absent in the TrackScheme view (**Fig. 2f’**). We also established in an independent benchmarking experiment that the performance is not affected significantly by the regularity of a lineage tree (**Extended Data Figure 13-15**). Interacting with the TrackScheme window while a BDV window is open results in the same lagging behaviour as the BDV itself. This occurs because the BDV window must continuously redraw whenever the user selects annotations. This lag can be eliminated by either closing all open BDV windows or zooming into them. This shows that the BDV window is the primary performance bottleneck in Mastodon.

The behaviour, where basic settings outperform default ones, was reproduced with a more realistic, gastrulation-like simulated dataset. Since this dataset was significantly smaller, the delays are much shorter (**Fig. 2g**). For a real biological tracking dataset of *Platynereis dumerilii* and *Drosophila melanogaster* (**Fig 2h,i**), there is very little difference in behavior between basic and default settings, except for the first time point of *Drosophila*. This is because, while in *Platynereis*, the number of spot annotations and tracks increases monotonously from 39 initial annotations, the *Drosophila* lineage starts at 8000 annotations, leading to a delay caused by loading many tracks in the default setting for the first time (since a link connects two spots, no links are present at frame 0). This indicates that drawing tracks is a significant factor influencing performance in BDV. However, it does not compromise interactivity for biologically relevant lineage datasets.

We next used the Simulator to explore potential limitations of Mastodon beyond the data size typically used in research (**Fig. 2k**). The system became unresponsive shortly before reaching 10.000.000 spots in a single Mastodon session (**Fig. 2l**). We found that this is due to the “undo” function of Mastodon, which enables the rolling back of an arbitrarily large number of previous operations. The second hard limitation was encountered at 23.342.213 annotations. We traced this showstopper to the array data structure (SingleArrayMemoryPool) used by Mastodon (**Fig. 2l**). We resolved it by switching to an alternative data structure (MultiArrayMemoryPool), currently available in an experimental version of Mastodon. With those roadblocks cleared, it will be possible to work with projects containing more than 150.000.000 annotation spots, although this requires workstation computers with a large amount of memory (120GB and more).

Finally, we asked whether we can use the capabilities of the Blender view to create high-quality renderings of very large datasets. Our tests with the “cubic volume” dataset (**Fig. 2a**) indicate that 130.000 spots can be interacted with, although it takes tens of minutes to pass the necessary data between the two running programs (*Blender-view linked to Mastodon* **Supplementary Information**). Once loaded, the synchronization latency between Blender and Mastodon is about 2 seconds. Additionally, more than 2 million spots (**Fig. 2a**) can be displayed at one time when Blender is detached from Mastodon, i.e., no interactivity with Mastodon (*Blender-view advanced visuals* **Supplementary Information)**. When dealing with time lapse data, even datasets containing cumulative 22 million spots can be imported into Blender for rendering (**Fig. 2c**).

In summary, the Mastodon lineage analysis command center infrastructure scales far beyond the most extensive lineage datasets available today (**Fig. 2k,l**) and thus represents a future-proof computational solution for lineage analysis.

### Comparison of Mastodon performance and functionality with other software platforms

To understand what is behind Mastodon’s scalability, we compared the performance of Mastodon with its ancestor, MaMuT (17). The software architectures of these two frameworks are highly comparable, analogous to closely related species in a phylogenetic tree. However, the graph data structures in MaMuT rely on a general graph library, JGraphT (29), while Mastodon is based on a custom library that we developed specifically to address the challenges of tracking cells in large samples. We argue that such a performance comparison is fairer than contrasting platforms built in different programming languages, which would be like comparing different phyla. To compare the performance of MaMuT, we used the same artificial dataset of cubic volumes of vertices with an entirely regular lineage topology employed in our Mastodon benchmark tests (**Fig. 2a**). Our results show that while MaMuT can still handle more than 65.000 spots aggregated in a single time point (time point 17 of the cubic volumes), with a noticeable response time of under one second, it is unable to handle around 130.000 objects (timepoint 18 of the cubic volumes) in a fluid manner. The primary bottleneck in MaMuT does not lie in the MaMuT viewer (which is similar to the BDV view of Mastodon), but rather in the TrackScheme view. TrackScheme not only takes a significant amount of time to open, but also exhibits substantial lag, making it practically unusable for manipulating annotations or tracking. Notably, MaMuT requires approximately 5 times more memory to store a lineage than Mastodon, further contributing to its performance limitations (**Extended Data Table 1**). Mastodon’s TrackScheme overcomes these limitations due to a novel data structure we developed to store, analyze, edit, and display tracking data efficiently (see **Online Methods** and **online documentation, part E**). In short, Mastodon introduces a memory-local graph model, an allocation-free API for responsive editing, adaptive level-of-detail for scalable lineage visualization, and fast kD-tree-based object retrieval for efficient 3D rendering. These innovations are implemented in modular Java libraries designed for extensibility and reuse. Our results illustrate that creating tools able to harness large image data requires the development of specialised, low level, optimized data structures.

While quantitative comparison across platforms is inherently problematic, we can qualitatively compare the features of Mastodon to those of other lineaging software platforms. There is a vast landscape of available software tools for cell tracking, each offering different solutions for tracking, track editing, analysis, and visualization of generated tracking data, and sometimes even the simulation of trajectories. Some tools focus heavily on the tracking algorithm and do not provide user-friendly graphical interfaces (GUIs) or visualizations, often lacking the capability to edit linked objects. However, they might offer segmentation and deep-learning-based pipelines for tracking cells. Some tools focus entirely on visualization and analysis. Several different tools have integrated Mastodon as an integral part of tracking and track-editing within their visualization and analysis pipeline (30–32). Mastodon’s strength lies in its ability to handle large-scale data while covering most tracking solutions in a user-friendly way. **Extended Data Table 2** provides an overview of various tracking software’s unique features and capabilities.

Thus, we propose Mastodon as an excellent compromise between performance, accessibility, interoperability, and extensibility for lineage tracing.

Using Mastodon to extract, visualize, and analyze lineages in various contexts In the following, we illustrate Mastodon’s capabilities through three specific use cases that address almost all the challenges biologists face when analyzing large time-lapse datasets.

### De novo tracking and visualization

The most common use of Mastodon is the creation, curation, examination, and visualization of cell lineages in newly acquired microscopy datasets. We demonstrate this in the embryonic development of the marine annelid *Platynereis dumerilii,* a key model system for the abundant spiralian species (33). *Platynereis* exemplifies the mosaic mode of embryonic development, where different embryos exhibit highly similar, topologically invariant lineages. We obtained two recordings of *Platynereis* embryonic development transitioning a) from a 38-cell stage spiral cleaving embryo to a 754-cell early larva over 15 hours and b) from a 39-cell stage to a 1270-cell early larva over 15.5 hours. These two *in toto* light sheet microscopy datasets comprised 601 and 620 time points captured as 3D image stacks. The first embryo was tracked by running an automated tracking solution TGMM (21) and partially correcting the errors using a modified version of CATMAID (34) and, in part, using Mastodon. This approach yielded a lineage tree of 212,468 tracked nuclei. The computationally established initial lineage tree was fragmented into multiple sub-lineages. All these had to be examined and connected to the main tree, which had to be manually reviewed to correct errors. For the second embryo, we opted for a semi-automated tracking using Mastodon, proceeding as follows. The raw image was opened in Mastodon. Next, we performed semi-automatic tracking, following single cells across time points until the subsequent cell division. We then quickly reviewed and validated the new track, used Mastodon lineage editing tools to establish a cell division, and repeated the process until a complete lineage of 212,490 tracked nuclei. The advantage of the semi-automatic Mastodon approach is that it limits compounding errors. The annotator corrects tracking mistakes along the way, taking advantage of multiple interconnected BDV views that show the raw data from multiple viewing angles (see **Fig. 3a,a’,a’’, Supplementary Video 1**). These views are linked to the growing lineage in the TrackScheme view (**Fig. 3b**, right). Supervised semi-automated lineage extraction from raw data of this magnitude represents the most accurate deployment of the Mastodon lineaging infrastructure. One shared limitation of both semi-automated Mastodon and curated TGMM tracking is the lack of journaling of the tracking process. Therefore we cannot report how many clicking operations inside the respective user interface had to be performed to arrive at the final curated lineage. To better quantify the effort necessary for lineage reconstruction, we ran again the automated TGMM tracking (21) on the first embryo and compared it against the ground truth generated through manual curation (**Fig. 3**) using the TRA (tracking accuracy) measure (35,36). The result showed that correcting the TGMM result to the ground-truth state required 162,893 operations. This corresponds to a normalized TRA of 0.6, indicating that fewer than 40% of the operations needed for full reconstruction were required to curate the automated output. For the semi-automated Mastodon tracking, the number of operations can be estimated from the number of cell divisions, branches, and cells in the lineage. The embryo one has 731 divisions, 1504 branches and 212,468 cell detections in 601 time-points. Thus the least amount of operations needed to establish the lineage semi-automatically is 731 assuming the process only fails at cell division. If the user has to click on every single cell detection, the lineage tree will be built in 212,468 steps. From the experience of tracing the lineage in embryo two, we estimate that the number of operations needed is biased towards the lower bound. Thus, compared with the 162,893 clicks required to fix the TGMM result, the Mastodon process appears for this dataset less laborious.

**Figure 3.**
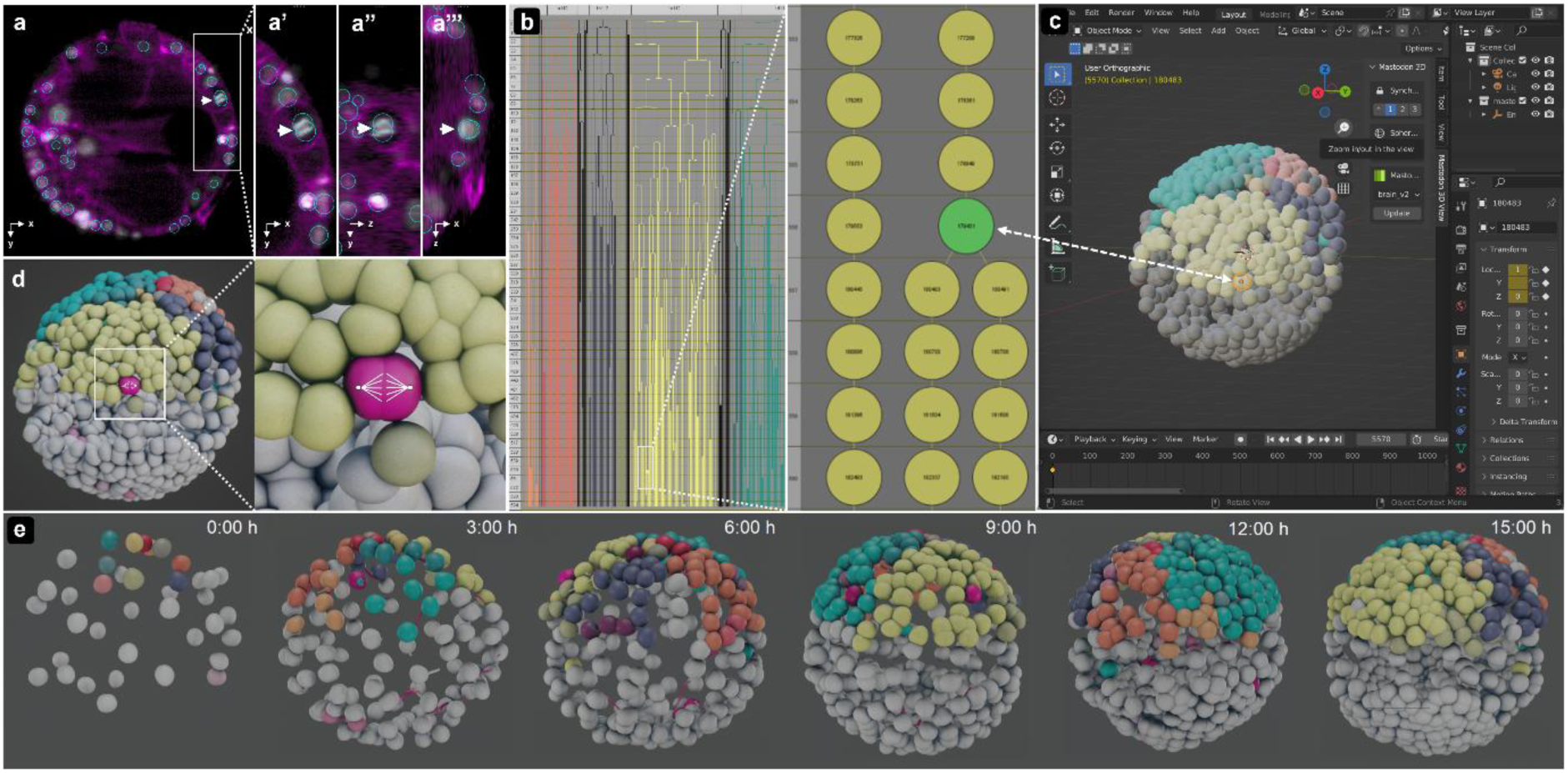
The power of integrating diverse visualization approaches through automatic synchronization. **(a)** Three synchronized BigDataViewer (BDV) windows display the same dataset from different perspectives. Each window shows the raw image data along with overlaid annotations, such as tracked nuclei highlighted by cyan circles. A dividing cell is marked with a white arrow. BDV windows **(a’-a’’’)** Zoomed-in BDV views from orthogonal angles: **(a’)** xy-plane, **(a’’)** yz-plane, and xz-plane **(a’’’)**, provide spatial context and improved inspection of raw data and annotations **(b)** The TrackScheme window functions as an interactive lineage browser and editor synchronised with the BDV windows in (a). Each tracked object, e.g. a nucleus, is visualized as a spot (vertex) that can carry metadata, such as labels and user-defined tags in various colors. Tracks are visualized as horizontally arranged spots, with time progressing vertically to connect them. **(c)** A 3D rendering of the tracking annotations is visualized in an interactive Blender window. Selecting a spot in Blender automatically highlights the corresponding object in TrackScheme (b), all BDV views (a) and vice versa. Blender can also retrieve and display the color-coded information from Mastodon. **(d-e)** 3D visualizations using the ‘Advanced Blender view’. Cell divisions are automatically highlighted in purple. In (d), a dividing cell is further annotated manually with a schematic representation of the mitotic spindle. **(e)** Overview of the embryo development of an annelid worm over 15 hours of in toto live imaging, capturing the transition from a 38-cell early cleavage stage to a 754-cell early larva.

Once the lineage tree is established, Mastodon provides powerful visualization and analysis tools. For instance, we manually labeled all descendants of brain progenitor cells using Mastodon’s tag feature to mark the fates of several specific cell lineages contributing to brain development (37). These tags, when selected, are immediately visible within the TrackScheme and BDV views in their assigned colors and can be passed to the 3D renderings of spheres in Blender (**Fig. 3b,c,d,e** and **Extended Data Video 4**). This greatly facilitates the separation of the lineages and examination of the spatio-temporal evolution of the lineage during development. Beyond lineages, one can establish a tag set to highlight, for example, cell divisions (**Fig. 3d** and **Extended Data Video 5**) or other features of interest in the data. The ‘Interactive Blender Window’ (**Fig. 3c**) allows for selecting any individual spot at any time point within Blender, and this spot will then be automatically synced and highlighted, e.g., within the TrackScheme/BDV windows.

In this use case, we demonstrated how Mastodon’s semi-automated tracking, flexible and interconnected GUI, and interactive visualization tools allow for comprehensive mapping and exploration of cell lineages.

### Comparison of multiple lineages

Any meaningful analysis of cell lineage trees requires comparing lineage trees of different specimens to each other. Even for stereotypically developing species, this is a challenging task. To compare lineages, the trees have to be registered. Only then can the annotations on the tree, represented, for example, by the color tags in Mastodon, be transferred from one manually annotated specimen to the next and compared in context. The major obstacle to lineage tree registration is that any node in the lineage tree can be flipped. As a result, even if the trees and their branch topologies appear identical, the actual spatial relationships between cells may not be. To ensure that sister branches do not differ between multiple datasets, it is crucial to determine the spatial arrangements of all daughter cells and their new branches after each cell division (**Supplementary Information**). To address this challenge, we used the Mastodon plugin architecture to implement a *Spatial Track Matching* tool (**online documentation, part C**). The plugin requires that the datasets to be registered have minimal morphological variance, root nodes are consistently labeled, and a minimum of three initial cell divisions are matched manually.

To demonstrate the functionality, we imported ten cell lineages of the ascidian species *Phallusia mammillata* (Pm), which had already been meticulously annotated using the ASTEC pipeline (38) (**Fig. 4a** and **Extended Data Video 6**). During the import (see online methods), we converted the cell fate annotations from the original paper into Mastodon tags. Embryo Astec-Pm-10 was subsequently rendered in various ways (fates, tracks, divisions, and left-right bilateral symmetry) using the Mastodon-Blender bridge, showcasing the powerful new visualization capabilities we have established (see **Extended Data Video 7**). To test the registration plugin, we first removed the tags from one 64-cell stage embryo (**Fig. 4b**), except for its first time point. After registration with a reference embryo (Pm-08) with tags (**Fig. 4b**, **Extended Data Video 8**), the tag set information and labels were transferred to the recipient embryo. The procedure was repeated across all ten embryos, with embryo Pm-10 as the reference (**Extended Data Video 9**). We then used Mastodon to explore and compare any two embryos in 3D. This was greatly facilitated by the plugin automatically linking and highlighting corresponding cells across the two embryos (**Fig. 4c** and **Extended Data Video 10**). Doing this, we identified occasional branch incongruences among the lineages based on differences in cell division directions, underscoring the need to re-examine computationally registered lineage data and the power of Mastodon to reveal such problems. In the original Guignard et al. study, the authors demonstrated significant similarity in the cell lineages of homologous cells, quantified using a normalized tree-edit distance. We implemented the tree edit distance metric in the *Hierarchical Clustering* plugin (**online documentation, part C**) for Mastodon and applied it to the registered, clean lineages. The resulting clusters revealed the relationship between cell lineage architecture and cell fates to be equivalent to the Guignard et al. study (**Fig. 4d**).

**Fig. 4.**
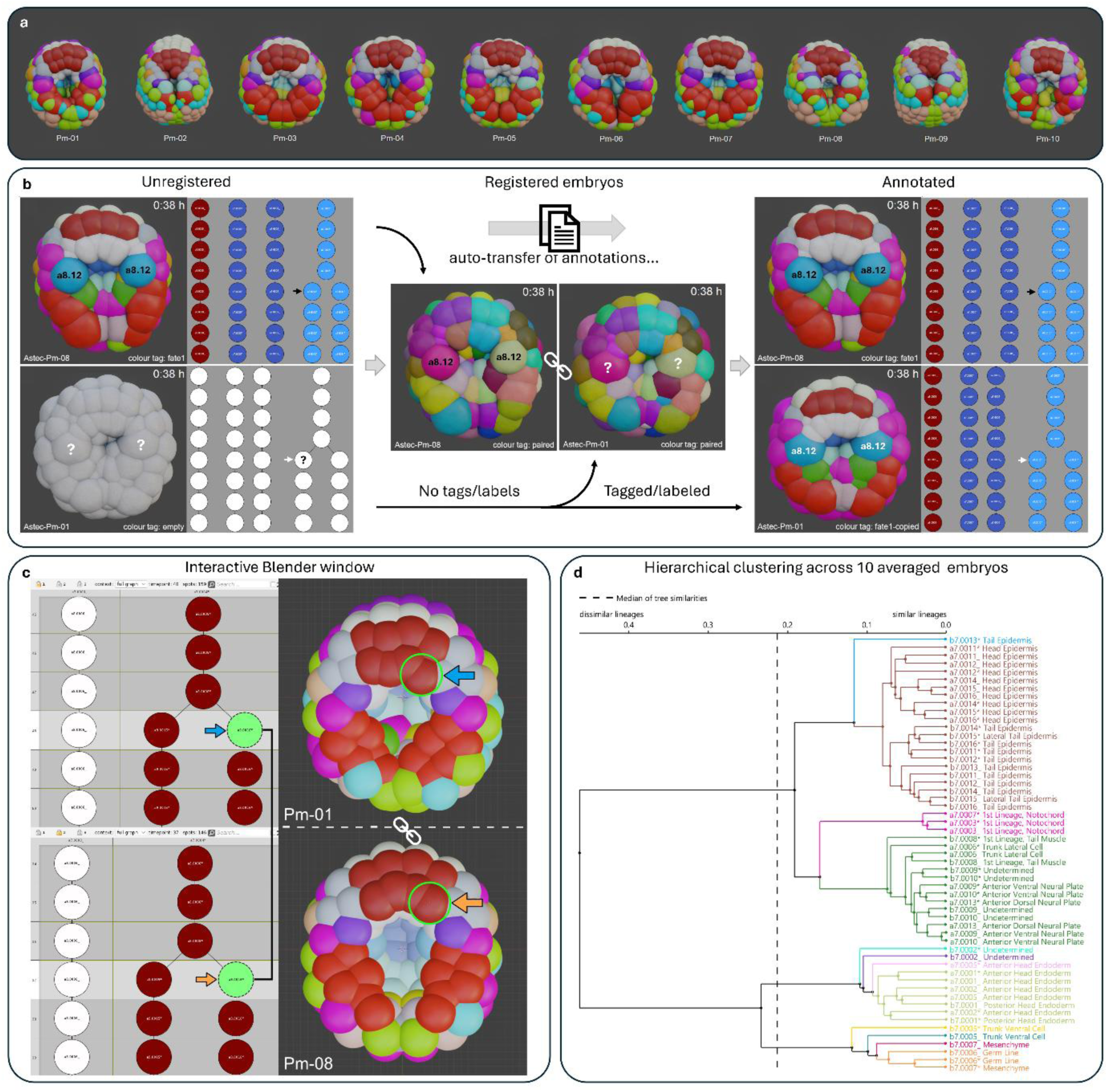
Working with multiple embryos inside Mastodon with invariant developmental patterns. **(a)** Ten digitised *P. mammillata* embryos were imported and 3D-rendered using the Mastodon-Blender bridge. Displayed from the vegetal view with the anterior at the top, these embryos are shown 2 hours and 2 minutes after the start of imaging, colour-coded according to Guignard et al. (2020). **(b)** Two selected embryos, Pm01 and Pm08, demonstrate the capabilities of the ‘Lineage Registration’ tool. In this hypothetical scenario, all annotations in Pm01 were completely erased and then automatically recovered using embryo Pm08, including colour tags and labelled spot annotations. Blastomere a8.12 is highlighted as an example, but the entire cell lineage tree’s annotations are recovered as long as the donor embryo provides corresponding information. **(c)** Illustration of the automatic selection feature for two registered embryos (Pm01 and Pm08). When a cell is selected in one embryo, either as a spot in the TrackScheme Window or as a 3D sphere in the Blender view, the corresponding spots in the sister embryo are automatically highlighted. This is indicated by the blue and orange arrows. **(d)** A dendrogram resulting from the hierarchical clustering of individual cells across multiple cell lineages of averaged embryos was created using Mastodon’s ‘Hierarchical Clustering of Lineage Trees’ plugin.

This use case demonstrates Mastodon’s flexibility in implementing registration and analysis routines for lineage trees. Mastodon’s integrated environment enables simultaneous comparison of tree topologies for multiple specimens and their quantitative analysis.

### Lineage exploration in Mastodon

In the previous use cases, we have shown the power of Mastodon in extracting, visualizing, and analyzing well-behaved, curated, and stereotypic cell lineages. Unfortunately, for reasons both technical (imperfect segmentation and tracking) and biological (regulative non-stereotyped development), most lineage data are fragmented and incomplete. For the visualization and analysis of such noisy lineages, Mastodon provides capabilities to extract numerical features from tracks and visualize them on the raw data annotations, the lineage trees, and create 3D renderings with the Blender view.

We demonstrate this on four diverse datasets (lymphocyte movement simulation, spheroid development, Drosophila and mouse embryogenesis), using six different numerical features (branch average movement, link velocity, movement speed relative to nearest neighbours, branch sinuosity, branch duration and displacement, spot intensities (see **Extended Data Video 11**) and cell divisions and by using them to visually tag objects in Blender (**Fig. 5a-d, Extended Data Video 12-16**).

**Fig. 5.**
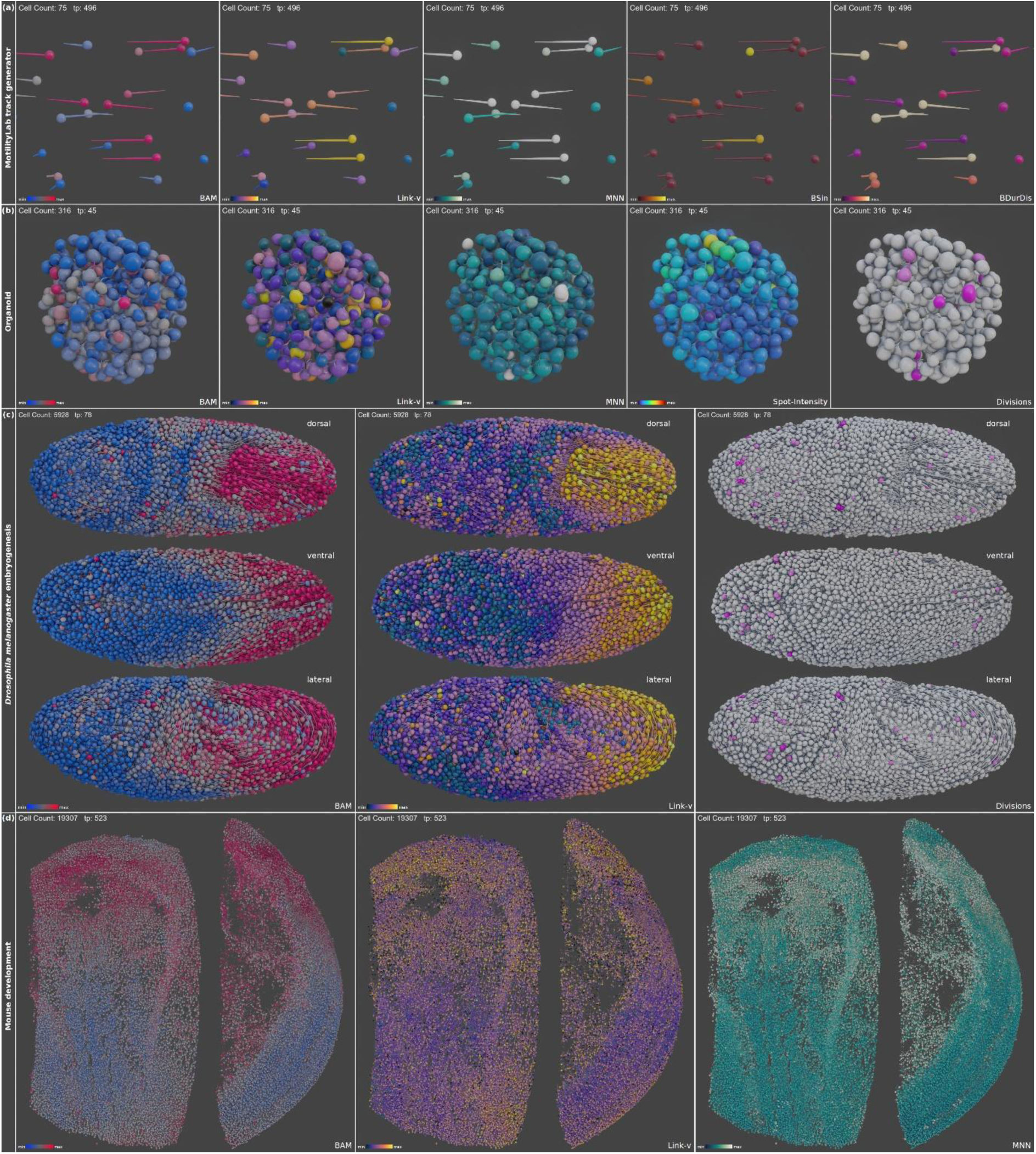
Feature-based coloring of simulated and biological lineage datasets with diverse developmental patterns and cell identities that cannot be mapped across embryos or specimens. **(a)** Generated tracks of simulated lymphocytes using MotilityLab displaying stop-and-go motion exemplify how various cell behaviour can be visualised after applying colour-tags to various computed feature values. Branch average movement (BAM) and Link-velocity (Link-v) highlight slower and faster moving cells, a feature also indicated by longer and smaller track ‘tails’. Visualising the average movement speed of cells relative to their five nearest neighbours (MNN) in this simulation also allows distinguishing between fast and slow-moving cells. Note that the simulated lymphocytes in Fig. 4a are only a cut-out of a larger field of view. The full field of view is separately available as Supplementary Video 13. The displayed cell count values refer to the full simulation. **(b)** A spheroid tracked semi-automatically over 143 time points by Carlos O. Gomez from the lab of Prof. Otger Campàs at TU Dresden, Germany. During the tracking period, the number of cells in the spheroid increased from 292 to 371. In addition, spot intensity features were extracted from the image data and added into the visualisation. (c) *Drosophila melanogaster* embryogenesis tracking data () visualised in Branch average movement (BAM) and Link-velocity (Link-v) **(d)** Mouse data visualised in Branch average movement BAM, Link-v and MNN.

We envision this framework being used in an exploratory manner. For example, a biologist may find a group of cells moving through the embryo in a particular direction. They can then examine those cells in the raw data, isolate them within the large and chaotic lineage, and collect summary statistics on their collective behaviors using Mastodon. This exploration can also be initiated from the other end. Starting with a search in the feature table, passing through the data to the 3D developmental context. Or they can start from the expert knowledge of the lineage and examine the features and developmental context of specific branches of the lineage tree.

Taken together, Mastodon allows biologists to explore complex tracking data and to extract meaningful information from it.

## Discussion

The lack of tools that can analyze the dynamics of single cells in large samples and long movies creates a blind-spot in Life Science, where phenomena at these scales evade researchers. Mastodon alleviates this bottleneck and enables scientists with an interactive tool that can create, visualize, analyze and curate cell lineages at scales previously unattainable. Importantly, Mastodon is designed as a user-friendly tool, widely usable by the scientific community, trivial to install thanks to its deployment within the Fiji ecosystem and fast to learn. Mastodon is interactive: it allows for the exploration and annotation of image data over time. Thanks to the novel libraries we developed, its user interface remains responsive even when dealing with millions of objects. Its combination of scalability, usability, and interactivity positions Mastodon as a transformative resource for quantitative biology, democratizing access to single-cell dynamics at scales previously out of reach of conventional tools. We also envision that Mastodon will have an indirect impact on the development of novel tracking algorithms. Currently, the development of AI-based tracking algorithms is hindered by the lack of tracking ground truth datasets (36). Such datasets are movies that have been manually tracked and curated by experts. Generating them represents a long and formidable task. Mastodon is a tool built with many facilities that considerably alleviate its difficulty and will greatly accelerate the generation of AI-ready tracking datasets.

Mastodon is implemented in pure Java, leveraging the BDV (19) framework to enable seamless visualization of large-scale 3D image datasets. To enhance compatibility with modern data formats, we extended BDV’s capabilities to support N5 (https://github.com/saalfeldlab/n5) and OME-Zarr (20) storage backends, including remote access to datasets hosted on the cloud. While Mastodon’s Java-based implementation ensures seamless integration with the Fiji ecosystem, innovation in algorithms for cell detection, segmentation, and tracking - such as StarDist (39), cellpose (40,41), Trackastra (26), and ultrack (42) - are predominantly developed in Python. Currently, Mastodon interfaces with these tools via import / export filters, a workflow that will benefit from emerging standardized formats like GEFF (https://github.com/live-image-tracking-tools/geff). However, achieving scalable, end-to-end cell tracking in large datasets will require direct integration of Python algorithms into Mastodon, necessitating both adaptation for high-performance computing and cross-language interoperability. Existing approaches, such as TrackMate’s subprocess-based execution (16), rely on file-based data exchange, but newer technologies like the JDLL library (43) and its appose extension (https://github.com/apposed/appose) enable low-latency, in-memory communication between Java and Python. Future developments in Mastodon will leverage these advancements to enable real-time interaction with Python-based tools, eliminating intermediate file handling and enhancing workflow efficiency for large-scale single-cell analysis.

This work introduces a framework for quantifying user interface responsiveness in scientific software when handling large datasets. Comparative benchmarks between Mastodon and its predecessor, MaMuT, demonstrate that Mastodon’s optimized data structures and visualization algorithms reduce memory requirements, enabling stable and interactive performance in scenarios where MaMuT fails. To maximize accessibility, we implemented Mastodon’s core visualization components in pure Java without hardware acceleration, allowing seamless operation on standard hardware or via remote connections. We anticipate that future extensions will introduce GPU accelerated views, further enhancing responsiveness. Since we developed Mastodon as a modular and extensible platform, these improvements can be developed independently as plugins, requiring minimal modifications to the core framework and avoiding the need for a complete redesign.

In Mastodon, cells are modeled as ellipsoids. They are well adapted to a vast amount of use cases in developmental and stem cell biology, where large samples are imaged at intermediate magnifications. However, as imaging technologies advance toward higher resolution enabling more complex morphological analysis, Mastodon must evolve to accommodate arbitrary object shapes. This presents a substantial technical challenge: Mastodon’s current data structures rely on fixed-length records (e.g., ellipsoids) to ensure computational efficiency and responsiveness. In contrast, polygons (2D) and meshes (3D) can represent complex shapes, but require variable-length records that complicate memory management and performance optimization. Future extensions to Mastodon will therefore require novel data structures that preserve the scalability and interactivity demonstrated in this work while supporting flexible shape representations.

Mastodon establishes a robust and future-proof platform for cell lineage reconstruction, curation, and analysis in developmental biology. Its scalability, extensibility, and ease of integration with existing tools make it ideally positioned to become a central hub for lineage-centric research workflows. Looking ahead, we envision several promising directions for Mastodon’s development and deployment. First, the plugin-based architecture enables rapid integration of emerging technologies in cell tracking, including deep learning-based segmentation, probabilistic tracking, and real-time feedback systems for live imaging experiments. As lineage datasets continue to grow in size and complexity, Mastodon’s performance-optimized core and modular design will allow it to adapt to increasing computational demands. Second, Mastodon’s interoperability with visualization platforms like Blender opens avenues for immersive exploration, 3D lineage storytelling, and integration with virtual or augmented reality environments for education and outreach. Third, we anticipate broader adoption of Mastodon beyond developmental biology, including applications in cancer evolution, regenerative medicine, and organoid modelling — domains where spatiotemporal tracking of cells is equally critical. Fourth, Mastodon will fuel the development of novel tracking algorithms by facilitating and accelerating the creation of tracking ground truth datasets. Finally, the community-driven nature of Mastodon ensures that it can evolve organically. By empowering developers and users to contribute custom modules, analysis routines, and data connectors, Mastodon can serve as a living, extensible ecosystem. As imaging technologies continue to advance and cross-disciplinary demands grow, Mastodon is well-positioned to bridge the gap between image data, computational lineage analysis, and biological discovery.

## Online Methods

### Core design and implementation of the Mastodon framework

Mastodon was developed with the primary objective of handling very large numbers of objects on standard computers, while ensuring a highly responsive user interface for editing and exploring these large datasets. The innovations that are at the core of Mastodon include (i) a new graph data model exploiting memory locality to improve the speed of iteration, (ii) an allocation-free Application Programming Interface (API) that improves the responsiveness of the user interface when editing tracking data, (iii) adaptive level-of-detail in the lineage display, enabling the interactive exploration of large lineages, and (iv) efficient object retrieval with kD-tree search in convex polytopes, to improve the speed of rendering and querying the 3D tracks. The resulting implementation is organized in several Java libraries, which are part of the Mastodon framework. We took care to make the code extensible, allowing researchers to easily build on Mastodon’s functionality and reuse its libraries in other domains, including - but not limited to - cell tracking and life sciences. The innovations mentioned here are described in detail in the **Supplementary Information Part 1: Mastodon core technologies**.

### Integration of Mastodon Data into Blender for 3D Visualization

We developed a pipeline to import Mastodon tracking data into Blender for high-quality 3D visualization. Spots are converted into vertices, and links into mesh edges, with spot attributes (e.g., radius, color, time point) stored alongside. Using Blender’s Geometry Nodes, we procedurally generate spheroids for spots and tapered tubes for tracks. This node-based system allows real-time customization of parameters such as size, resolution, and visual style. Two export modes are supported: 1) Blender View (Linked to Mastodon): Enables interactive 3D visualization synchronized with Mastodon’s BDV and TrackScheme via spot selection in Mastodon or sphere selection in Blender. Time interpolation and tag-based coloring are supported. This mode is ideal for exploration and inspection of smaller datasets. 2) Blender View (Advanced Visuals): Exports the entire mastodon project as a single object for performance-optimized rendering of millions of spots. Geometry Nodes presets visualize tracking data as spheroids and tracks, with adjustable modifiers for appearance and time windowing. To support high-quality rendering and publication-ready output, we provide a customizable .blend template (default_empty.blend) that enables advanced features such as surface displacement and ambient occlusion, and is fully compatible with Blender’s Cycles engine for point cloud rendering of massive-scale datasets. Additional implementation details, modifier controls, and usage guidelines are provided in the **Supplementary Information Part 2: Integration with Blender via the Mastodon Bridge**.

### Simulated cell lineages in Mastodon

We developed two distinct methods for generating large-scale synthetic datasets in Mastodon: agent-based simulations and a cubic-volume lineage generator. The latter generates perfectly regular cubic spot arrangements, ideal for scalable, controlled benchmarking, with the option to link spots into lineages or leave them unconnected. The more complex agent-based simulator models cells as rigid spheres in 3D space where each agent moves according to its behavioral state and local interactions, including stochastic Brownian motion and distance-dependent repulsion to avoid overlaps. Agents follow a simplified cell cycle with configurable division and lifespan dynamics, enabling realistic population growth over time. Lineages are constructed by linking simulated cells across time points using Mastodon’s native spot infrastructure. Crucially, the simulator interfaces directly with Mastodon’s optimized kd-tree–based querying system, allowing efficient neighborhood detection even at scale. It also supports seamless integration with pre-existing data, enabling hybrid simulations and interactive exploration of spatial constraints, phase changes, and dense lineage structures.

For the gastrulation-like simulation, we defined a structured 3D scene using layered spot arrangements acting as barriers and spatial constraints. A single agent was then seeded into this environment and simulated proliferation over hundreds of time points, producing realistic spatiotemporal lineage dynamics using Mastodon’s agent-based simulator, which models independent agents with full spatial awareness and behavioral rules.

Furthermore, the agent-based simulator was used to generate spherical cell populations with over 22 million linked annotations. These synthetic datasets can be generated within minutes on a standard notebook. Implementation details, generation scripts, and parameters are provided in the **Supplementary Information Part 2: Synthetic data generation via Mastodon’s *Simulator***.

### Performance benchmarking

We assessed Mastodon’s rendering performance for lineage annotations using both BDV and TrackScheme views. A custom benchmarking tool, integrated into Fiji, measures wall-clock time between issuing and completing individual rendering requests. To isolate overlay performance, benchmarks were performed on .mastodon projects containing only annotation data (no image pixels). The tool uses a command language to define and execute view-specific rendering tasks sequentially, ensuring reproducibility across platforms. Benchmark runs used predefined TrackScheme bookmarks and consistently opened one BDV and one TrackScheme window per test. Results reflect realistic user experience and help identify performance limits under varying annotation loads. More detailed descriptions of the benchmark setup, the BDV rendering options used, command syntax, and reproducibility measures are provided in the **Supplementary Information Part 2: Performance benchmark for rendering annotations**.

### Data creation benchmark (MaMuT versus Mastodon)

A synthetic dataset was programmatically generated using MaMuT and Mastodon data structures. The dataset starts with 10 cells, each undergoing 17 divisions at intervals of 5 frames. Reported numerical values represent the median of 5 independent runs. Benchmarks were performed on a Mac Studio (2023) with an Apple M2 Max CPU, 64 GB RAM, running macOS 15.6 (aarch64). The Java environment was Java HotSpot™ 64-Bit Server VM (build 25.441-b07), version 1.8.0_441.

### Biological dataset generation/Import

#### Imaging and lineage tracking of Platynereis embryos

Two *Platynereis dumerilii* embryos were imaged from 5 hours post-fertilization (hpf), with imaging intervals of 90 seconds and a z-step size of 2.031 μm. Pixel size was 0.406 μm (6.5 μm camera pixels with 16x objective). Membranes and nuclei were fluorescently labeled in separate channels by microinjection of H2B-eGFP mRNA (185 ng/μl) and CAAX-mCherry mRNA (200 ng/μl). Embryo 1 was imaged from 5 hpf to 44 hpf (starting at the 38-cell stage), and tracked up to time point 600 using TGMM, followed by manual curation in CATMAID. Embryo 2 was imaged from 5 hpf to 24 hpf (starting at 39 cells), and tracked up to approximately time point 600 using Mastodon, which was also used for manual curation. Embryos were then color-tagged in Mastodon, and Embryo 1 was visualized using the Blender View (Advanced Visuals) via our Mastodon-Blender Bridge, as described in **Supplementary Information Part 2: Integration with Blender via the Mastodon Bridge**.

#### Imaging and lineage tracking of Spheroid dataset

The dataset comprising 143 semi-automatically tracked time points was kindly provided by Carlos O. Gomez from the laboratory of Prof. Otger Campàs (TU Dresden, Germany).

The spheroid was generated from 4T1 murine breast cancer cells transfected with miRFP670-H2B. Cells were cultured in hanging drops for 3 days, then transferred onto a 1% low-melting-point agarose pad and allowed to further compact for 1 day. On day 4, the spheroid was imaged using a Zeiss Lightsheet Z.1 microscope equipped with a 20x objective. Time-lapse imaging was performed for 12 h at 5 min intervals. The resulting dataset was tracked by semi-automatic tracking using Mastodon.

### Importing Ascidian lineage datasets into Mastodon

Lineage datasets from *Phallusia mammillata* embryonic development (38) were imported into Mastodon. Original data in ASTEC format were first converted to CSV using published scripts, then imported into Mastodon using a custom ASTEC-to-Mastodon parser. Image data was not included, and only datasets Pm01–Pm10 were converted for use in hierarchical lineage clustering, spatial registration across embryos (spatial track matching), and visualization tasks. More details about the import and the scripts used are available at the following link: https://github.com/mastodon-sc/mastodon-example-data/?tab=readme-ov-file.

#### Importing Mouse and Drosophila datasets

The mouse and *Drosophila* datasets (28) were imported from published tracking files (*mouse_full_tracks_071621.txt* and *drosophila_side_1_tracks_071621.txt*), available at: https://janelia.figshare.com/collections/Whole-embryo_lineage_reconstruction_with_linajea/7371652.

The files were processed using Mastodon’s built-in CSV Importer, which supports spot coordinates, metadata, and lineage links via ID and parent ID columns. For these datasets, we reassigned the cell_id and parent_id values to smaller integers, as the original IDs exceeded Mastodon’s CSV importer limit of 2,147,483,647. The importer is available in Mastodon under File > Import > CSV Importer, with further details provided in the **online documentation, part C: CSV Importer**.

### Analysis of datasets in Mastodon

#### Phallusia mammillata dataset (ASTEC)

The initially randomly assigned color annotations (color-tags) of Ascidian embryos (Pm01–Pm10) were manually adjusted to closely match the ‘Fate1’ colors used in the original publication (Guignard et al., Science 2020). To ensure consistency, all lineages were aligned to start at time point 0. This was done via the command ‘Plugins > Spots management > Spots shuffling (Simulator) > Translate spots in space and time,’ available when the Mastodon-Simulator update site is active in Fiji. Subsequently the ten embryos were sequentially loaded into a single Blender file for visualization (see **Supplementary Information Part 2: Integration with Blender via the Mastodon Bridge - Enhancing visualizations and enabling true rendering**). Mastodon’s Hierarchical Clustering of Lineages tool (**online documentation, part C**: track editing, analysis and export add-ons) was applied across all ten averaged embryos, using Embryo Pm01 as the reference. Clustering was performed with average linkage and the Normalized Zhang Tree Distance on data cropped between 64 and 180 cells, with no minimum division threshold. Defining 12 clusters resulted in 64 lineage trees. The Spatial Track Matching tool (see **online documentation, part C: track editing, analysis, and export add-ons**) was used to copy tag sets, their colors, and spot labels from the reference embryo Pm10 to all other embryos, processing one embryo at a time, as only two Mastodon projects can be registered simultaneously. To achieve this, we used the 64-cell stage (available in all ten datasets) as the set of parent cells, providing a known correspondence based on identical cell labeling. The algorithm then computes cell division direction vectors between daughter cells in both embryos and compares them after spatial alignment. For alignment, we used the dynamic track matching method, which computes transformations (rotation, translation, and scaling) based on the positions of tracked root cells and their descendants over time, allowing the method to compensate for embryo movement, particularly rotation, during development. Finally, the following operations based on the correspondence information were performed: copying tags between corresponding cells in both embryos, and copying cell names.

#### Feature-Based coloring of simulated lymphocytes and biological lineage datasets

For all datasets used in feature-based coloring (Simulated Lymphocytes, Spheroid, *Drosophila*, and Mouse), we first computed all available features (Spot, Link, and BranchSpot) in Mastodon (**online documentation, part A: Numerical features and tags**). Additionally, for the Mouse dataset, we computed the average relative movement at the branch level relative to the five nearest neighbors using a custom tool in Mastodon (Plugins > Compute Features > Movement of spots relative to their nearest neighbors). The resulting features were then adjusted for colormap and min-max range within Mastodon’s Feature Color Mode before being visualized in Blender using the Mastodon-Blender Bridge.

#### References cited in Extended Data Table 2

3DeeCellTracker (2021) (44)

AceTree (2006, 2018)(45,46)

Angler (1997) (47)

ASTEC (2020) (38)

BioEmergences/Mov-IT (2016) (48)

DeepCell Kiosk (2021) (49)

CATMAID, modified (2014) (21)

CeLaVi (50)

CellTrackVis (2023) (51)

Cell-ACDC (2022)(52)

cellPLATO (2024) (53)

CellProfiler (2006, 2021), CP Tracer (2015)

CellTrack (2008) (54)

(The Tracking Tool - tTt (2016) (55)

CellTracker (2021) (56)

CellTrackR (2021) (57)

CellTracksColab (2024) (58)

DeLTA 2.0 (2022) (59)

EmbryoMiner 2018 (60) Icy(61)

Ilastik (2019) (62)

IMARIS (2007) (63)

LEVER 3-D (2011, 2014, 2016) (64–66)

LIM Tracker 2022 (67)

Lineage Mapper (2016) (68)

LSTree (2022) (31)

MaMuT (2018) (17)

Mastodon (this paper)

MorphographX (2014, 2022) (69,70)

MorphoNet (2019, 2025) (71,72)

MotilityLab (57)

Paleontologist (30)

Simi BioCell (3)

TGMM (2014) (21)

TrackMate (2017, 2022) (16,73)

u-track3D (2023) (74)

ultrack (2024) (42)

Volocity (https://www.volocity4d.com/)

Zeiss arrivis Pro (formerly Vision4D) (https://www.zeiss.com/)

## Code availability

https://github.com/mastodon-sc/

## Acknowledgements

We thank Carlos O. Gomez and Prof. Otger Campàs (TU Dresden) for generously allowing us to use their tracking data for visualisation. We are grateful to members of the Tomancak Lab, in particular Bruno Vellutini, for sharing their expertise in image processing. We also thank Florian Jug, Michalis Averof, and Anastasios (Tassos) Pavlopoulos for their support, valuable feedback, and ideas throughout the development process. We gratefully acknowledge the support of the MPI-CBG Light Microscopy Facility in Dresden. Our gratitude further extends to Curtis Rueden, whose maintenance of Fiji keeps the foundation strong on which Mastodon builds. Finally, we thank all test users and the broader Fiji community for their helpful feedback. This work was supported by the ANR–DFG Funding Programme (grant nr. ANR-21-CE13-0044 and grant nr. 490966236) and by core funding from the MPI-CBG.

## Author information

These authors contributed equally: Johannes Girstmair, Tobias Pietzsch Co-corresponding authors: Jean-Yves Tinevez, Pavel Tomancak

## Author Contributions

Conceptualization: PT, JYT, TP

Funding acquisition: PT, JYT, RH

Core framework development: TP, JYT

Development of extensions: SH, MA, VU, JYT, KS, SP, JG

Benchmarking and simulator implementation: VU, JG

Data generation and annotation: MHT, JG, VU, SH

Blender implementation: MA, SP, SH, JG

Validation and evaluation: SH, VU, JG

Visualization / figure preparation: JG

Supervision: PT, JYT, JG, RH

Writing – original draft: JG, PT

Writing – review & editing: JG, PT, JYT, SH, SP, VU, MHT

## Ethics declaration

The other authors declare no competing interests.

## Supplementary information

Refer to SupplementaryInformation.docx

## Source data

The source code of our Mastodon-based cell tracking tool is available at https://github.com/mastodon-sc/.

## Rights and permissions

Mastodon itself is distributed under the BSD 2-Clause License

---

**Extended Data Figure 1.**
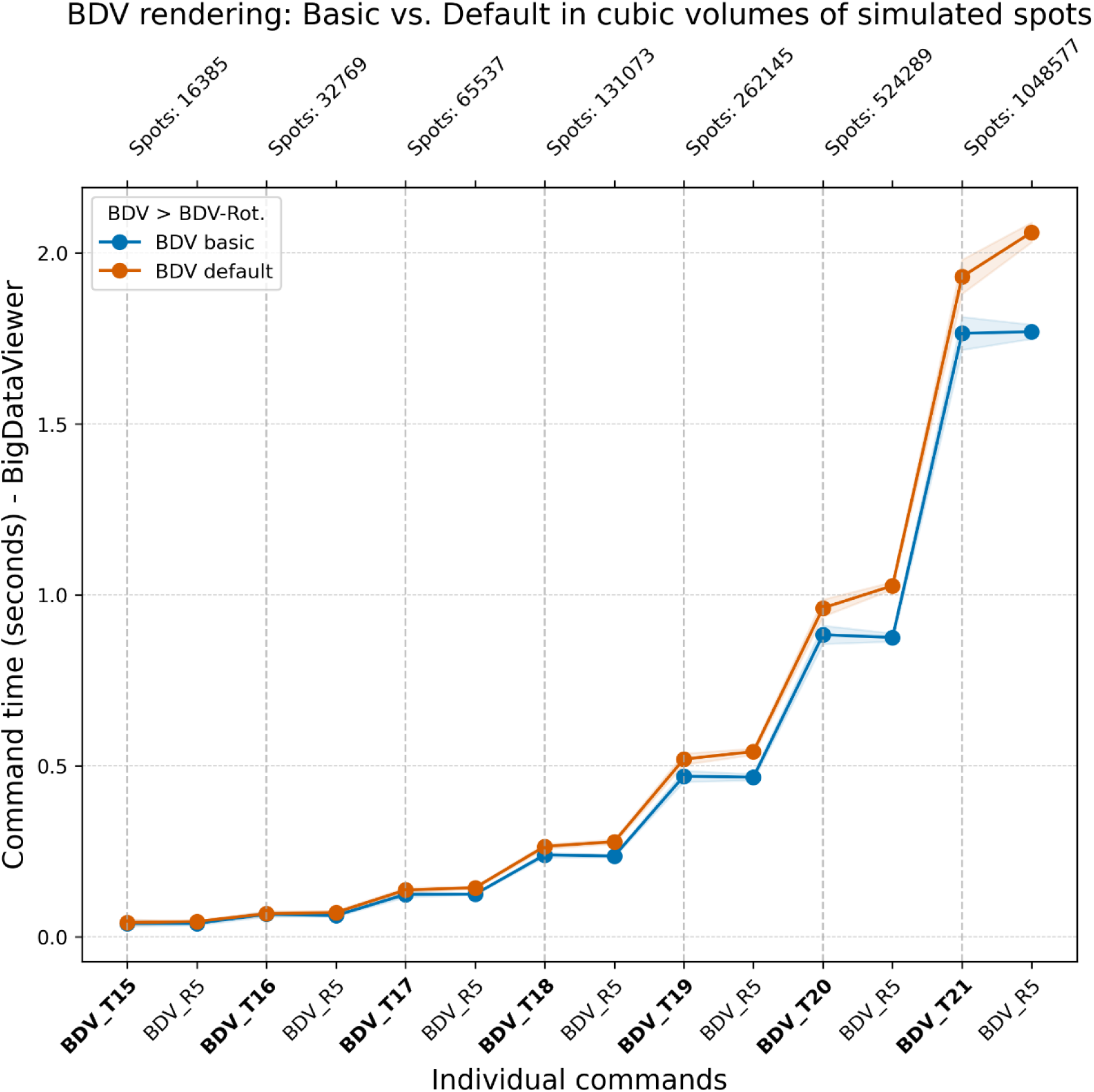
Performance of BDV window commands on cubic volumes of simulated spots, measured as the time (in seconds) required to either render annotations (BDV_Tn, where n is the time point) or perform a 360° rotation (BDV_Rn, where n is the number of frames used for the rotation), using either Basic or Default rendering settings.

**Extended Data Figure 2.**
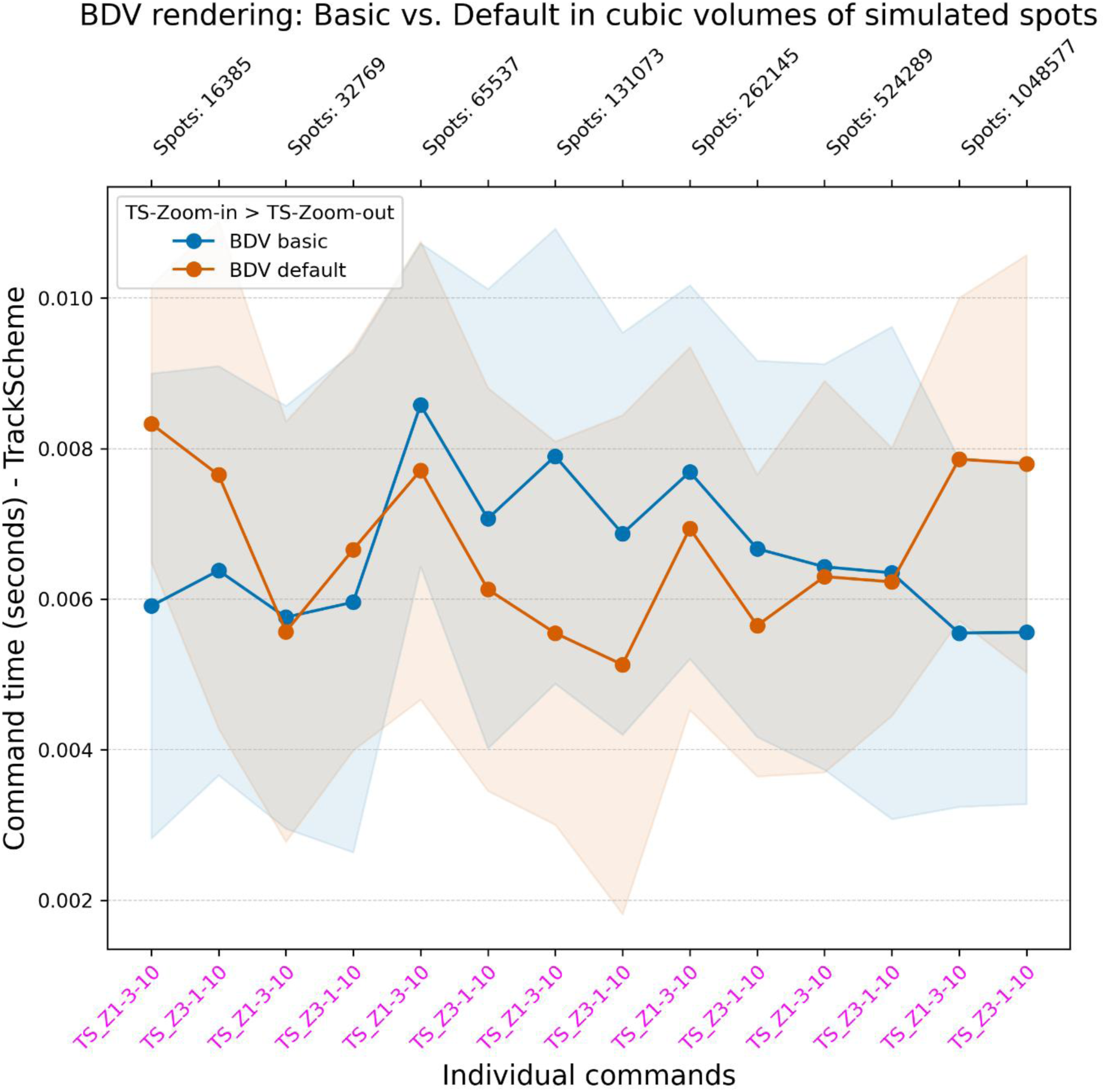
TrackScheme command performance on cubic volumes of simulated spots, measured as the time (in seconds) required to zoom in (TS_Z1-3-10) and then zoom out (TS_Z3-1-10), using 10 interpolation frames per zoom operation.

**Extended Data Figure 3.**
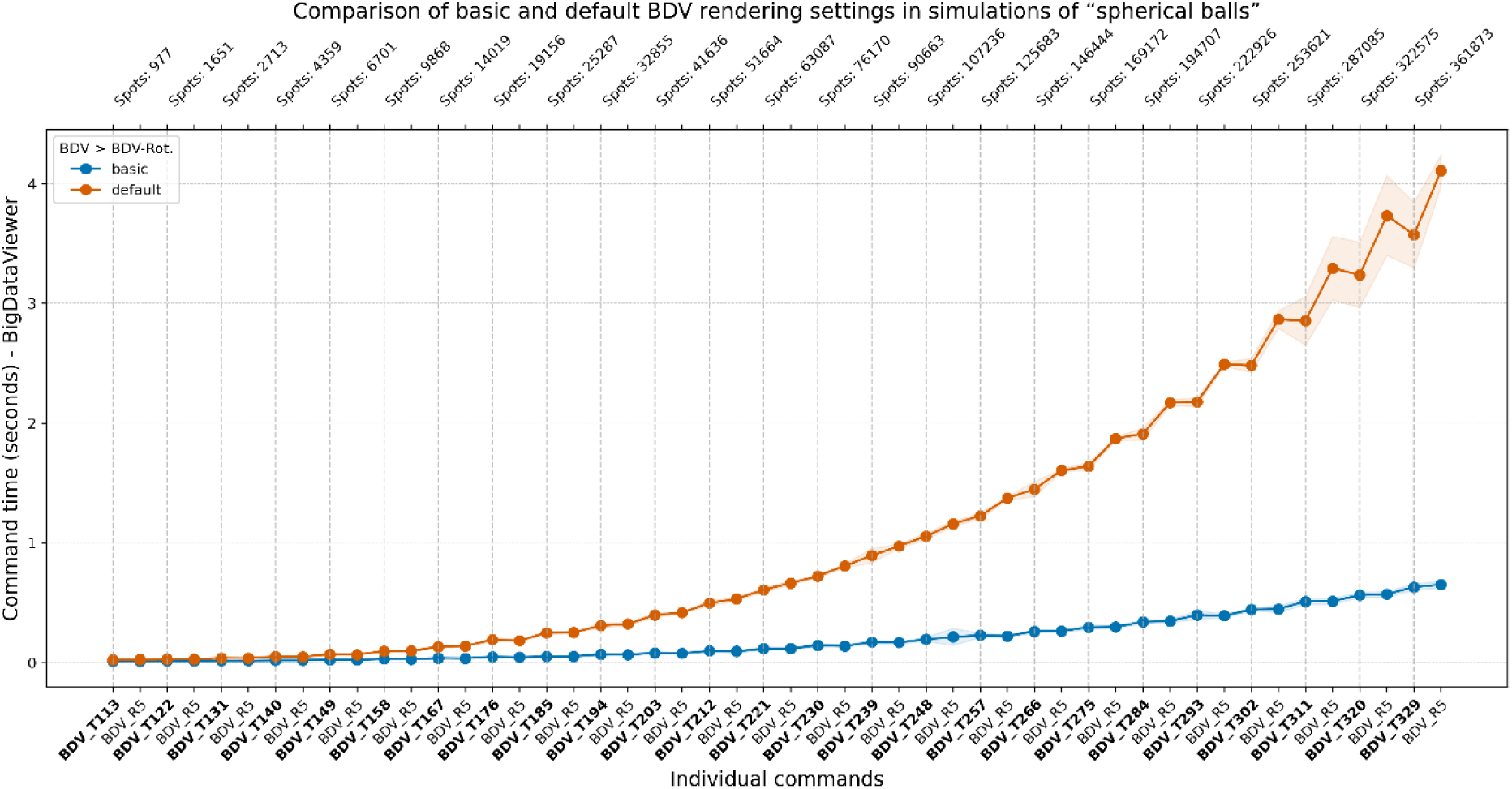
Performance of BDV window commands on a simulation of spherical cell populations, measured as the time (in seconds) required to either render annotations (BDV_T*n*, where *n* is the time point) or perform a 360° rotation (BDV_R*n*, where *n* is the number of frames used for the rotation), using either Basic or Default rendering settings.

**Extended Data Figure 4.**
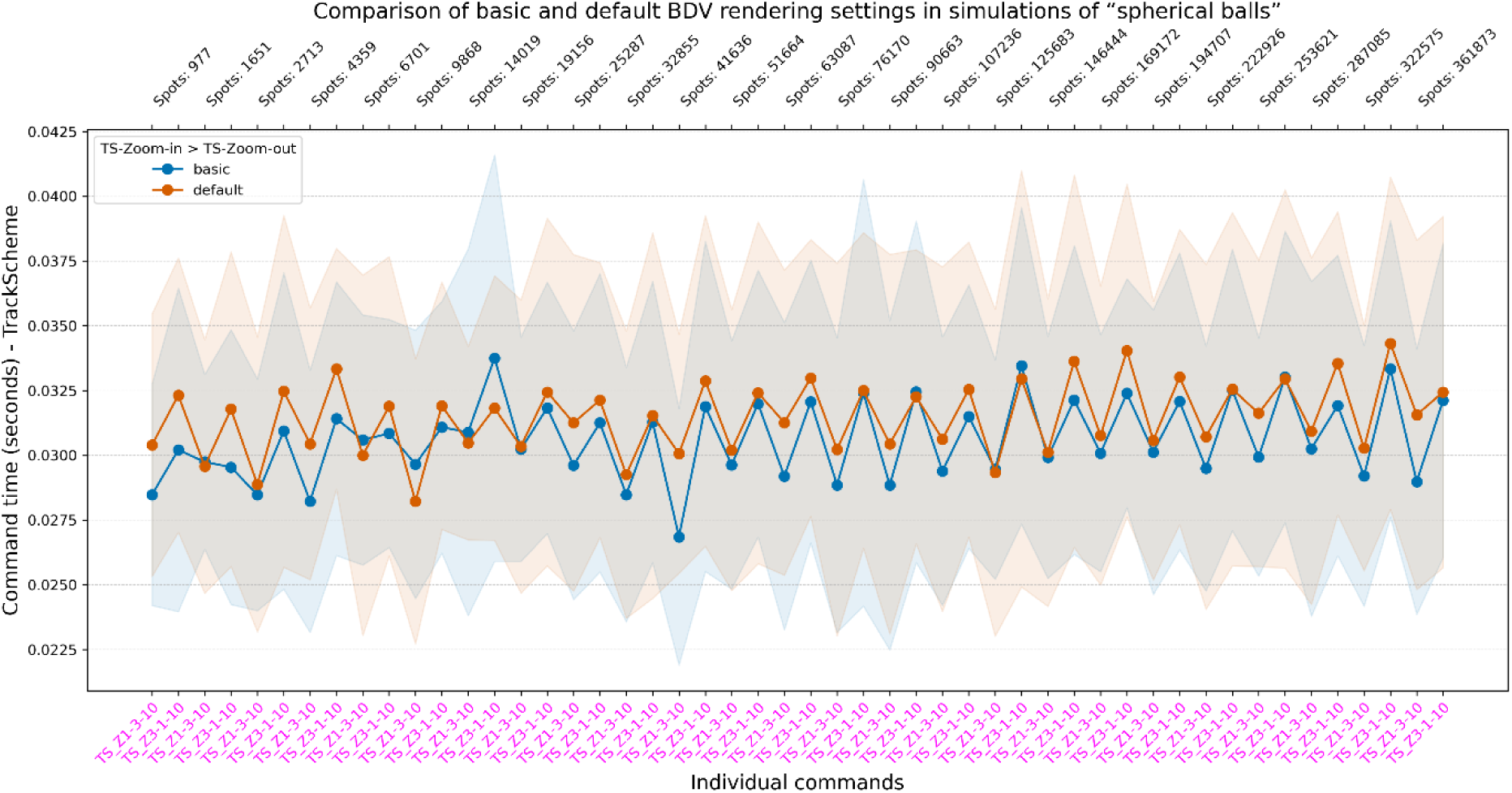
TrackScheme command performance on a simulation of spherical cell populations, measured as the time (in seconds) required to zoom in (TS_Z1-3-10) and then zoom out (TS_Z3-1-10), using 10 interpolation frames per zoom operation.

**Extended Data Figure 5.**
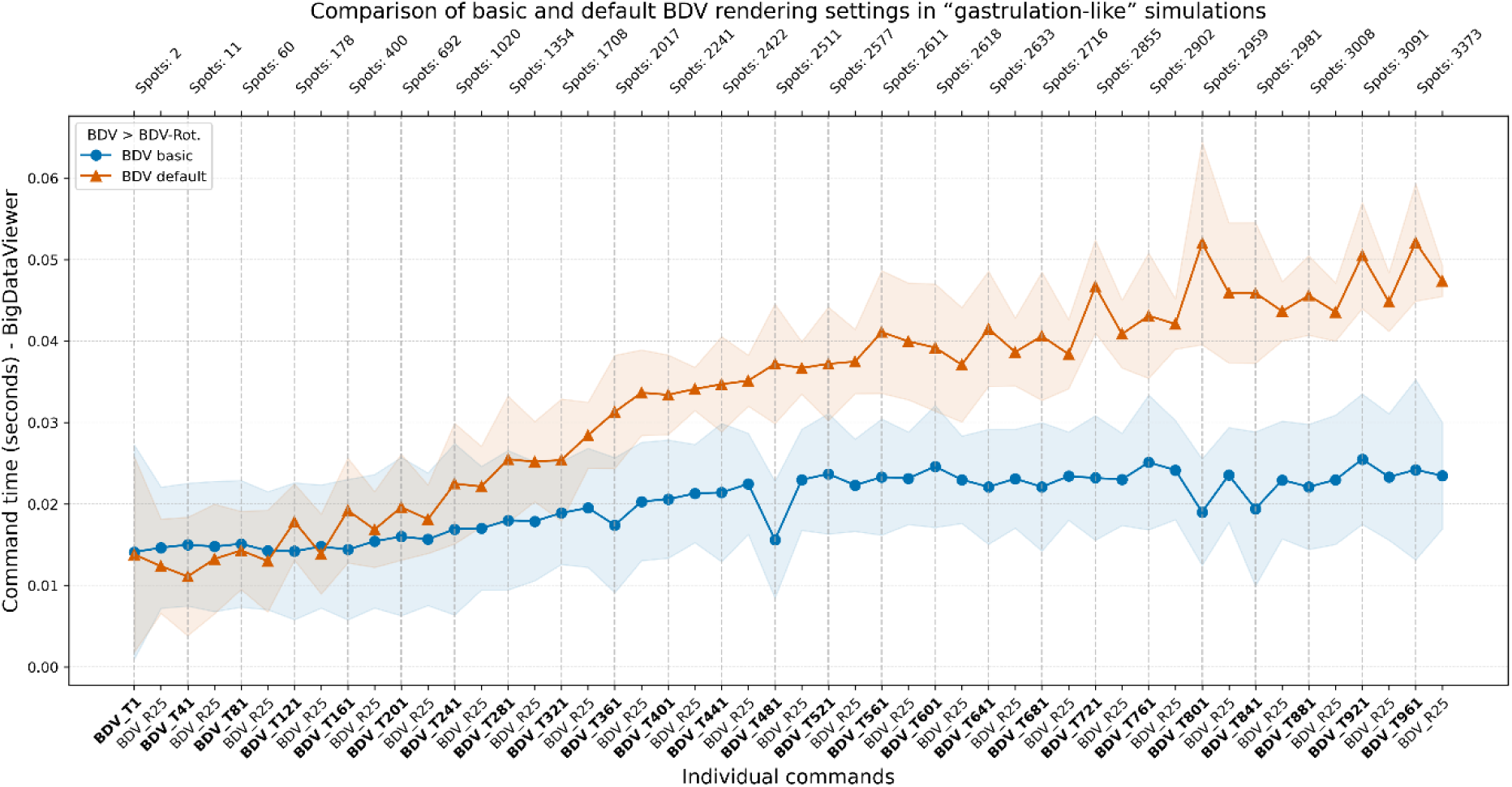
Performance of BDV window commands on a ‘gastrulation-like’ simulation, measured as the time (in seconds) required to either render annotations (BDV_T*n*, where *n* is the time point) or perform a 360° rotation (BDV_R*n*, where *n* is the number of frames used for the rotation), using either Basic or Default rendering settings.

**Extended Data Figure 6.**
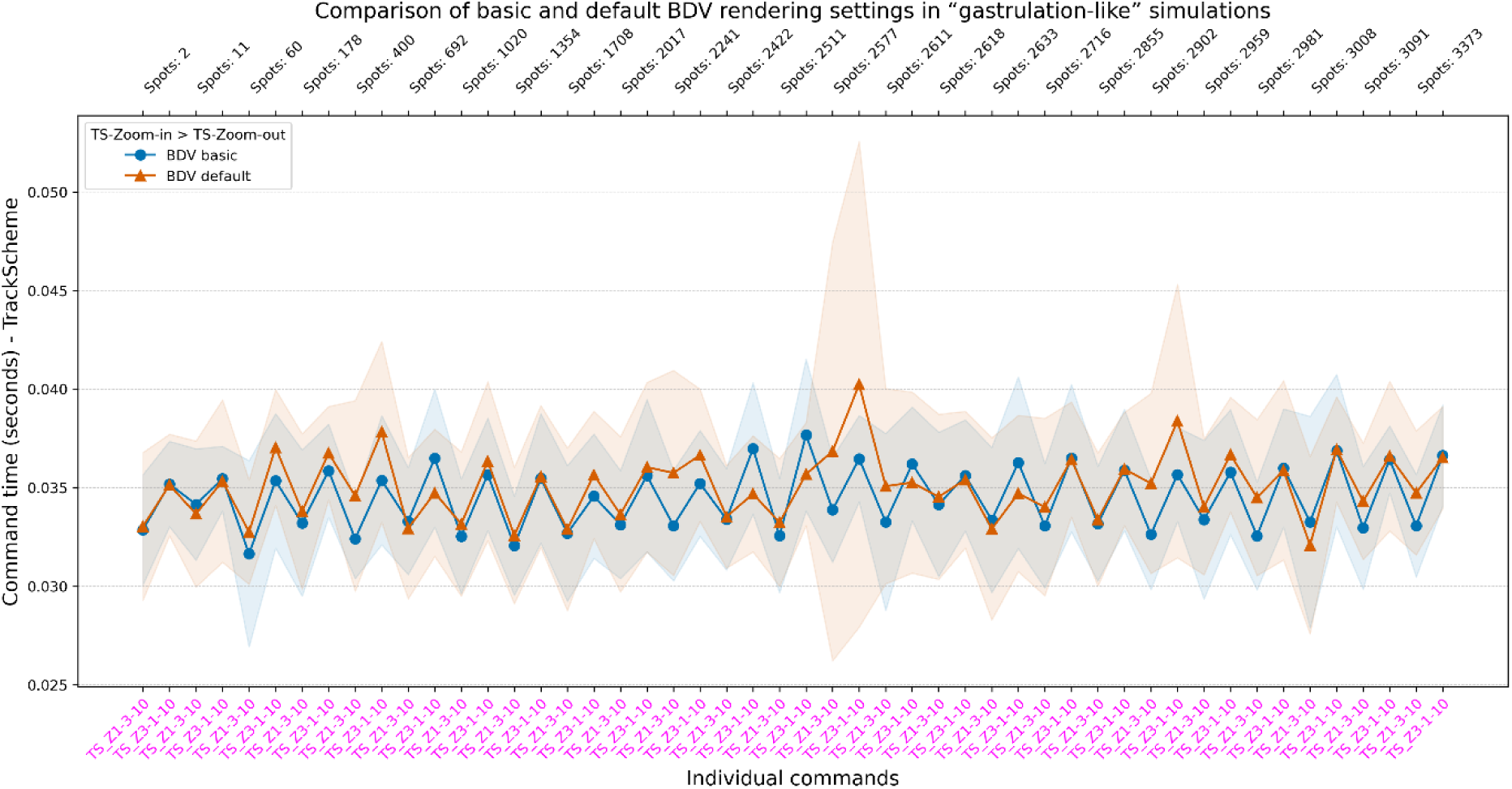
TrackScheme command performance on a ‘gastrulation-like’ simulation, measured as the time (in seconds) required to zoom in (TS_Z1-3-10) and then zoom out (TS_Z3-1-10), using 10 interpolation frames per zoom operation.

**Extended Data Figure 7.**
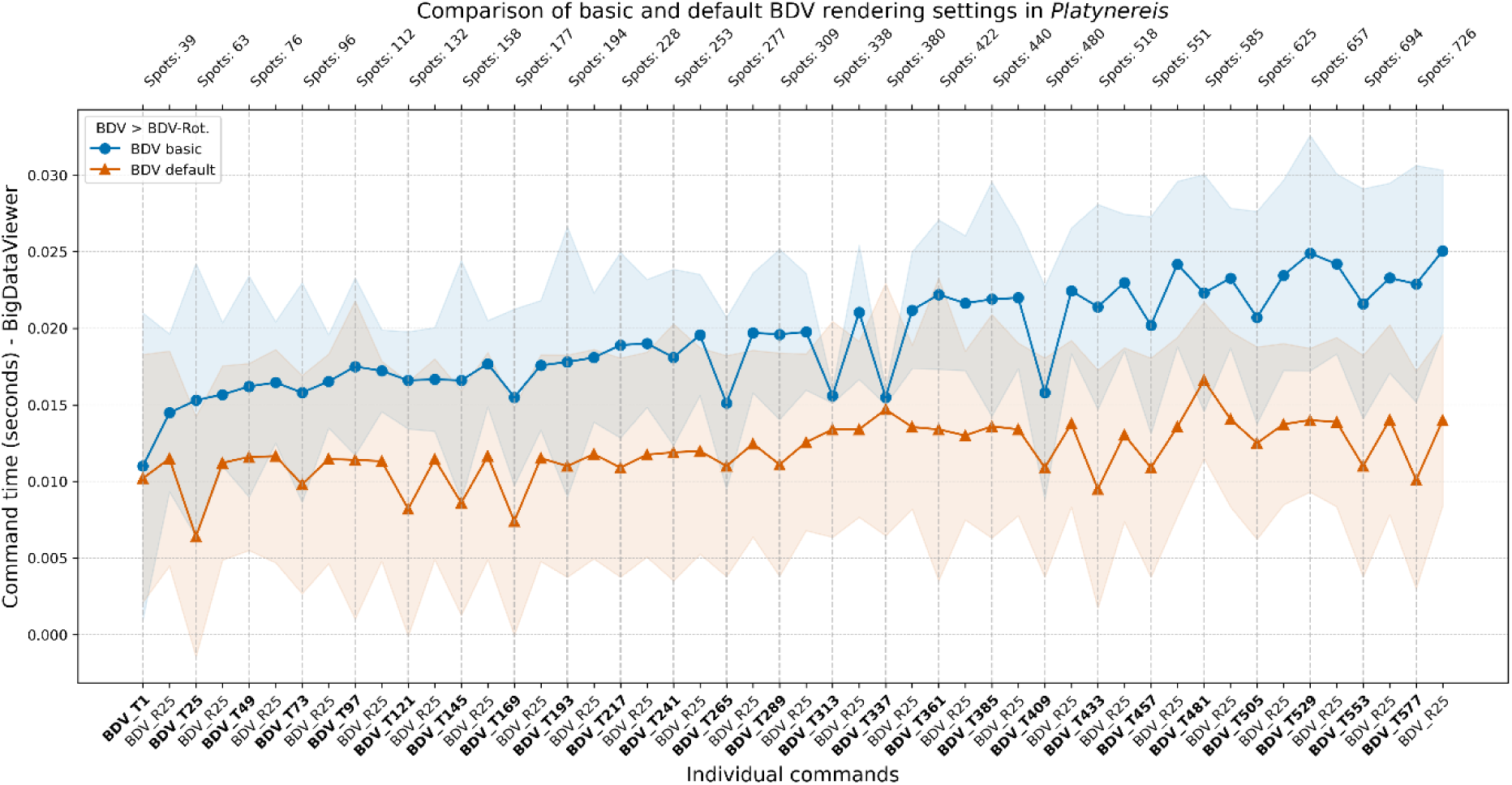
Performance of BDV window commands on a *Platynereis* dataset, measured as the time (in seconds) required to either render annotations (BDV_T*n*, where *n* is the time point) or perform a 360° rotation (BDV_R*n*, where *n* is the number of frames used for the rotation), using either Basic or Default rendering settings.

**Extended Data Figure 8.**
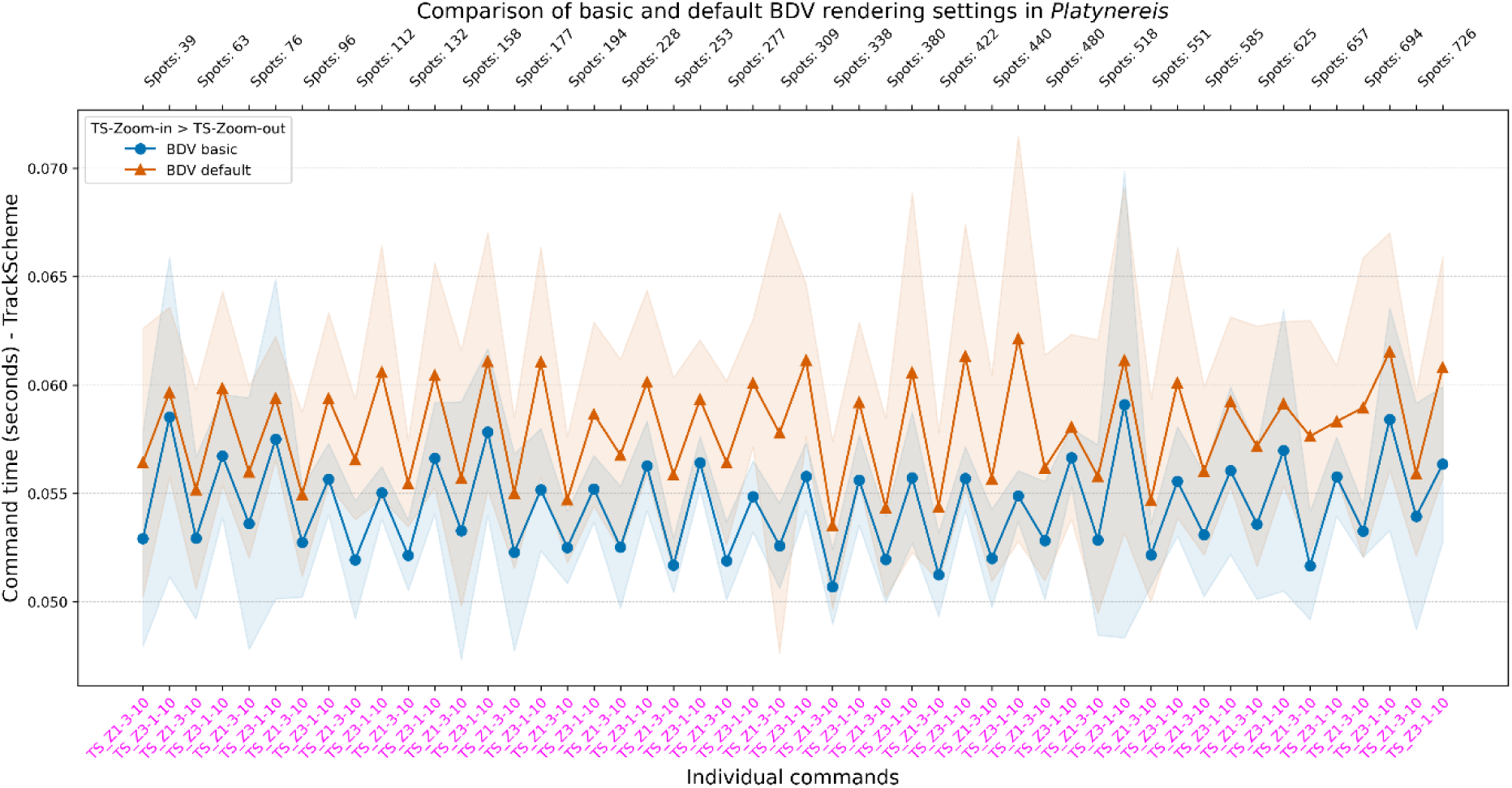
TrackScheme command performance on a *Platynereis* dataset, measured as the time (in seconds) required to zoom in (TS_Z1-3-10) and then zoom out (TS_Z3-1-10), using 10 interpolation frames per zoom operation.

**Extended Data Figure 9.**
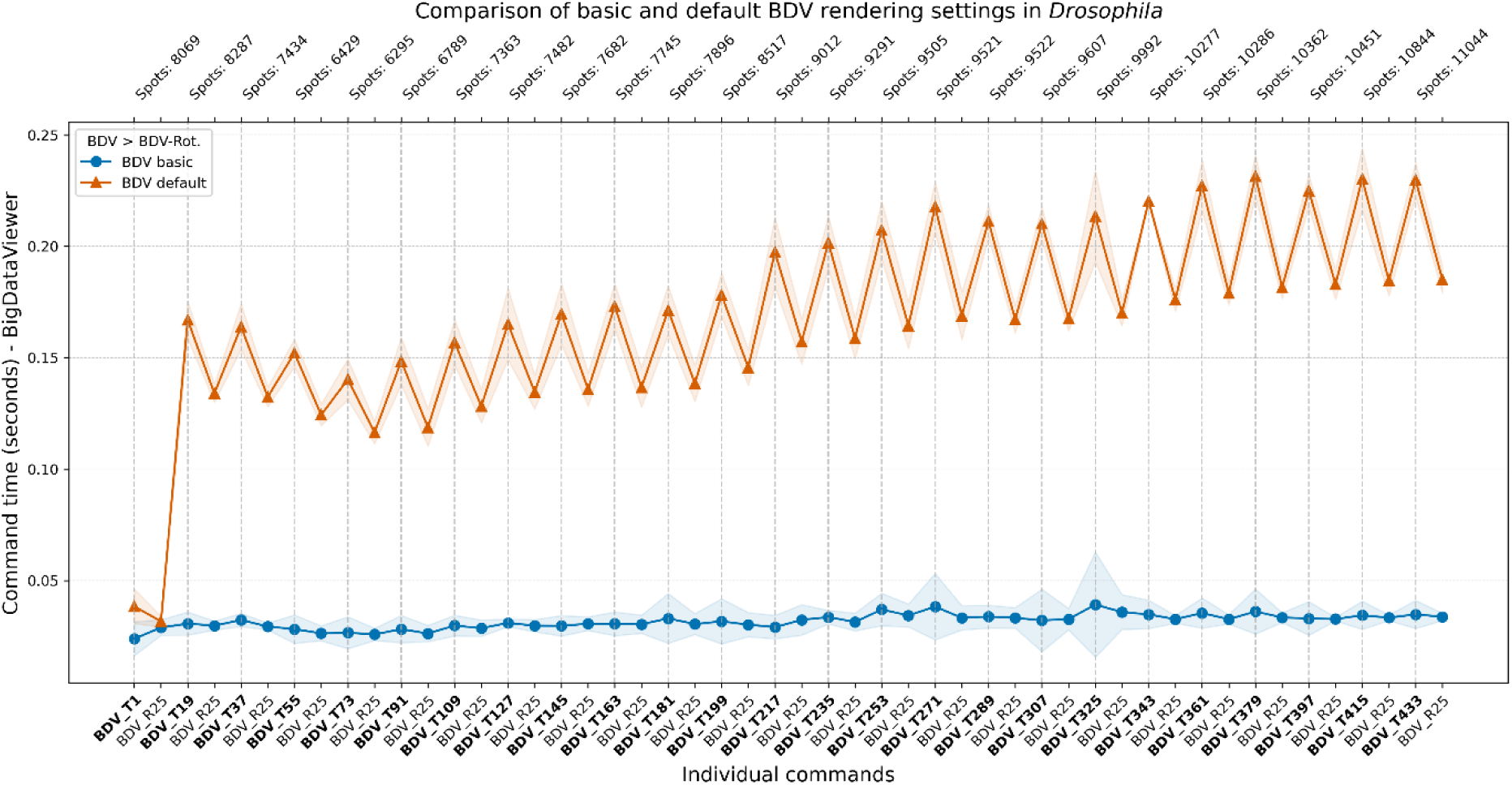
Performance of BDV window commands on a *Drosophila* dataset, measured as the time (in seconds) required to either render annotations (BDV_T*n*, where *n* is the time point) or perform a 360° rotation (BDV_R*n*, where *n* is the number of frames used for the rotation), using either Basic or Default rendering settings.

**Extended Data Figure 10.**
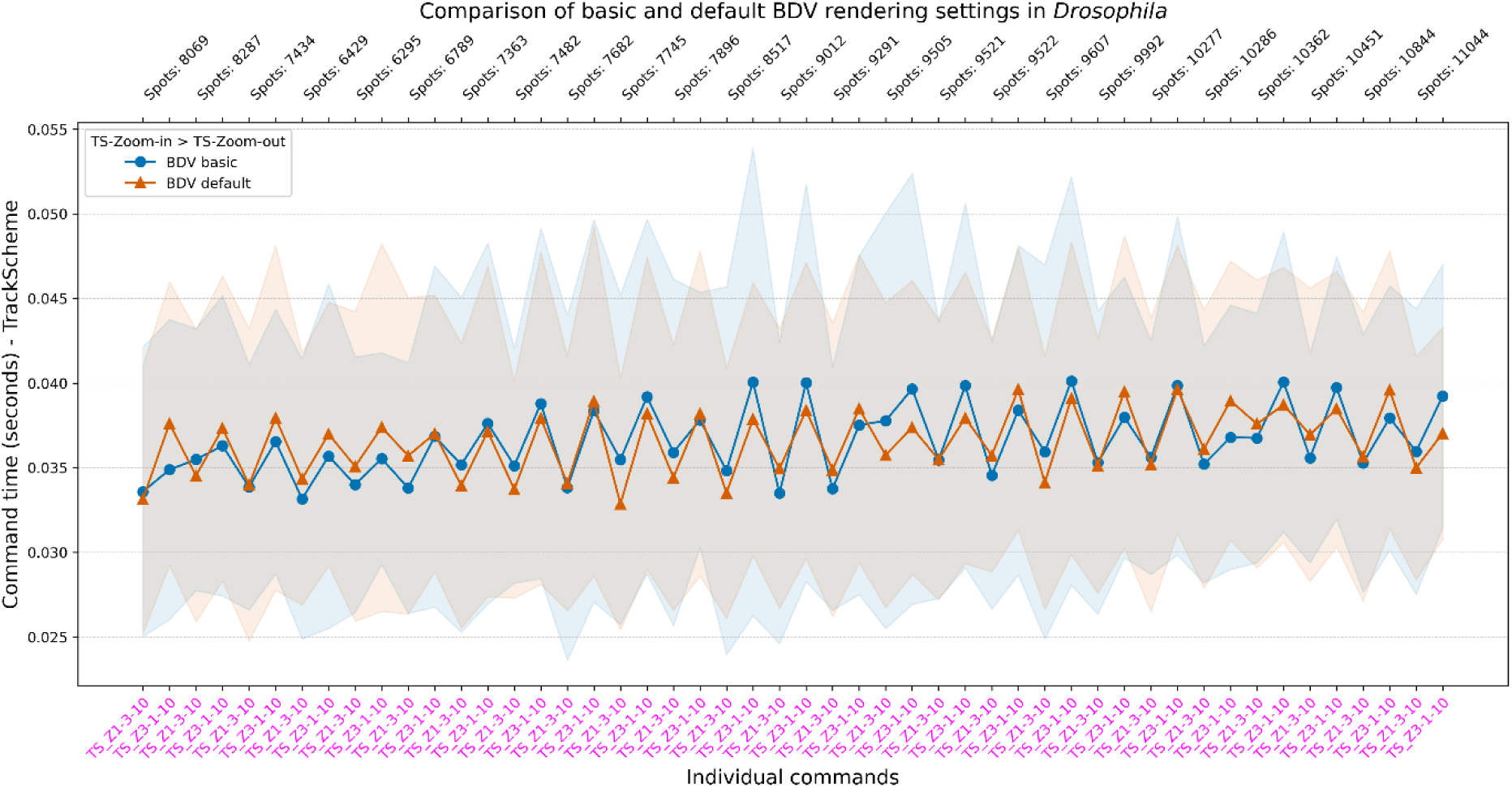
TrackScheme command performance on a *Drosophila* dataset, measured as the time (in seconds) required to zoom in (TS_Z1-3-10) and then zoom out (TS_Z3-1-10), using 10 interpolation frames per zoom operation.

**Extended Data Figure 11.**
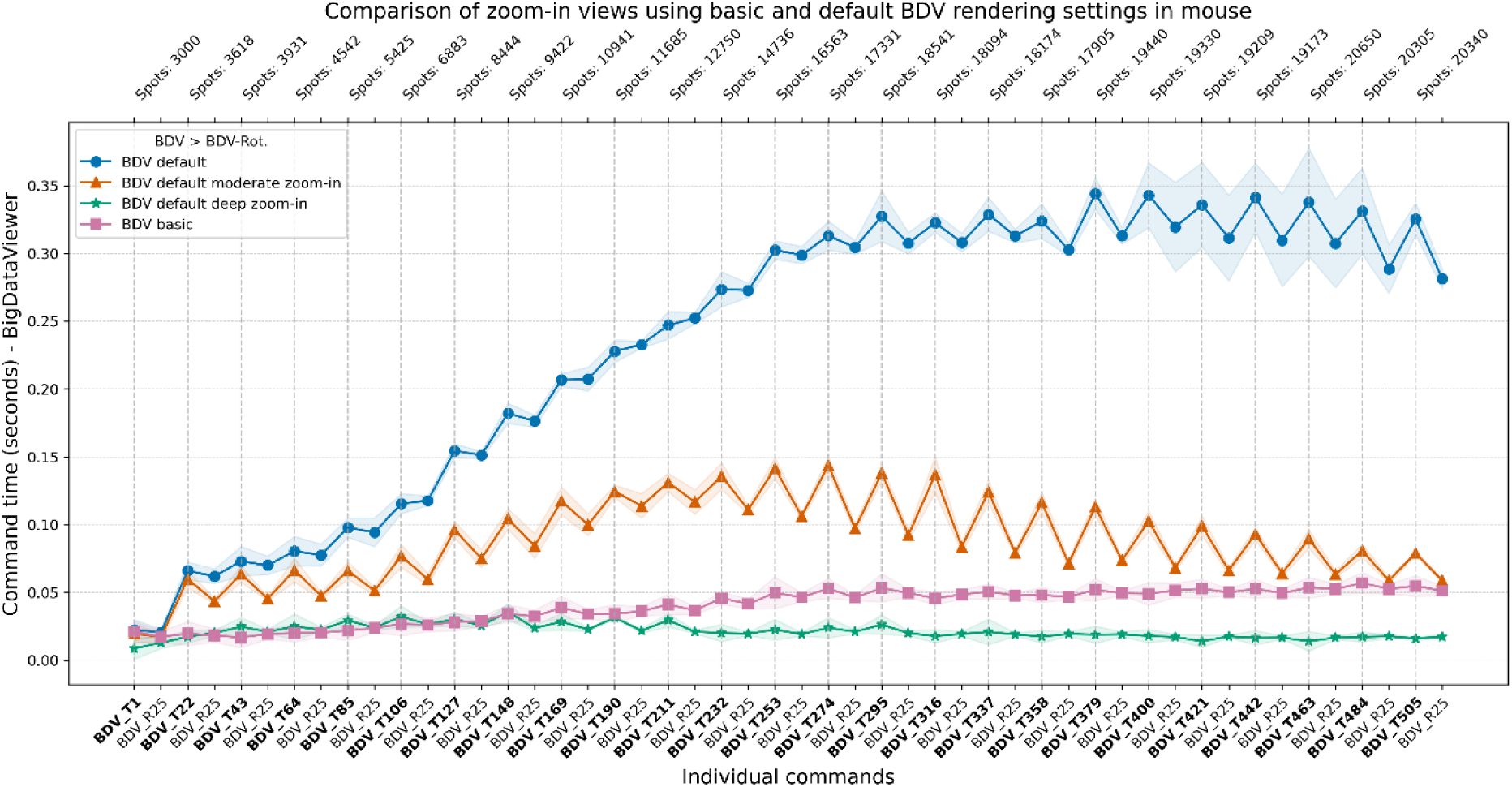
Performance of BDV window commands on a mouse dataset, measured as the time (in seconds) required to render annotations (BDV_Tn, where n is the time point), using Basic or Default rendering settings. For Default settings, performance was additionally measured at two zoom levels: moderate and deep zoom-in.

**Extended Data Figure 12.**
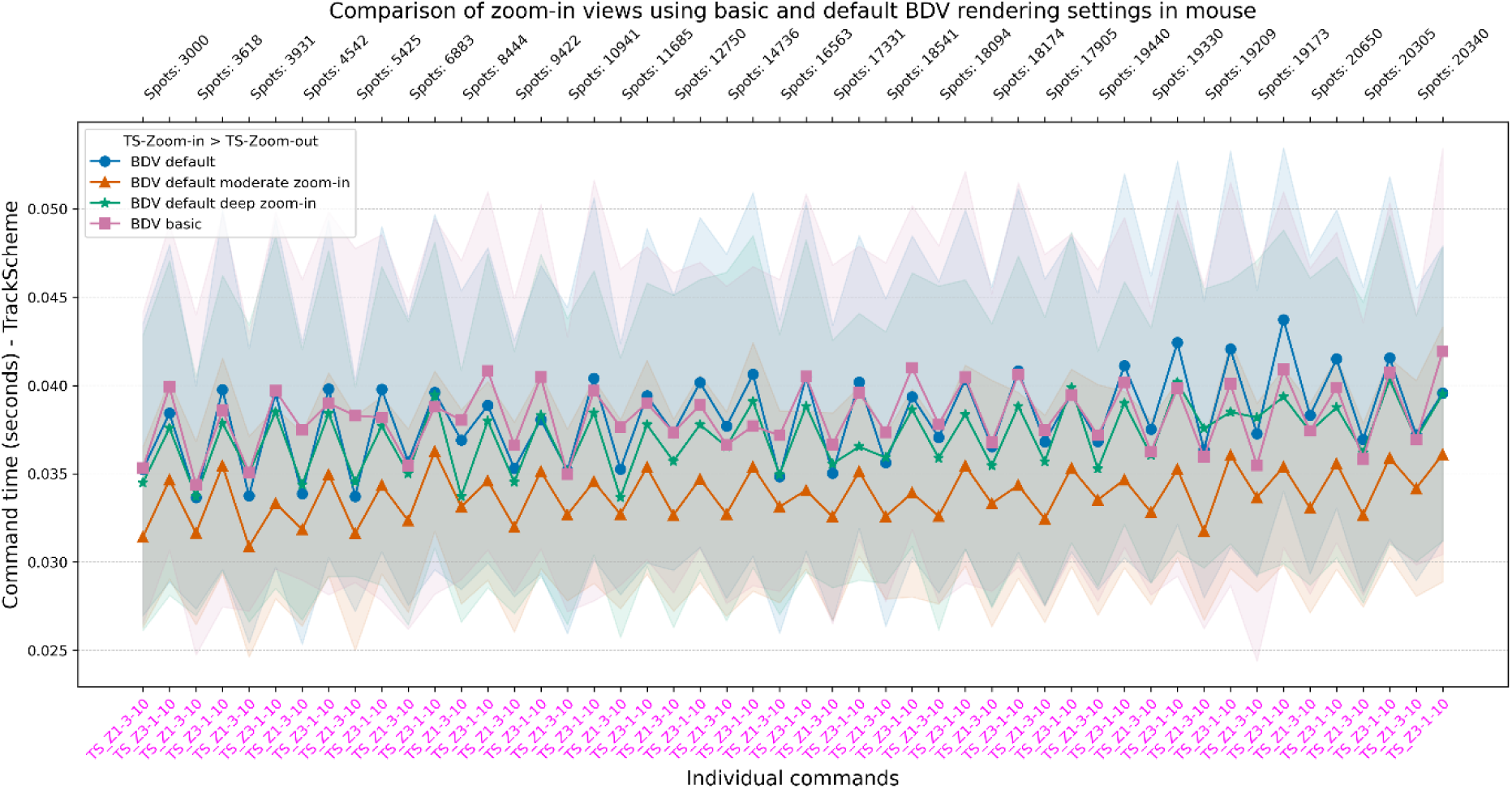
TrackScheme command performance on a mouse dataset, measured as the time (in seconds) required to zoom in (TS_Z1-3-10) and then zoom out (TS_Z3-1-10), using 10 interpolation frames per operation and Basic or Default BDV rendering settings. For Default settings, performance was additionally assessed at two zoom levels: moderate and deep zoom-in.

**Extended Data Figure 13.**
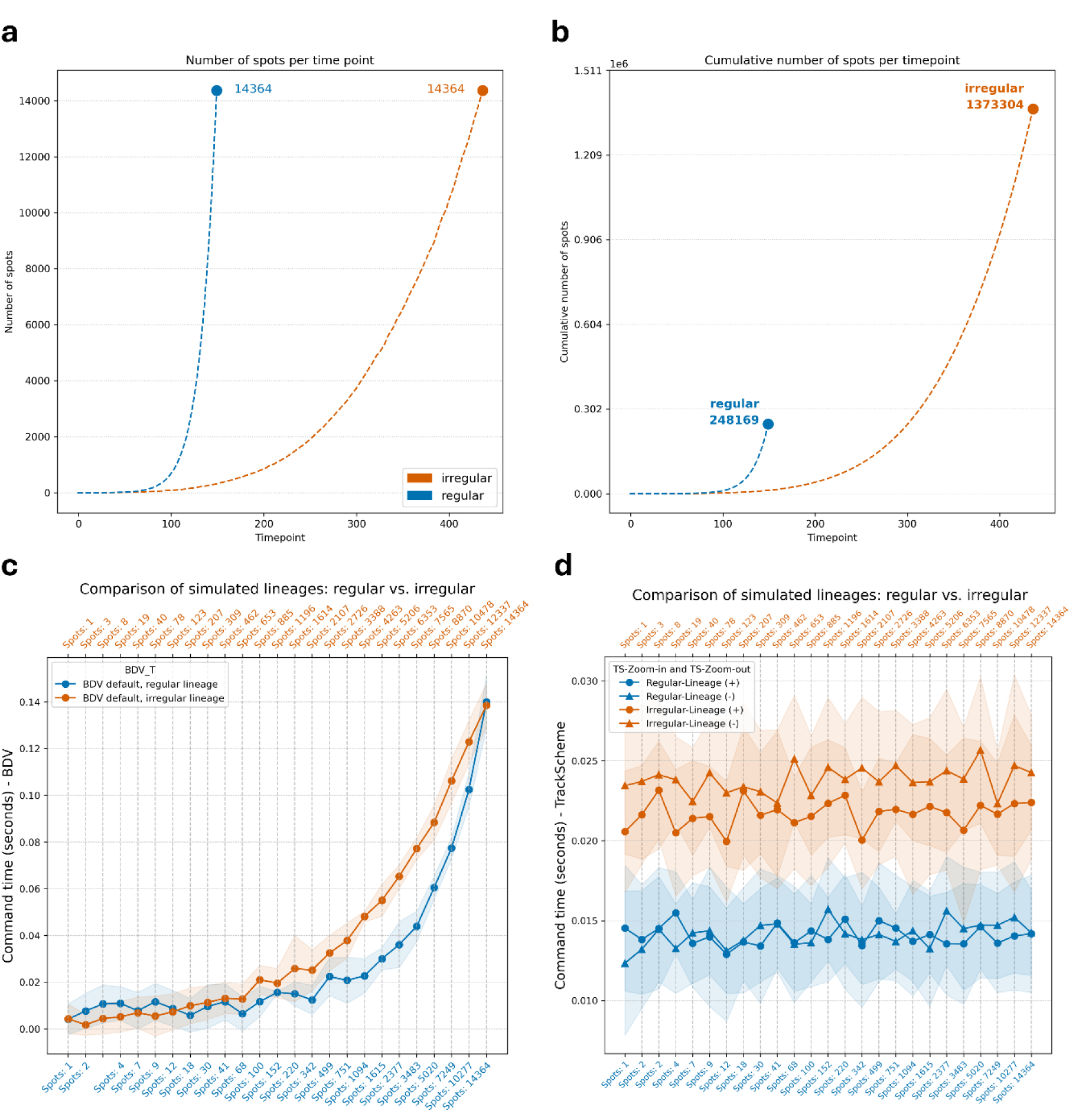
Performance comparison of regular versus irregular lineage datasets. Two simulations generated 14,364 artificial and linked annotation spots using the Mastodon *Simulator*. The regular lineage required only 150 timepoints, while the irregular lineage needed 437 **(a)**, resulting in a higher total number of accumulated annotations in the latter **(b)**. Despite this, benchmarking showed nearly identical performance in the BDV window at the final time point, when annotation counts matched **(c)**. TrackScheme performance consistently favored the regular lineage but only marginally, likely due to its lower overall annotation count **(d)**.

**Extended Data Figure 14.**
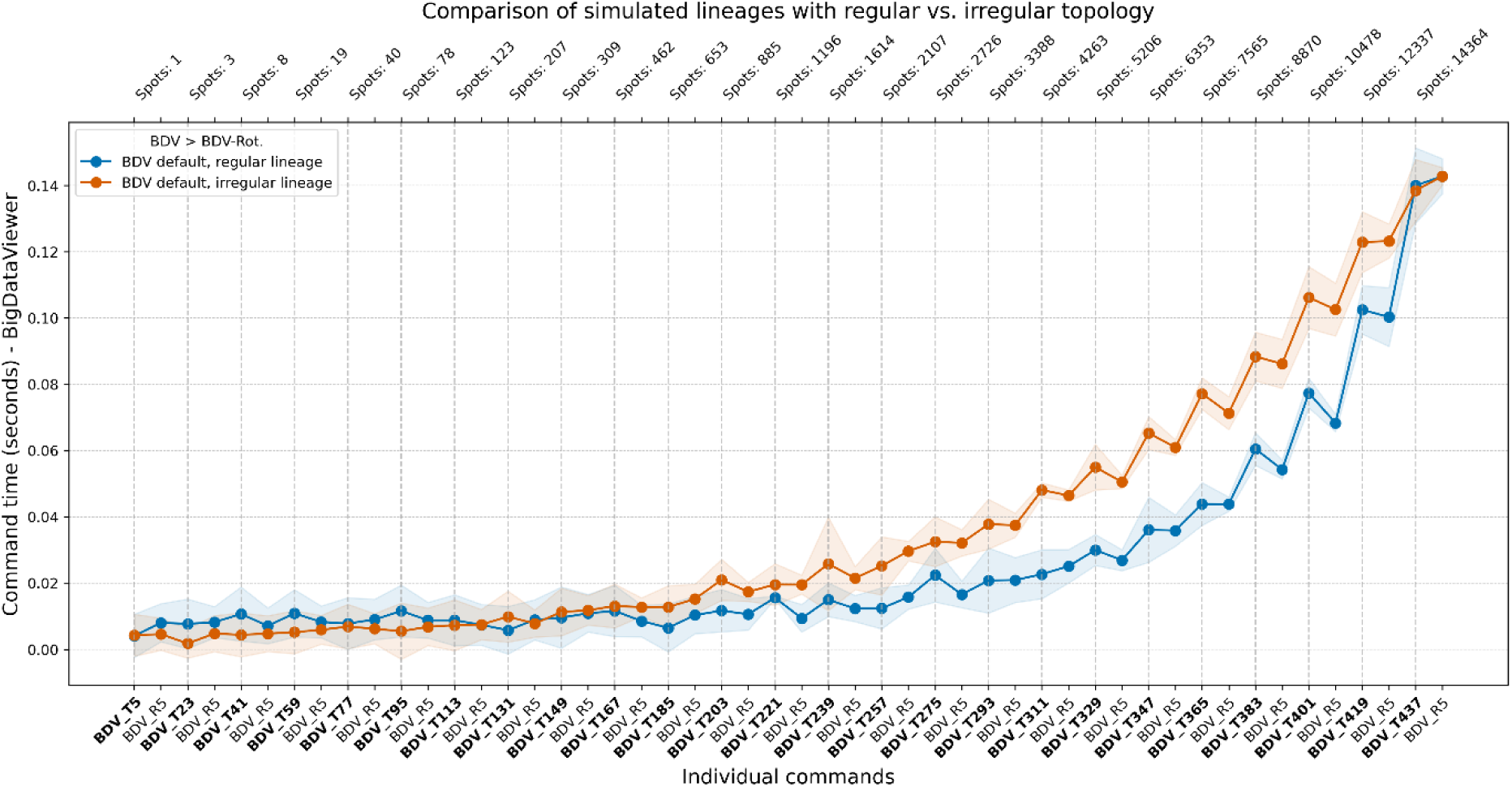
Performance of BDV window commands on two simulated datasets with either regular or irregular lineage topology, measured as the time (in seconds) required to either render annotations (BDV_T*n*, where *n* is the time point) or perform a 360° rotation (BDV_R*n*, where *n* is the number of frames used for the rotation), using Default rendering settings.

**Extended Data Figure 15.**
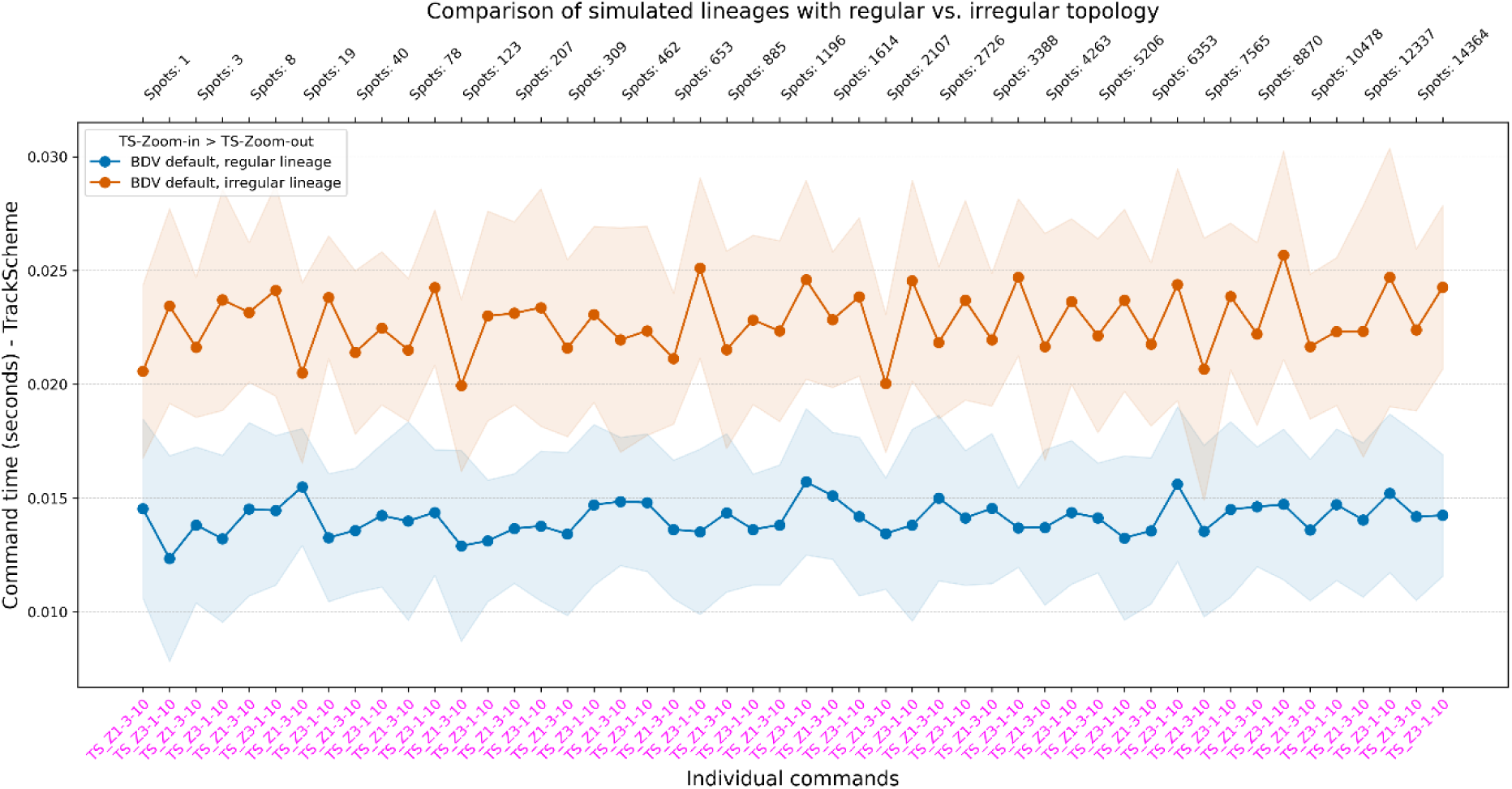
TrackScheme command performance on two simulated datasets with either regular or irregular lineage topology, measured as the time (in seconds) required to zoom in (TS_Z1-3-10) and then zoom out (TS_Z3-1-10), using 10 interpolation frames per zoom operation.

**Extended Data Video 1.**
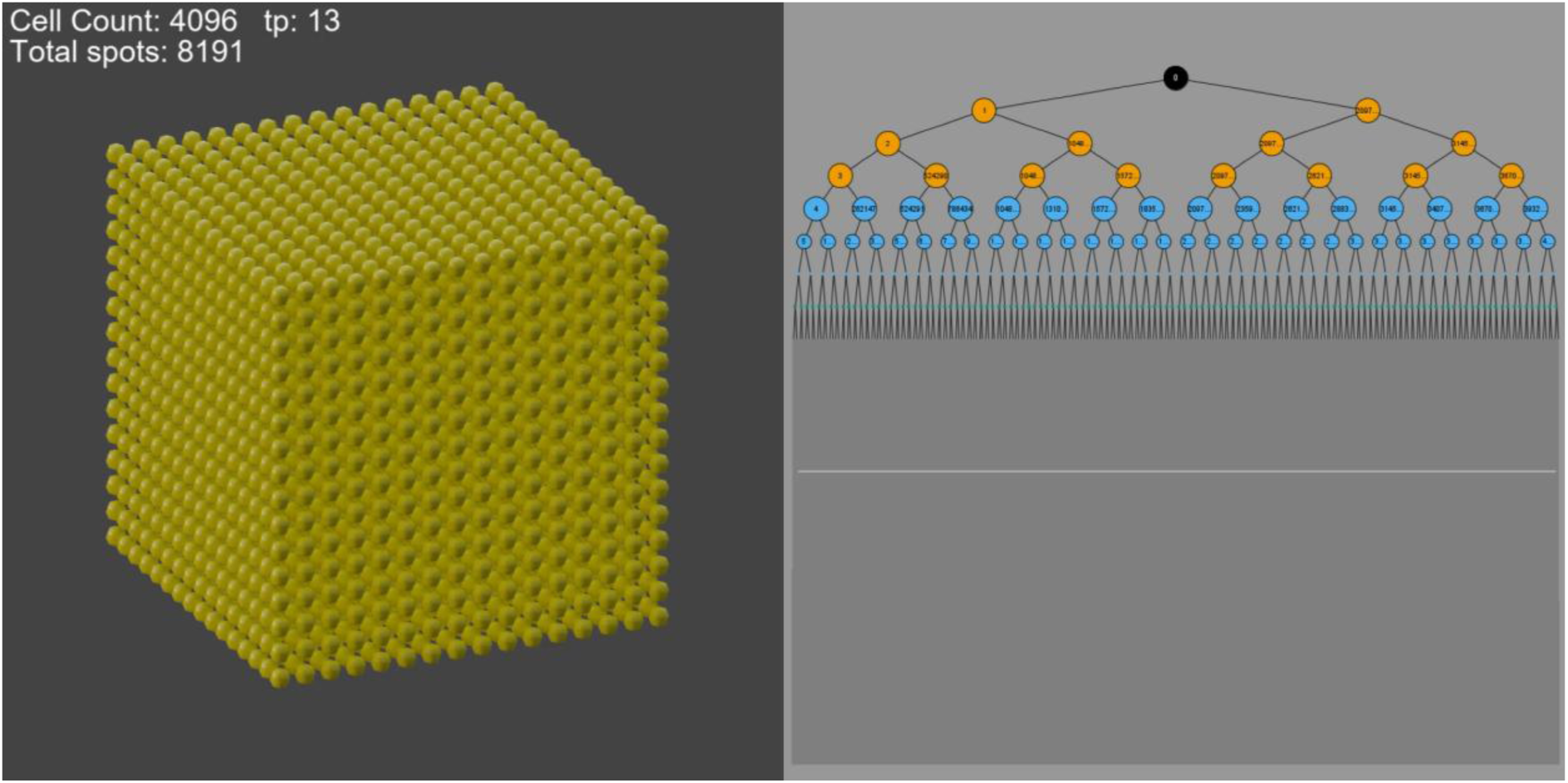
Synthetic Benchmark Data 1, generated as cubes that expand triaxially from a single spot annotation to 2,097,152 spots with a lineage. Each triaxial growth cycle is color-coded consistently. The Blender view with rendered cubic volumes is shown on the left; the corresponding lineage tree is displayed on the right.

**Extended Data Video 2.**
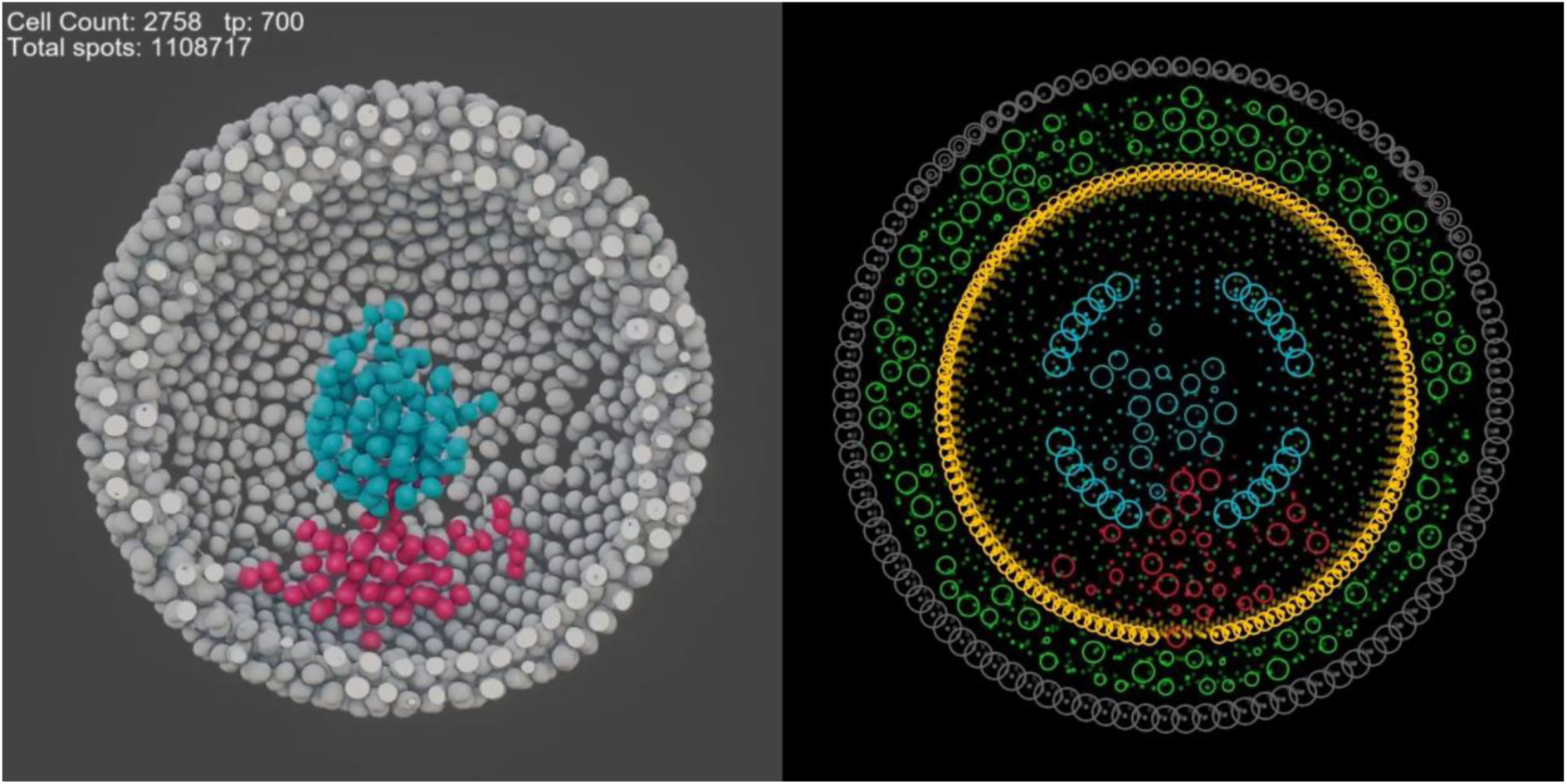
Dynamic, cell-like behaviors simulated in Mastodon. Rendered dataset in Blender (left). Scene elements, including nested spherical boundaries and seeded agents, were defined in the TrackScheme window and are depicted here as a single section with the BDV (right). After 500 simulation frames, epibolic-like expansion was completed and was subsequently followed by gastrulation-like inward migration through a central opening.

**Extended Data Video 3.**
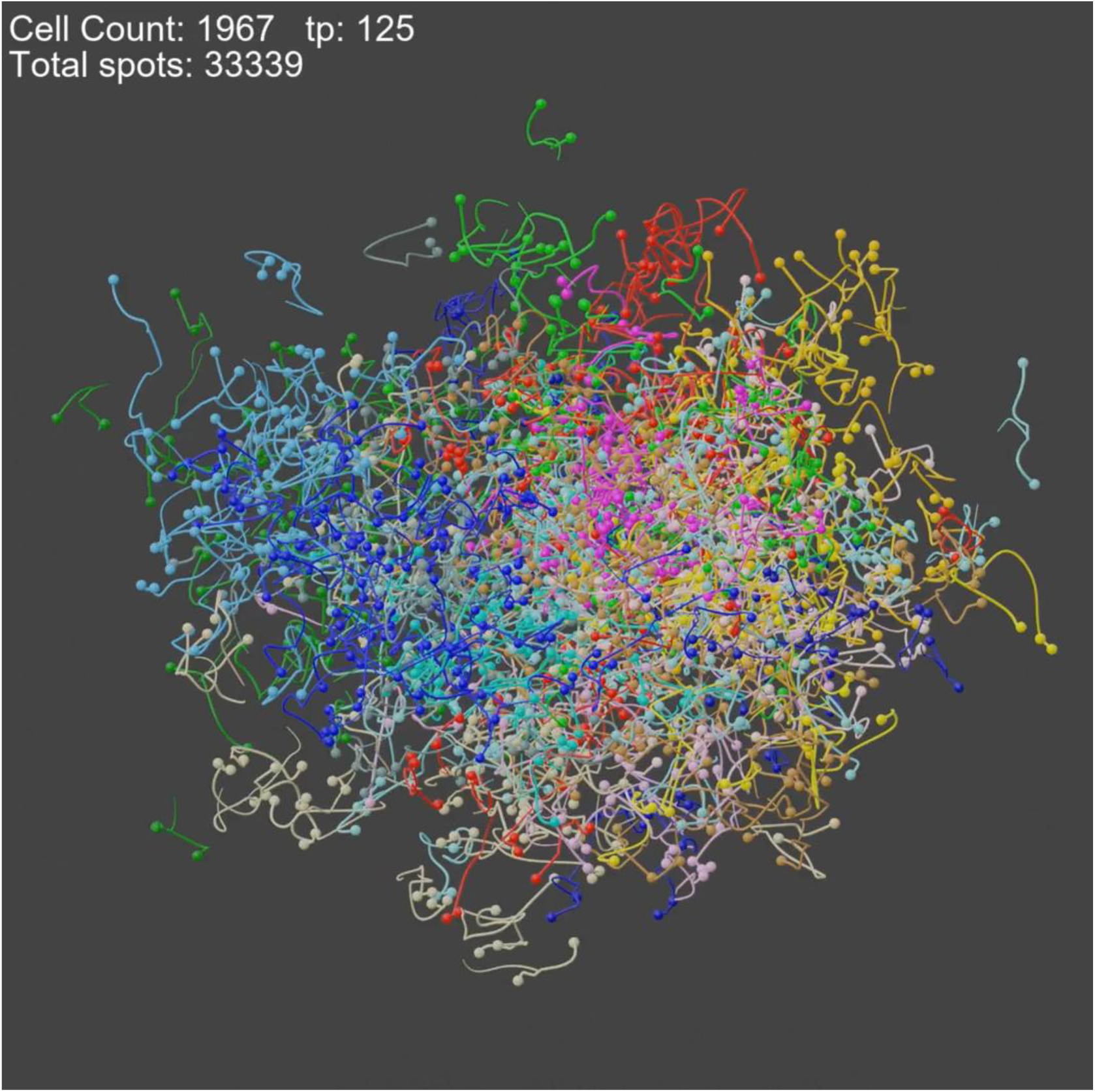
A spherical ball grown within Mastodon’s Simulator develops into a large-scale Mastodon project containing a total of 22 million spots.

**Extended Data Video 4.**
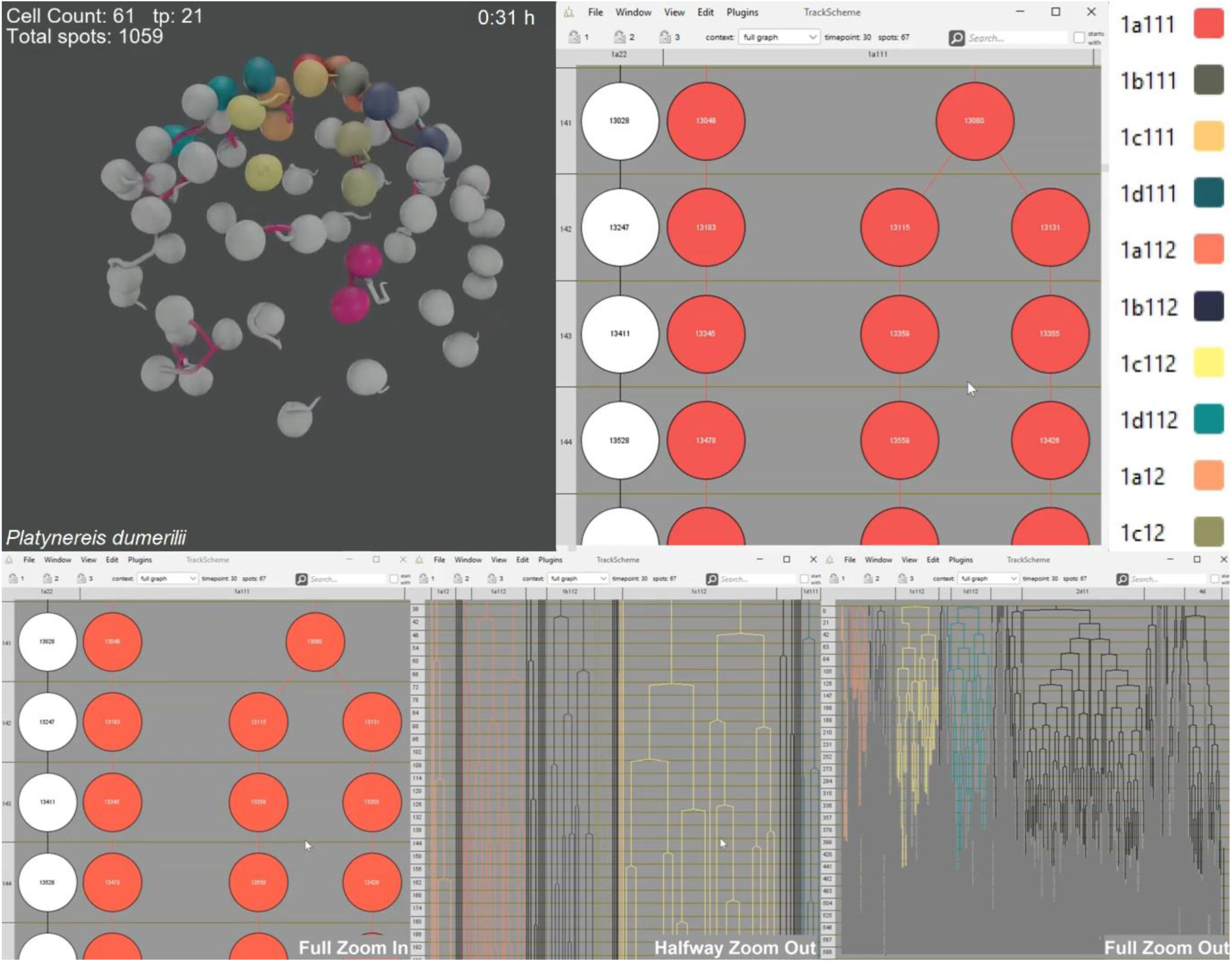
3D visualisation of the reconstructed cell lineage of the annelid *P. dumerilii* displaying 15 hr of embryogenesis. **Top Left:** Unfolding development rendered using Blender, a powerful open-source 3D creation suite, that we bridged with Mastodon. **Top Right:** Mastodon’s TrackScheme window showing a zoomed-out view of the fully reconstructed cell lineage of the annelid worm, *P. dumerilii*, primarily utilising semi-automated tracking. The complete tracking data comprises an aggregate of 212,468 spots. Cell divisions are superimposed in magenta. Descendants of brain progenitor cells 1q^111^,^,^ 1q^112^ (q = a, b, c, d) are labelled in different colours using Mastodon’s tag feature (see also legend on the right). **Bottom:** Depicted are three snapshots of the TrackScheme window during the zoom-out (Full Zoom In, Halfway Zoom Out and Full Zoom Out).

**Extended Data Video 5.**
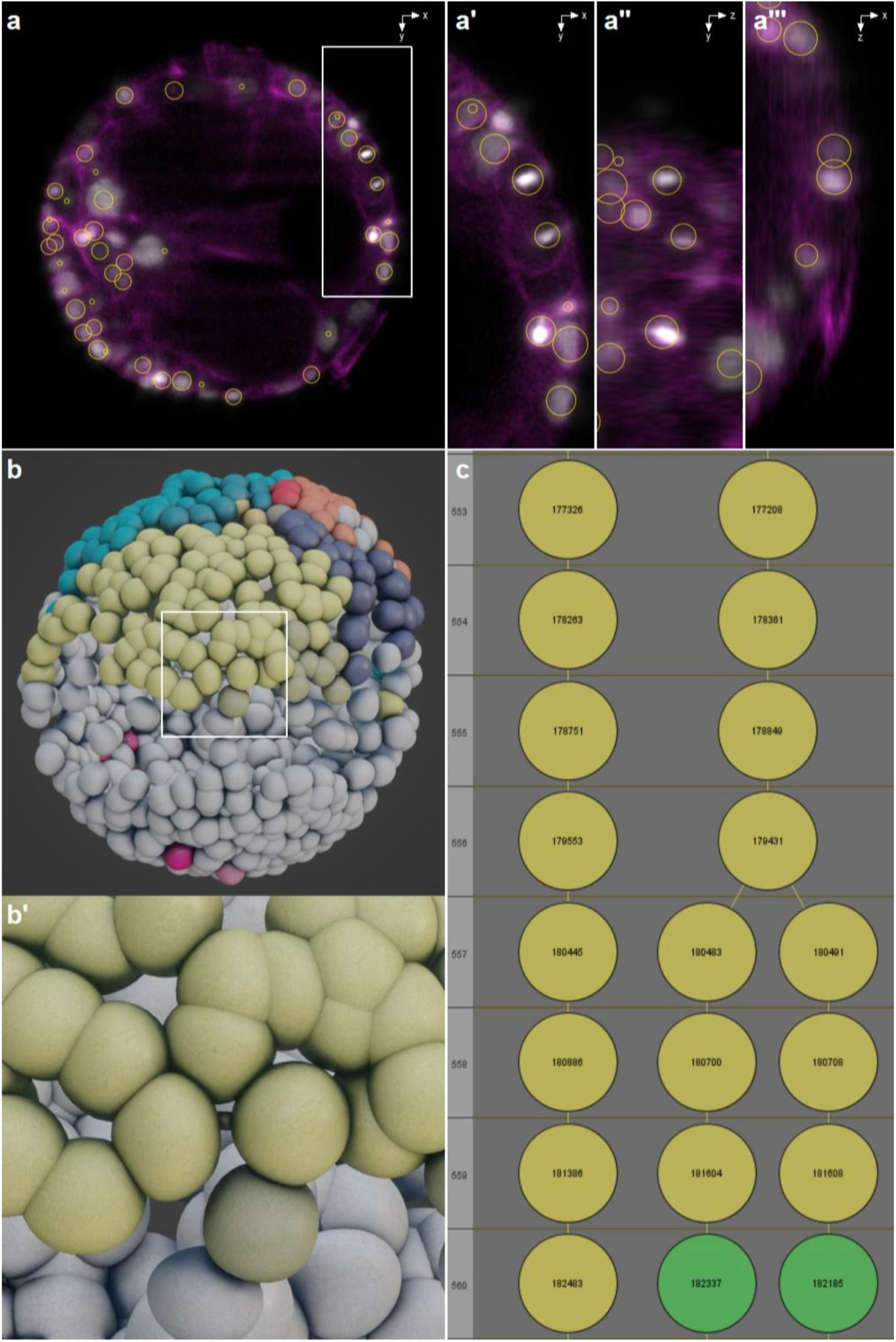
Different visualisations of the same branch split after using Mastodon’s automatic highlighting of cell divisions. **(a)** Mastodon’s BigDataViewer window shows some annotations. The white arrow points to a nucleus just prior to undergoing a division. (a’-a’’’) Zoomed-in area of (a) using three interconnected BigDataViewer windows (xy, yz, and xz) that simultaneously depict the same nuclei division from three different perspectives. **(b and b’)** A 3D visualisation of the same nuclei highlighted using a custom-made Mastodon extension to tag all cell divisions and rendered in Blender. **(c)** The splitting of the branches during the division of the nuclei as seen in Mastodon’s TrackScheme window.

**Extended Data Video 6.**
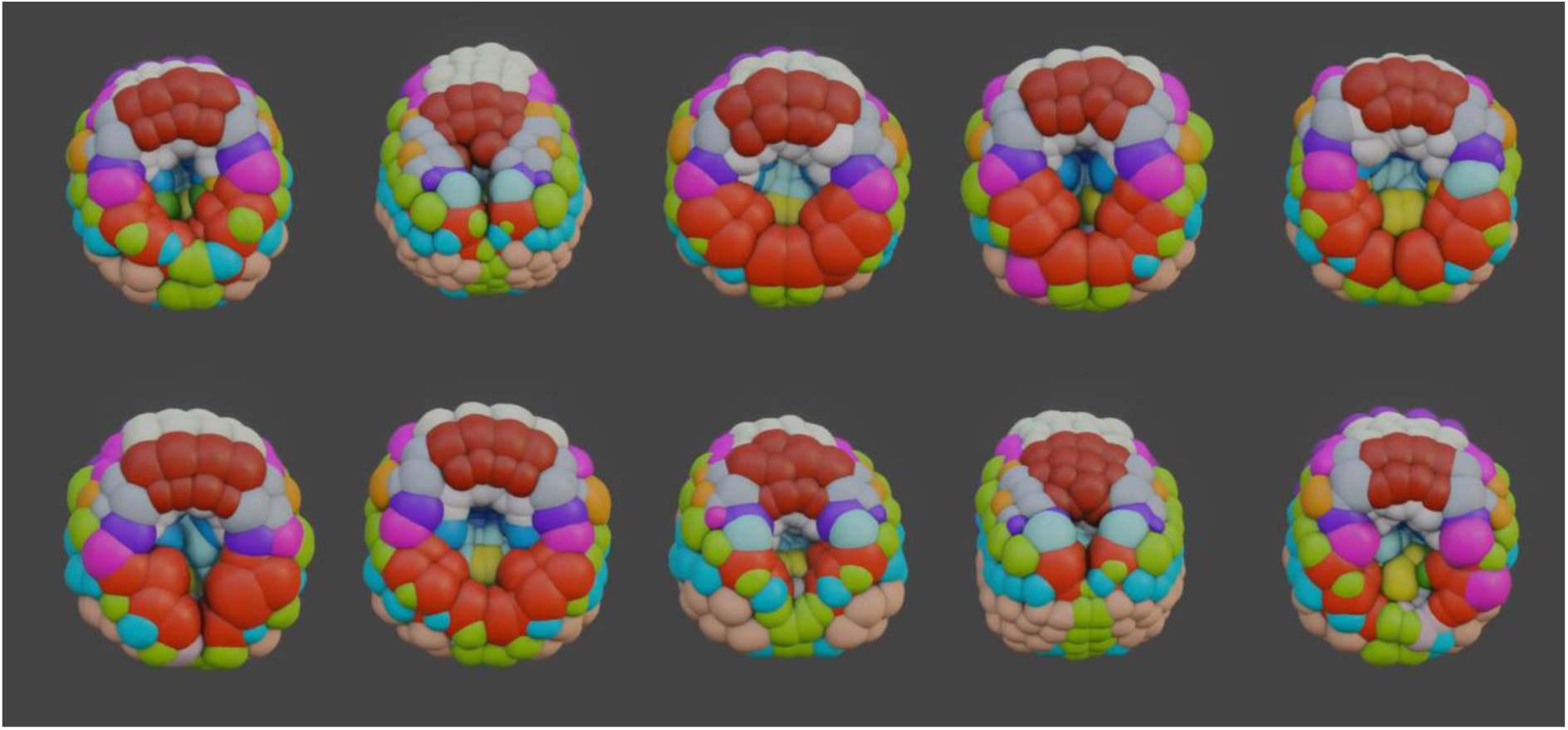
Ten embryos of previously published data on ascidian embryogenesis (Guignard, Fiúza et al. 2020. Science) have been imported into Mastodon, complete with various tracking information such as position, volume, and fate and are visualised here using 3D volume renderings generated using Blender.

**Extended Data Video 7.**
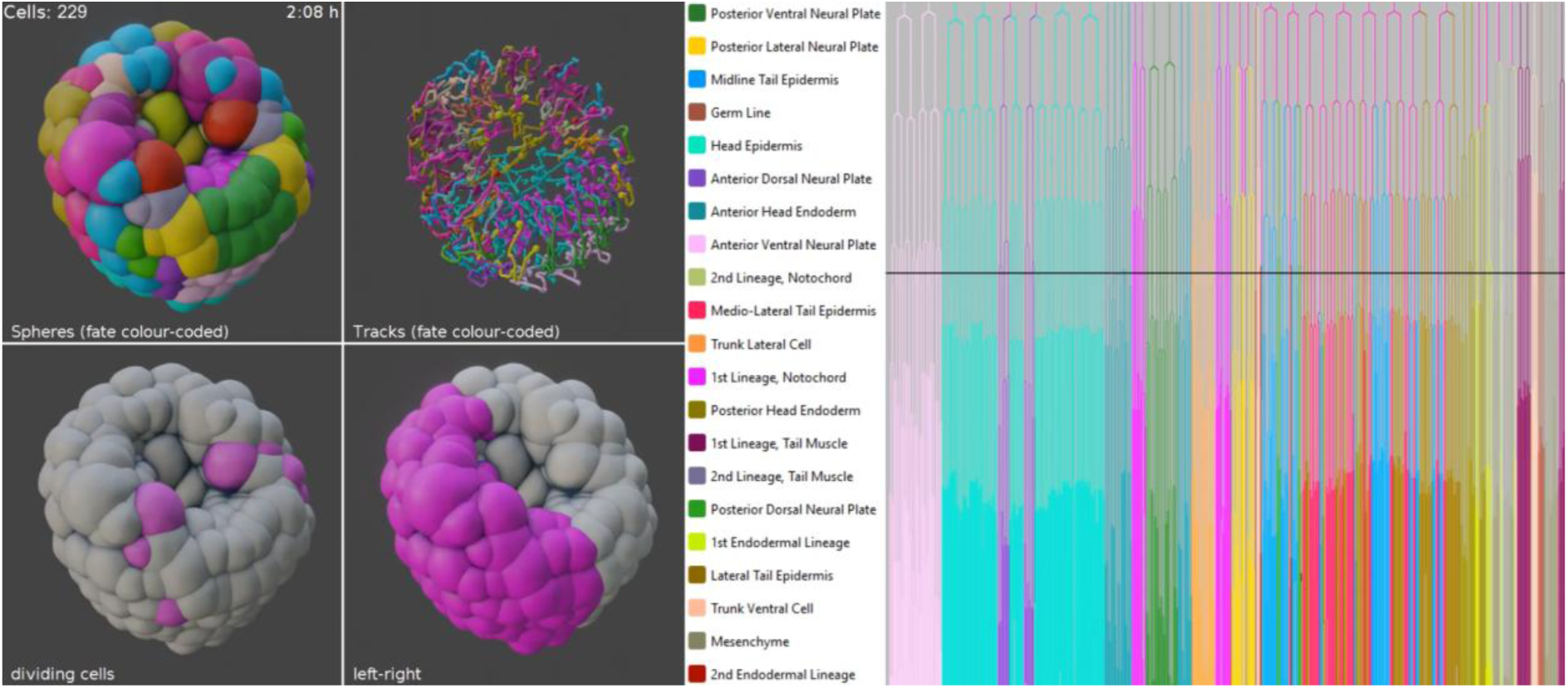
3D visualizations and corresponding lineage trees of embryo Astec-Pm-10, color-coded for fate 1 (random colors). The 3D renderings depict cell fates, trajectories, divisions, and left-right symmetry (left), while the TrackScheme window shows the corresponding timeline, marked by a black line (right).

**Extended Data Video 8.**
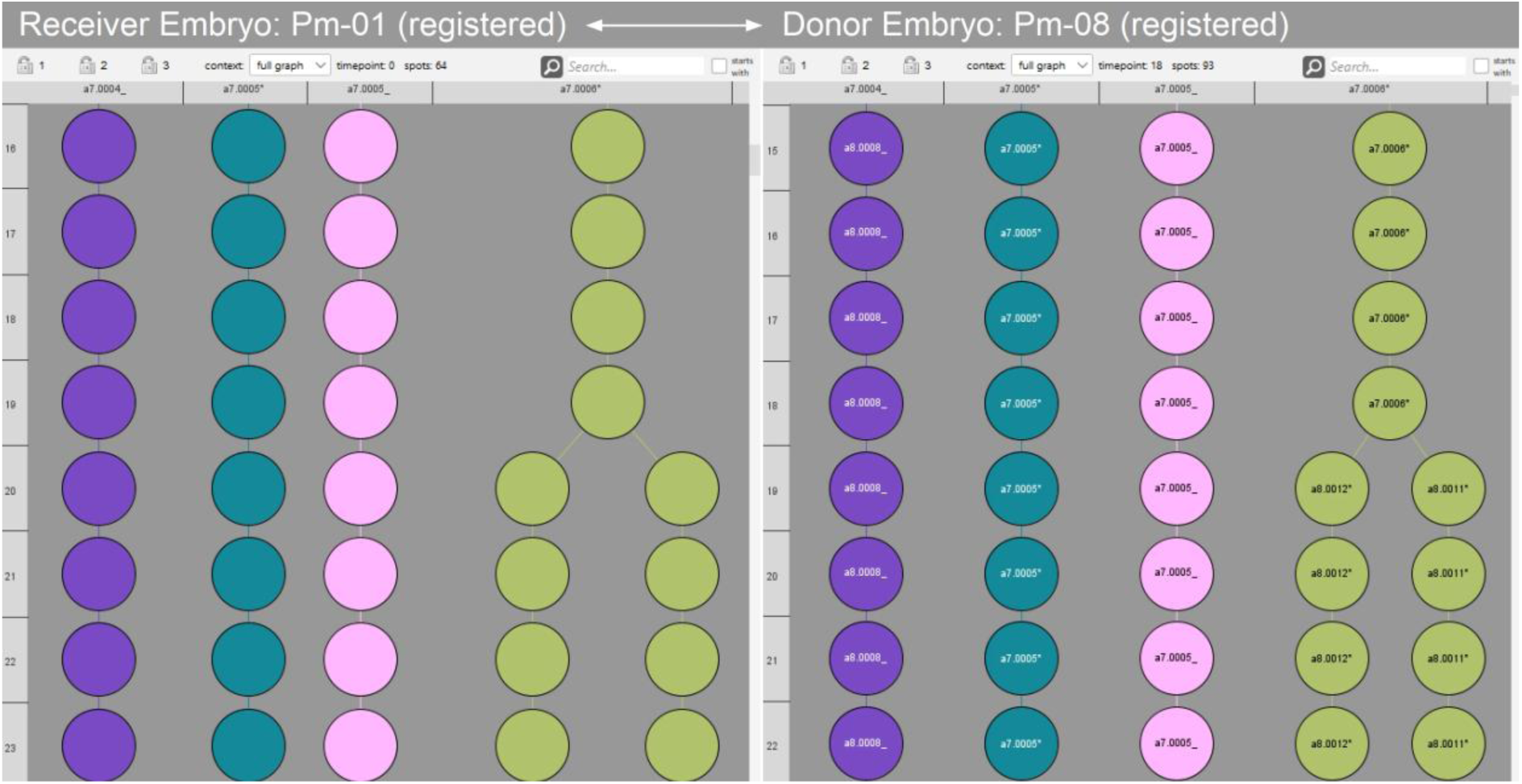
This video uses published tracking data on ascidian embryogenesis (Guignard, Fiúza et al., 2020, Science). The TrackScheme windows of ascidian embryos Pm-01 (left) and Pm-08 (right) illustrate the automatic transfer of color-tags and spot labels from one embryo to the other using Mastodon’s registration plugin.

**Extended Data Video 9.**
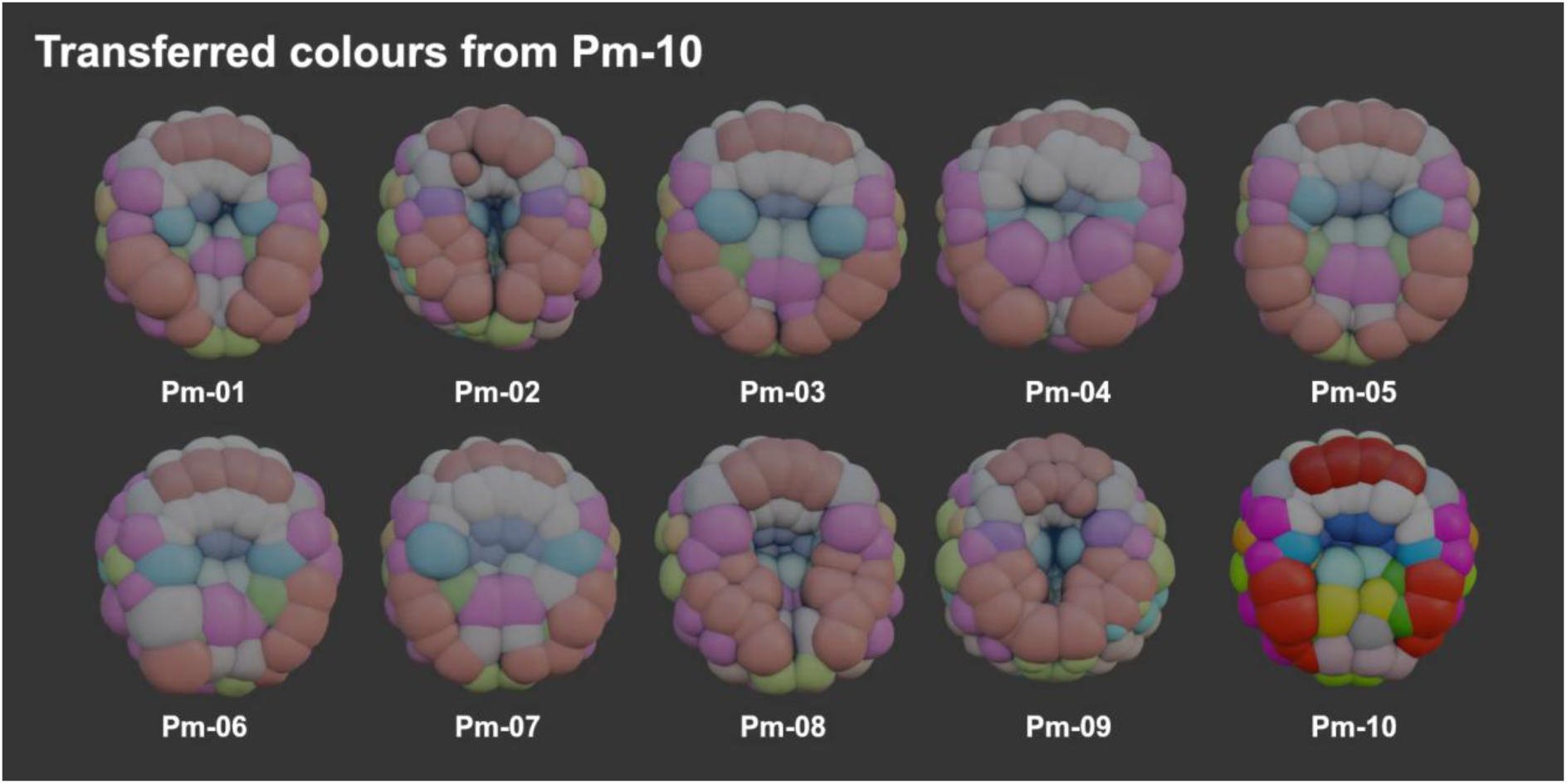
This video uses published tracking data on ascidian embryogenesis (Guignard, Fiúza et al., 2020, Science). Three-dimensional renderings of all ascidian embryos imported into Mastodon were generated using Blender. In this scenario, embryo Pm-10 serves as the donor. Using Mastodon’s registration plugin, colour-tags and labels were automatically transferred to the remaining nine embryos.

**Extended Data Video 10.**
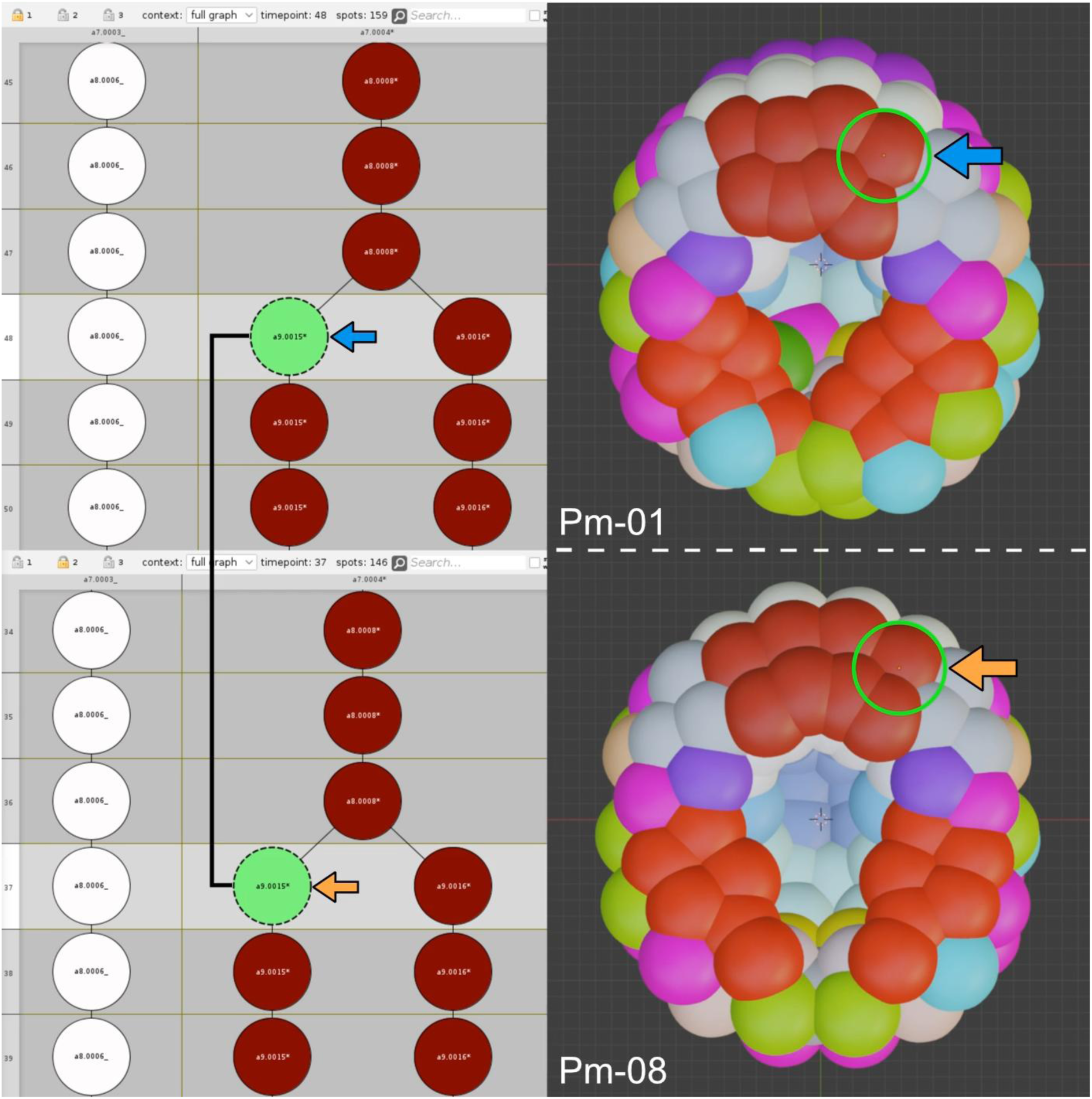
This video uses published tracking data on ascidian embryogenesis (Guignard, Fiúza et al., 2020, Science) to showcase the capabilities of Mastodon’s Spatial Track Matching plugin. The plugin automatically recognises geometric information from cell divisions in datasets from two embryos and applies a track matching algorithm, based on the assumption that the embryos develop in a stereotyped manner. As a result, the plugin links the two cell lineage trees, enabling interactive real-time data exploration and comparison between the datasets. The video features two embryos: Embryo 1 (Astec-Pm-01) at the top and Embryo 08 (Astec-Pm-10) at the bottom. Both embryos have been imported into Mastodon, including tracking information such as cell positions over time, volumes, and fates. The tracks are visualised in the TrackScheme window, while the embryos are displayed in 3D using an interactive Blender window. The Blender view responds to spot selections made in TrackScheme and vice versa, allowing bidirectional interaction between the lineage tree and the 3D visualisation.

**Extended Data Video 11.**
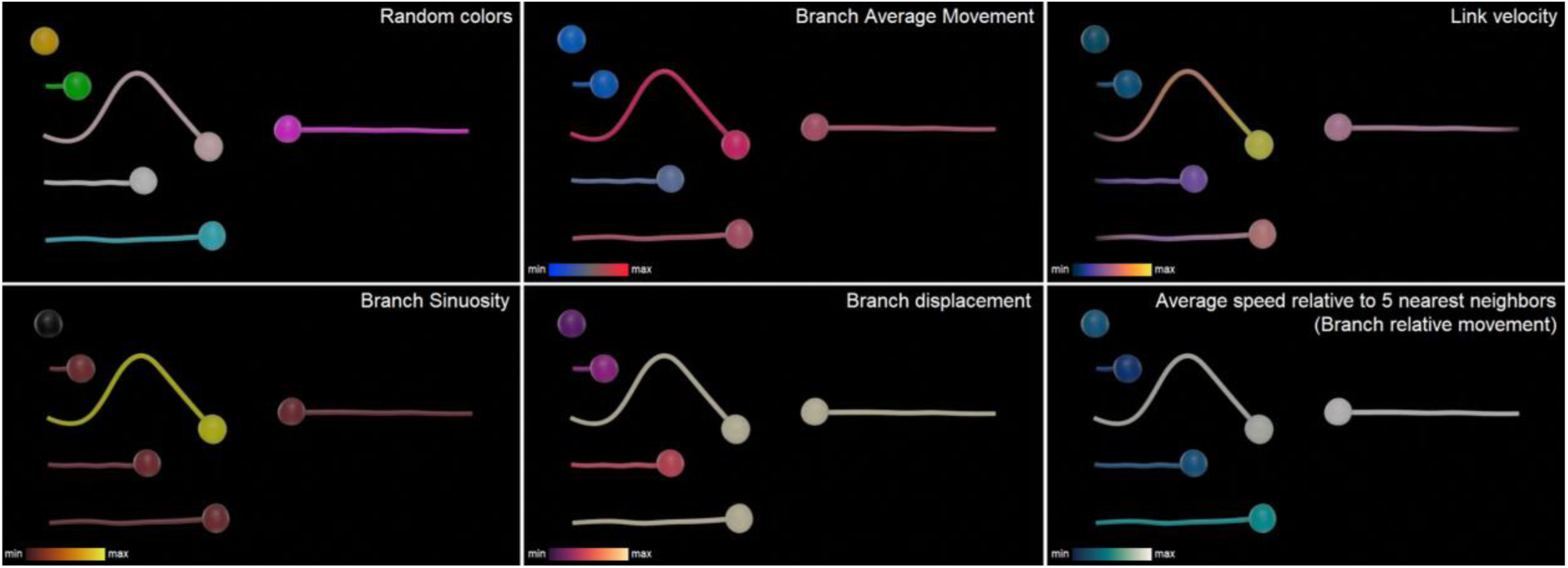
Visualization of Numerical features Using Mastodon and Blender. A simple artificial dataset was created in Mastodon by manually ‘drawing’ short tracks (15 timepoints) to demonstrate a selection of numerical features that can be visualised using Mastodon’s ‘Feature Color Modes’. The examples, shown from top left to bottom right, include: branch average movement, link velocity, average movement speed of cells relative to their five nearest neighbours, branch sinuosity, and branch displacement. Three-dimensional renderings were generated using Blender.

**Extended Data Video 12.**
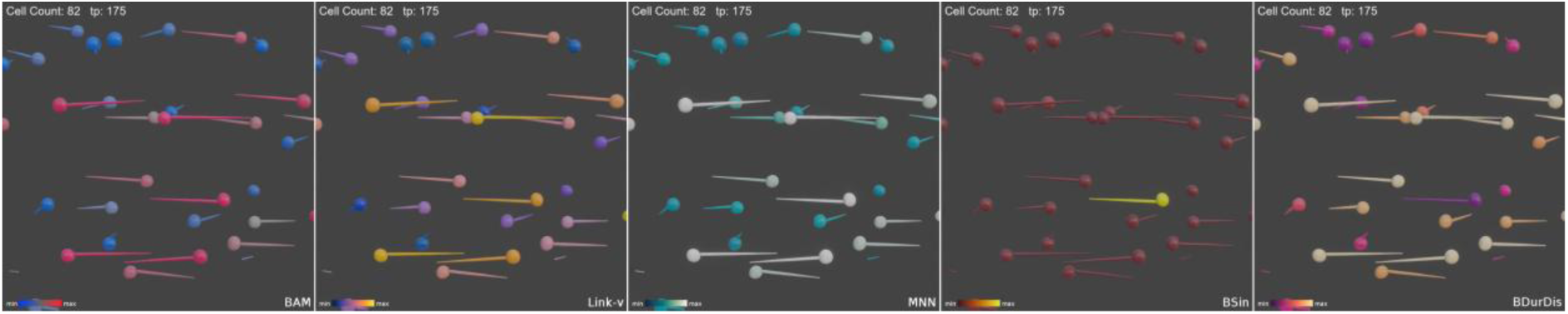
Simulation of lymphocyte motility using the <u>MotilityLab</u> track generator. In this dataset, cell divisions are absent, as this functionality is not supported by MotilityLab. The dataset features several calculated numerical features: Branch Average Movement (BAM), Link Velocity (Link-v), the average movement speed of cells relative to their five nearest neighbours (MNN), Branch Sinuosity (BSin), and Branch Duration and Displacement (BDurDis). The Blender-rendered dataset visualised in Extended Data Video 13 is an inset of the full simulation shown here, and the cell count indicated in the video refers to the entire simulation.

**Extended Data Video 13.**
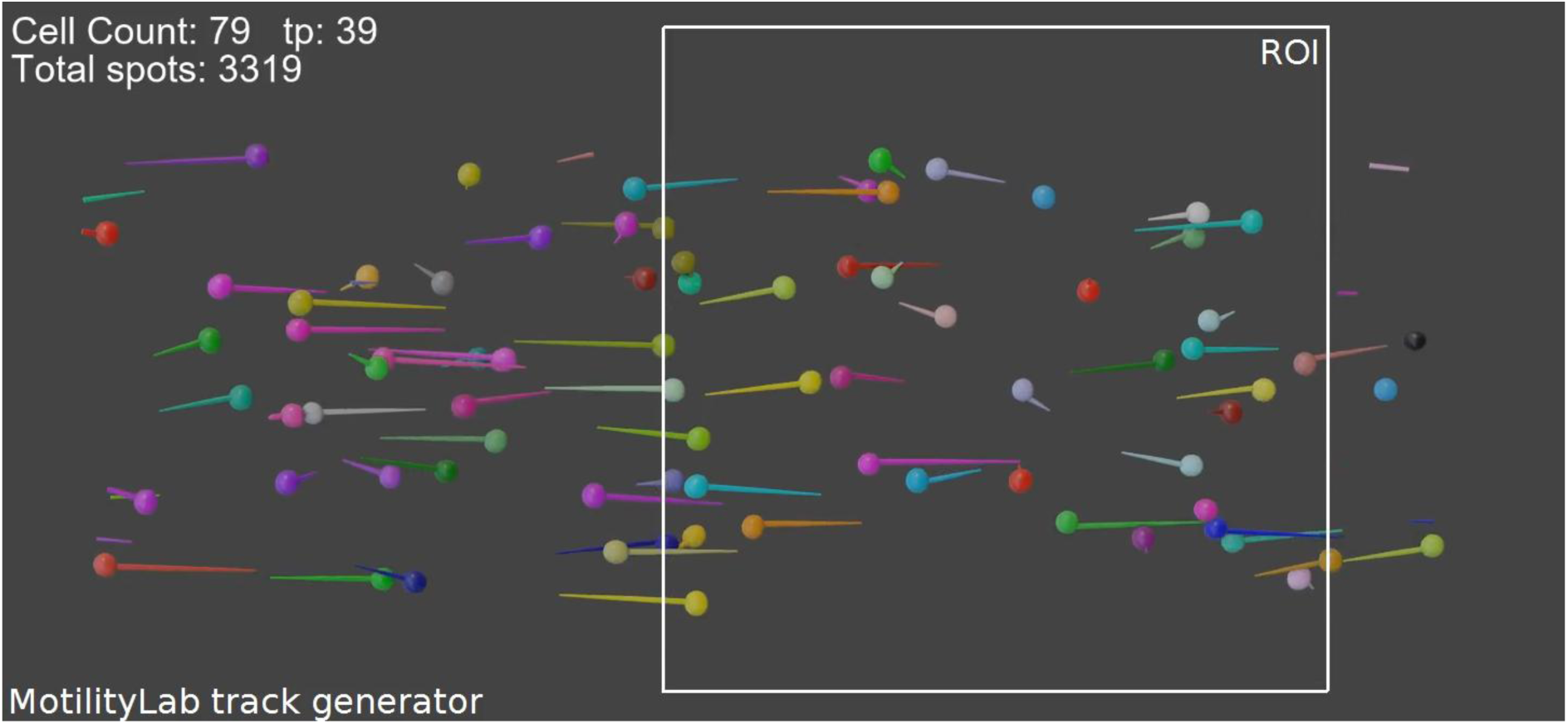
Full field-of-view simulation of lymphocyte motility using the <u>MotilityLab</u> track generator. The Blender-rendered dataset visualised in Extended Data Video 12 is an inset of the full simulation shown here, and the cell count indicated in the video refers to the entire simulation.

**Extended Data Video 14.**
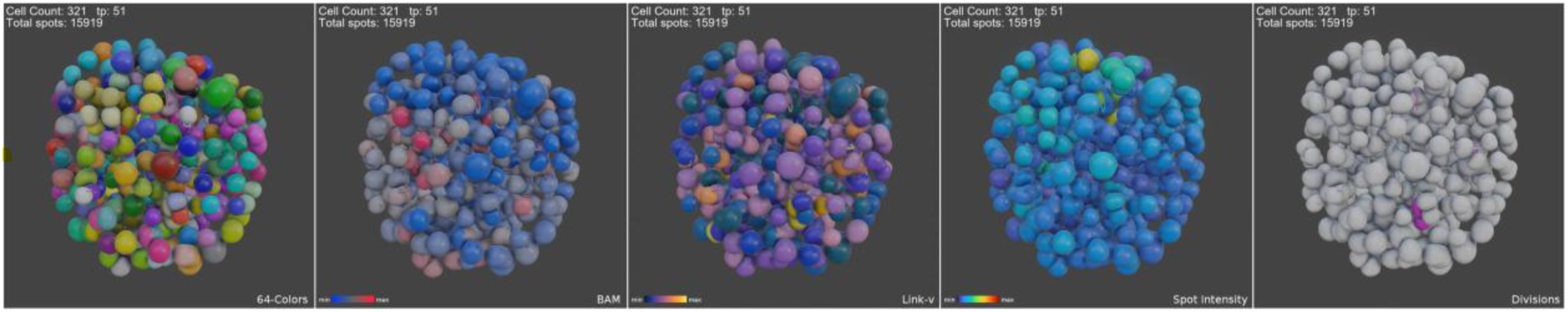
Example of a semi-automatically tracked Spheroid. A total of 143 time points were semi-automatically tracked by Carlos O. Gomez from the lab of Prof. Otger Campàs at TU Dresden, Germany. During the tracking period, the number of cells in the spheroid increased from 292 to 371. In addition to Branch Average Movement (BAM) and Link Velocity (Link-v), spot intensity features were extracted from the image data and incorporated into the visualisation. Link-v visualisations have dark spots whenever a new track starts as velocity calculations are based on links (edges) and need at least two spots (vertices).

**Extended Data Video 15.**
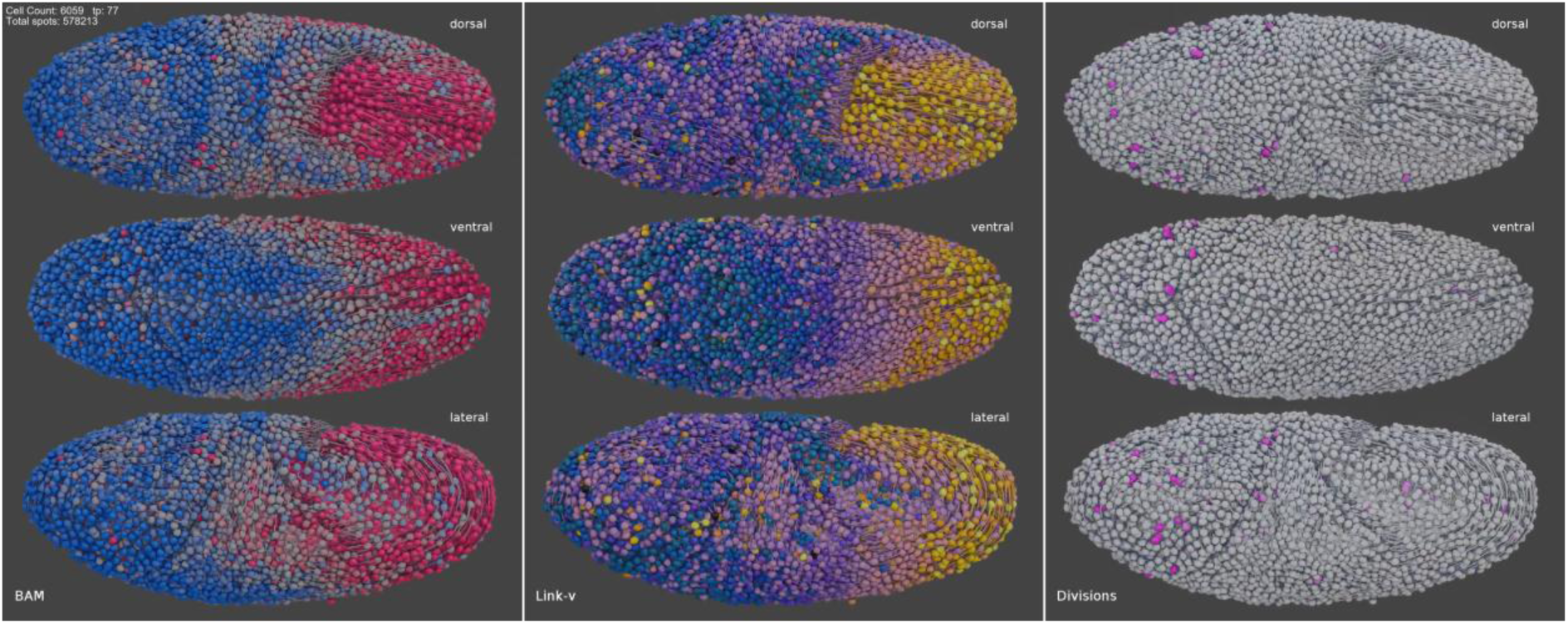
**3D renderings of the most extensive publicly available tracking datasets for the fruitfly *Drosophila melanogaster***. Visualisations show Branch Average Movement (left), Link Velocity (middle), and cell divisions (right) across a total of nearly 4 million automatically detected spots and links, with up to 10,000 spot detections in a single timepoint. Note that in the Link Velocity visualisation, dark spots appear where new tracks begin, as velocity calculations are based on links (edges) and require at least two spots (vertices). Tracking dataset from Malin-Mayor et al., 2023.

**Extended Data Video 16.**
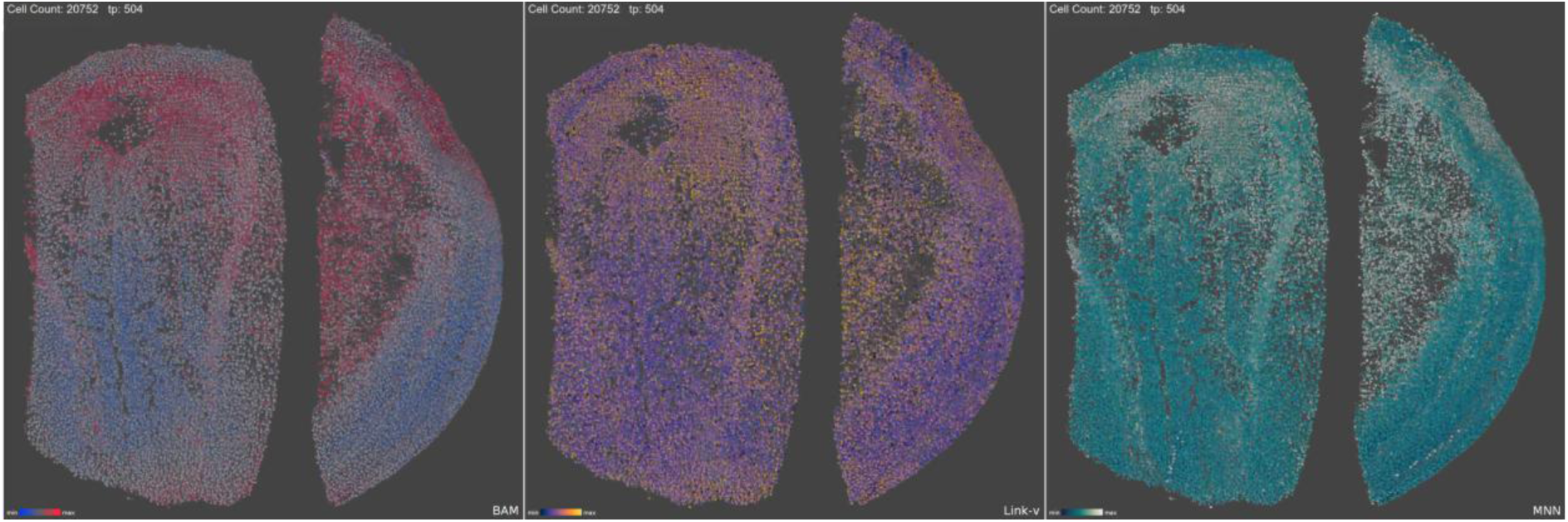
3D renderings using the most extensive publicly available lineage trees of the mouse tracking dataset. Visualisations show Branch Average Movement (left), Link Velocity (middle), and the average movement speed of cells relative to their five nearest neighbours (right), across a total of nearly 7.2 million automatically detected spots and links, with up to 21,000 spot detections in a single time point. Note that in the Link Velocity visualisation, dark spots appear where new tracks begin, as velocity calculations are based on links (edges) and require at least two spots (vertices). Tracking dataset from Malin-Mayor et al., 2023.

**Extended Data Table 1.**
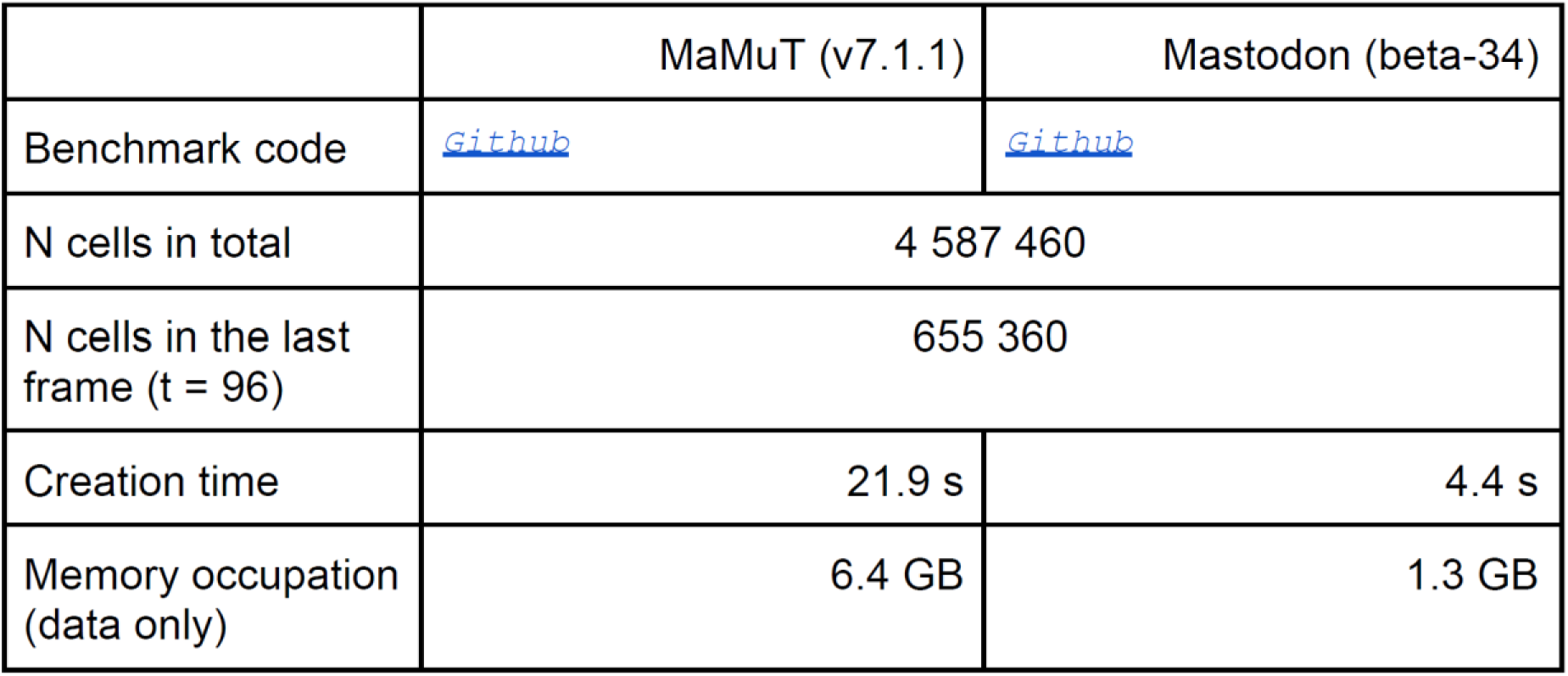
Data creation benchmark. MaMuT or Mastodon data structures are used to create programmatically a synthetic dataset where 10 starting cells divide 17 times, every 5 frames, resulting in the total number of cells reported in the table. In the case of MaMuT, the image view and the TrackScheme view crash or become unresponsive.

**Extended Data Table 2.**
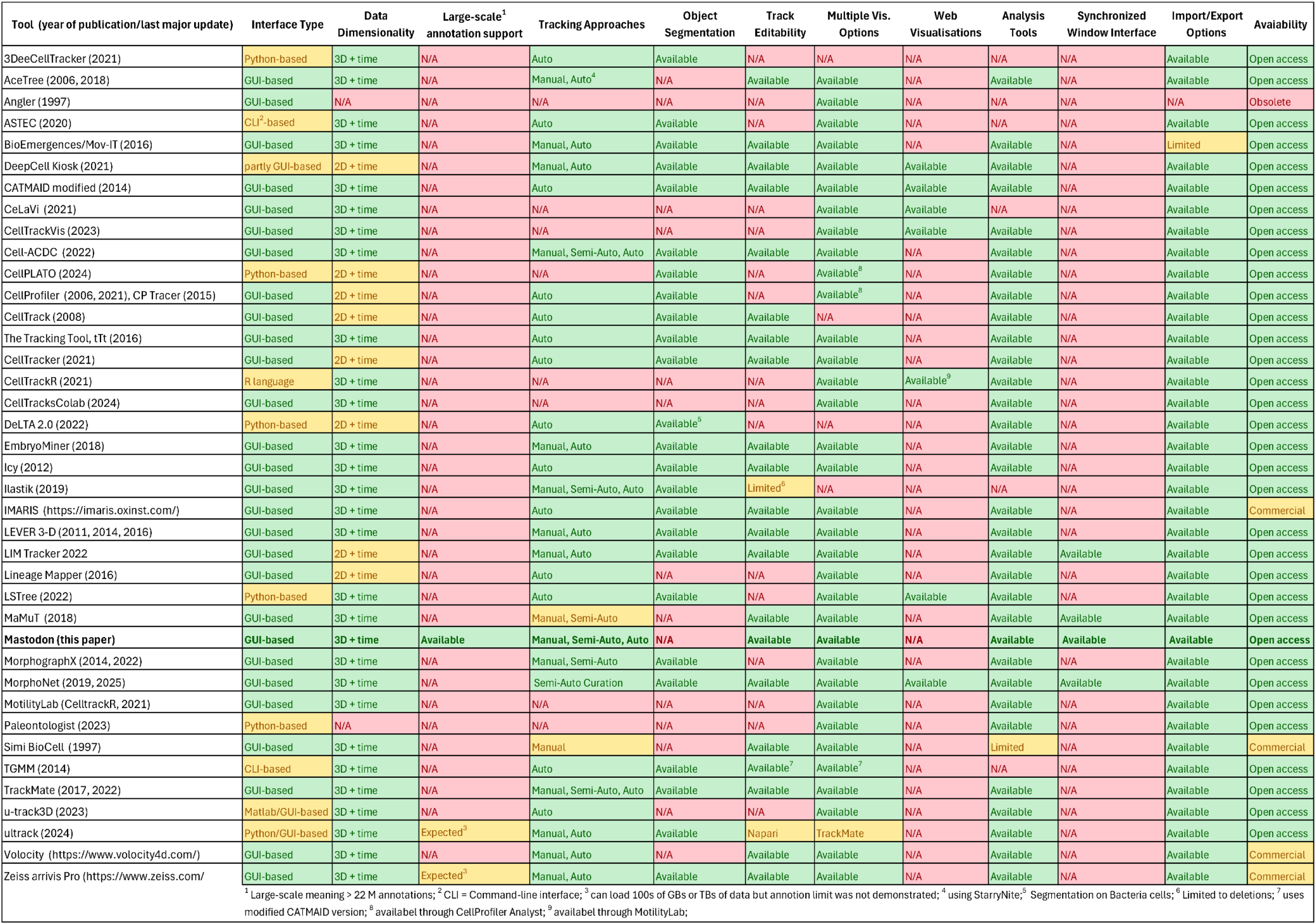
Feature comparison of lineage tracking software tools. Overview of 46 lineage-tracing software platforms, highlighting key capabilities such as data dimensionality, support for large-scale annotations (>22M), tracking approaches, object segmentation, track editability. Mastodon is compared alongside its ancestor MaMuT and other commonly used or emerging tools. Platforms vary widely in scope: some focus on segmentation and tracking algorithms without offering interactive GUIs, while others emphasize visualization or analysis. This comparison illustrates Mastodon’s positioning as a scalable, extensible solution for interactive lineage tracing.

**Supplementary Video 1. Manual, semi-automated, and automated tracking in Mastodon.**
Tracking can be performed manually or semi-automatically by selecting cells in the image data and tracing them over time. Mastodon provides advanced tools to link and edit spots and tracks, and to quickly define cell division events. When image quality permits, manual and semi-automated tracking can be supplemented with built-in automated tracking algorithms.

