## Supplementary Information for "Mastodon: the Command Center for Large-Scale Lineage-Tracing Microscopy Datasets"

#### Part I: Mastodon core technologies.

We developed a novel data model to represent, store and edit tracking data, able to support a large number of objects and offer a responsive user interface. We detail below several of the core components of the Mastodon code that help achieve these aims. When relevant, we compare the Mastodon data model to the one used in MaMuT [1] and TrackMate [2], based on the JGraphT [3] library. JGraphT is a powerful library to manipulate mathematical graphs that is instrumental to these two software. However, JGraphT was not developed with the aim of harnessing very large data.

##### 1. A Java end-user application to create, inspect, edit and analyze large lineages in Life-Science.

Mastodon aims to offer a scientific tool that enables end-users to automatically, manually or semi-automatically track cells in large images. It is a highly interactive application that allows inspecting and analyzing the resulting data and manually curating and editing it. The analysis of large images will result in large lineages, possibly built over millions of objects. So these features must be implemented in the context of very large data. When used in an application that aims at being interactive and responsive, they require special data structures and techniques that we detail below. While the languages of choice for high-performance computing are typically C or C++, we chose to use Java. This choice required building a special graph data structure, but allowed us to connect to the Fiji [4] ecosystem, facilitating the deployment of Mastodon and its use by the biologist community. Additionally, we could build upon the BigDataViewer framework [5] based on the ImgLib2 library [6], which allowed Mastodon to seamlessly interact with very large images, stored locally or remotely. In summary, the development of Mastodon has the following constraints:

- It is an end-user application that allows editing and inspecting the data. It must therefore offer a user interface that is interactive and responsive. We are therefore in need of a data structure that offers low latency.
- It must harness large data, in the sense that the lineages it will build will be made of numerous objects. The first target being 100 millions of objects. The data structure must therefore be compact and require little to no memory overhead per object.
- It must be extensible. Our experience with software like TrackMate [2] showed the usefulness of a software that can be reused or repurposed for other tasks beyond what they were initially meant for. This is particularly important in life sciences, where the number of developers is small, and redeveloping scientific software from scratch is very time consuming. We therefore turned to Java, an object-oriented language that

can handle complex code, is widely used and supported beyond the life sciences, and has proven invaluable for developing extensible ecosystems of scientific tools [4].

The next sections detail how these goals were achieved. They involve some basics in programming.

#### 2. A compact data model exploiting memory locality.

In programming, one of the key issues that limits the speed of iteration across a collection of objects is the time taken by transferring data from the computer memory to the CPU. CPU performance has improved tremendously in the past decades, by a factor of over 10,000 between 1980 and 2010. However in the same period of time, the performance of memory hardware only improved by a factor of 10 (see figure 2.2 in [7]). The data transfer from the RAM to the CPU is therefore likely to become the main bottleneck in responsiveness with current hardware. This is particularly unfortunate when iterating over a large collection of Java objects, using for instance the MaMuT and TrackMate JGraphT graph representing lineage data. The vertices of this graph are Java objects called 'Spots' and they represent cells. The Spot object is built on a key-value mapping that stores the name of several properties (such as 'POSITION_X') and their numerical values, all stored also as Java objects. Iterating through a large collection of Spots will trigger an even larger number of data transfers from RAM to CPU, amounting to significant slowdown of the user interface. Additionally, each of these individual objects contribute an overhead in memory space, to store the metadata required to manage them by the Java Runtime. This data structure is convenient and adequate when the number of vertices in the graph is not very large, typically below a few 100k of objects. But it will generate a significant slowdown and memory overhead for the use-cases we target with Mastodon.

A workaround involves limiting the number of data transfer events thanks to CPU cache, and designing a data structure that can exploit the cache. CPU cache is a high-speed memory component located between the CPU and the RAM. Without going too deep into details, the cache is used to store a copy of a large region in the RAM. When the CPU requests the content of a specific memory location in the RAM, a whole region around this location is copied to the CPU cache. If the next CPU instructions require data that is already in the cache, they are used directly, bypassing the need for a new, slow data transfer event. This will improve the speed of iteration, but only in cases where the data to iterate over is organized contiguously in memory. This is unfortunately not the case for the graph data structure based on JGraphT, where the data is split over objects spread all over the memory in a non-contiguous manner.

The graph data structure in Mastodon exploits memory locality. There are no single spot objects, but instead a very large primitive byte array that stores the data of all the spots in the lineage. The spot data, such as its position, shape (an ellipsoid represented by its covariance matrix), and edges in the graph it connects to, are laid out in a fixed-length part of the byte array (Supp. Fig. 1). The data of another spot is laid out just after in the array, resulting in the storage of a large collection of objects that is both local in memory and free of overhead. When the data of a spot is requested by the CPU, the data of several contiguous spots are copied on the cache. These data are likely to be used when iterating over the collection, skipping the need for several data transfer events and resulting in improved performance. This programming pattern, where a data structure is optimized to improve performance, is commonly used in applications that require low latency, such as video games [8]. When this pattern is used on linear arrays, one can expect a speedup of about 50 times in C++ [8]. With Mastodon, which uses this pattern but with a graph data structure in Java, we observe a speedup of about 30 times (main text). The code that allows designing and customizing such a data structure is shipped in the *mastodon-collection* library [9]. This library was built to be extensible and used in applications beyond tracking or life sciences.

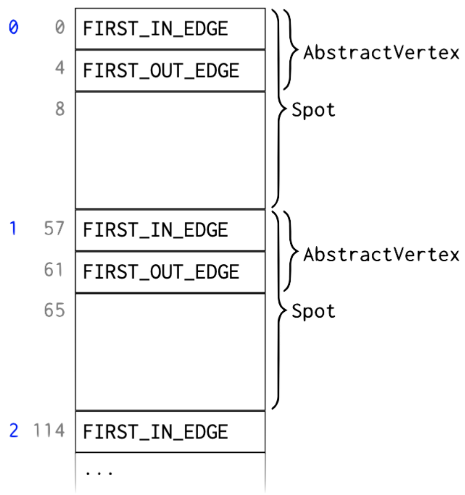


***Supp. Fig. 1**: Layout of a collection of spots in Mastodon. The data for the whole collection is stored in a single byte array. Gray numbers: index in the byte array. Blue numbers: index of spots in the collection. In this example the data size for a single spot is 57 bytes. The first two elements in the spot object are indices in the edge array (see below) pointing to the first incoming link to this spot and to its first outgoing link. They are specified in the mother class* AbstractVertex*, from which the* Spot *class is derived. These two indices are* int *numbers and occupy 4 bytes each. The rest of the 57 bytes are used to store the spot position, shape and time point.*

#### 3. Implementation within an object-oriented paradigm.

The data structure strategy presented above offers significant performance improvements but requires manipulating a byte array in a low-level manner to access single object properties. This can be cumbersome in an object-oriented language such as Java. To reconcile our approach with a Java application and facilitate reusing and extending Mastodon, we designed core Java classes that encapsulate the byte array and hide the low-level operations in core methods. The byte array encoding the information of a collection of objects is wrapped in a class deriving from the *Pool* mother class[1]. The instances of this class represent the large collection of objects we will manipulate. A pool can create proxy objects, deriving from the *PoolObject* class[2] and that represents the elements of the collection. In Mastodon, a *Spot* is a *PoolObject*, and has methods to access property values. When a property of a specific spot with index *i* in the collection is requested, the *Spot* object deserializes several bytes from the array, at a position computed from the spot index *i* and the offset of the property. For instance, taking the example of Supp. Fig. 1, to read the index value of to the first outgoing link (the second property) of the spot at index *i*, which is an *int*, the *Spot* proxy object will read the 4 bytes of the pool between positions $57 \times i + 4$ and $57 \times i + 7$ included, and deserialise them in an *int* (Supp. Fig. 2).

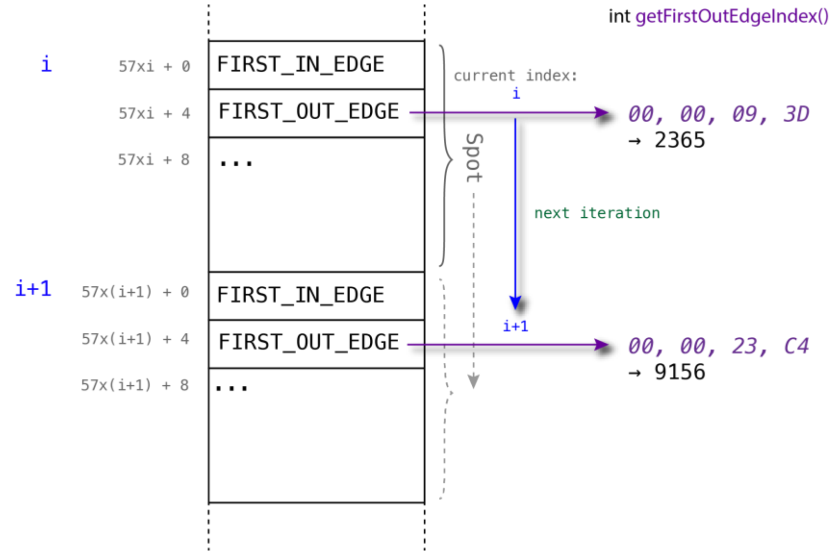


***Supp. Fig. 2****: Accessing property values in a Mastodon collection. The* Spot *class has a method called* getFirstOutEdgeIndex *that returns the index of the first outgoing edge. The* Spot *instance keeps track of the current index it is currently set at. When the value of the edge index is requested, the method reads the 4 bytes of the byte array between positions $57 \times i + 4$ and $57 \times i + 7$ included, and internally converts them to an* int, *which is returned by the method. When iterating, the same* Spot *instance just updates its current index* i *and the method will now return the edge index for the next spot in the collection.*

#### 4. Garbage-collection-free programming.

The *PoolObject* framework bridges the compact data model based on the pool array with object-oriented programming paradigms. We extended it further to limit ineffective object creation and garbage-collection events, improving further the performance of the Mastodon collection framework. Consider the following Java code, collecting the first outgoing edge indices of 1000 spots in a spot list:

[1] https://github.com/mastodon-sc/mastodon-collection/blob/master/src/main/java/org/mastodon/pool/Pool.java
[2] https://github.com/mastodon-sc/mastodon-collection/blob/master/src/main/java/org/mastodon/pool/PoolObject.java

```java
RefArrayList< Spot > pool = ...
int n = 1000;
int[] timepoints = new int[ n ];
for ( int i = 0; i < n; i++ )
{
	Spot spot = pool.get( i );
	int t = spot.getTimepoint();
	timepoints[ i ] = t;
}
```

The *RefArrayList* class wraps a *Pool* and behaves like a Java *List*.

In line 6 of the code, a *Spot* object is requested from the collection. As stated above, this will cause the *Pool* to create a new *Spot* instance and return it with its internal index value set to *i*. In the next line the *Spot* is used to deserialize the time-point value. The loop then iterates. This means that for each iteration, a new *Spot* instance is created, which triggers memory allocation. At the end of the iteration, the instance has no further use, and might be collected by the Java garbage-collector. Even with modern garbage-collector strategies, object creation and garbage collection events amount to significant delays when iterating large collections [10]. This will result in a downgraded performance, in particular when iterating over large collections (*n* large) where these undesired calls will be numerous.

To limit the impact of object creation and garbage collection, the Mastodon-collection framework offers an API that allows skipping them almost entirely. An improved version of the code above is the following:

```java
RefArrayList< Spot > pool = ...
int n = 1000;
int[] timepoints = new int[ n ];
Spot ref = pool.createRef();
for ( int i = 0; i < n; i++ )
{
	Spot spot = pool.get( i, ref );
	int t = spot.getTimepoint();
	timepoints[ i ] = t;
}
pool.releaseRef( ref );
```

At line 4, a proxy object is explicitly requested from the *Pool*:

```java
Spot ref = pool.createRef();
```

This proxy object is then reused within the loop to retrieve a specific spot at line 7:

```java
Spot spot = pool.get( i, ref );
```

Importantly, this same instance will be reused and returned in this call. The value of *spot* is equal to *ref*; these two variables point to the same instance in memory. The call *get( i, ref )* simply sets the internal value of the index *i* in the existing *Spot* instance, giving access to the desired spot properties, without creating or destroying a Java object. This effectively abolishes creating new objects at each iteration of the loop, and in turn, having them garbage-collected. At line 7, we call the *releaseRef* method on the proxy object:

```
pool.releaseRef( ref );
```

This call signals that the current code won't be using the proxy object anymore. The object is then returned to the pool, and made available to other places in the code, where the method *pool.createRef()* will be called. This strategy is used everywhere in the Mastodon application. It makes it possible to create a lineage model containing hundreds of millions of spots by only effectively creating less than a dozen Java objects and without triggering garbage collection.

#### 5. A graph data model based on Mastodon-collections.

The sections above describe strategies to build an optimized data structure to manage a large collection of objects. In Mastodon we need a mathematical graph data structure to represent lineages. In such a graph, the vertices are *Spot* objects that represent cells at a given time-point. The graph edges link spots across time-points, and are used to propagate a cell identity over time. When a spot in frame *t* links to a spot in frame *t+1* (when there is an edge in the graph that connects them), it means that they represent the same cell over time. When a cell divides, there is one spot for the mother cell in frame *t* that has two outgoing edges pointing to two spots in frame *t+1*, corresponding to the two daughter cells. The *mastodon-graph* library [11] provides such a data structure, based on the optimization presented above. As for the collection part, we built the graph data structure to be extensible and reusable beyond tracking. The graph data structure in Mastodon is built on two memory arrays, or pools, one for vertices and one for edges. We detail below the memory layout of these pools.

##### Vertex memory layout.

The first array is used to store vertices, or spots, that we introduced above (Supp. Fig. 1). The fixed *AbstractVertex* part comprises element indices *FIRST_IN_EDGE* and *FIRST_OUT_EDGE*, occupying 4 bytes each. The remaining 49 bytes are *Spot* attributes (position, shape, time-point, ...).

- *FIRST_IN_EDGE* is the element index (in the edge memory array) of the first incoming edge, i.e., an edge pointing to this vertex. The remaining incoming edges of the same vertex are stored as a linked list in the edge memory as described below. If this vertex does not have any incoming edges, *FIRST_IN_EDGE* is -1.
- Similarly, *FIRST_OUT_EDGE* is the element index of the first outgoing edge, i.e., an edge starting from this vertex. The remaining outgoing edges of the same vertex are

stored as a linked list in the edge memory as described below. If this vertex does not have any outgoing edges, *FIRST_OUT_EDGE* is -1.

##### Edge memory layout.

The second pool stores the graph edges, or links. In the example displayed in Supp. Fig. 3, each element of the *AbstractEdge* subclass *Edge* requires 24 bytes to store. The data for the $i^{th}$ Edge starts at byte *i x 24*. The fixed *AbstractEdge* part comprises element indices *SOURCE*, *TARGET*, *NEXT_SOURCE_EDGE*, and *NEXT_TARGET_EDGE*, occupying 4 bytes each. The remaining 8 bytes are other *Edge* attributes.

- *SOURCE* is the element index (in the vertex memory array) of the vertex from which this edge starts.
- *TARGET* is the element index (in the vertex memory array) of the vertex to which this edge points.
- *NEXT_SOURCE_EDGE* is the element index (in the edge memory array) of the next outgoing edge of the source vertex, , the next edge that has the same *SOURCE*. If there is no such edge then *NEXT_SOURCE_EDGE* is -1.
- *NEXT_TARGET_EDGE* is the element index (in the edge memory array) of the next incoming edge of the target vertex, , the next edge that has the same *TARGET*. If there is no such edge then *NEXT_TARGET_EDGE* is -1.

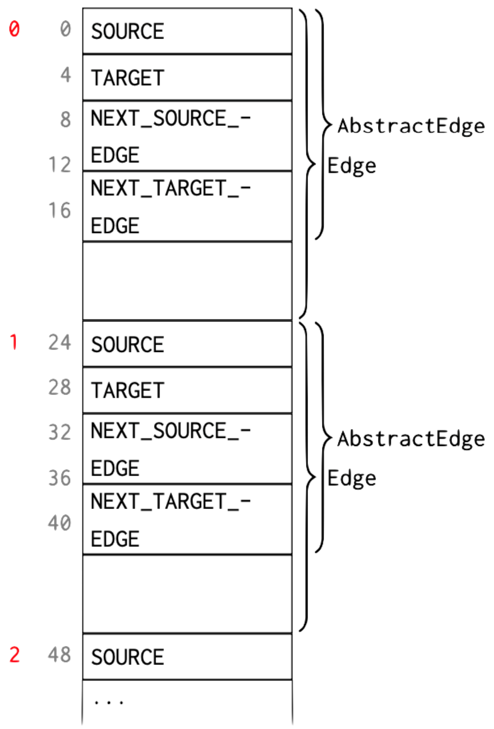


***Supp. Fig. 3**: The graph edge data structure. In the left-most column, element indices are shown in red, followed by byte indices in gray.*

**Example.**

Consider the following example graph comprising vertices *A*, *B*, *C*, *D*, *E* and edges *a*, *b*, *c*, *d* (Supp. Fig. 4).

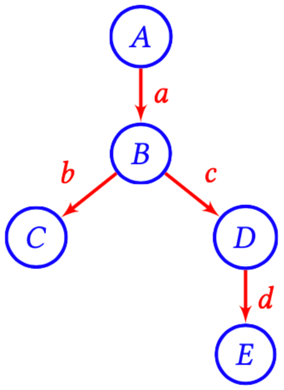


***Supp. Fig. 4***: *Example graph.*

This is laid out in memory as follows (Supp. Fig. 5):

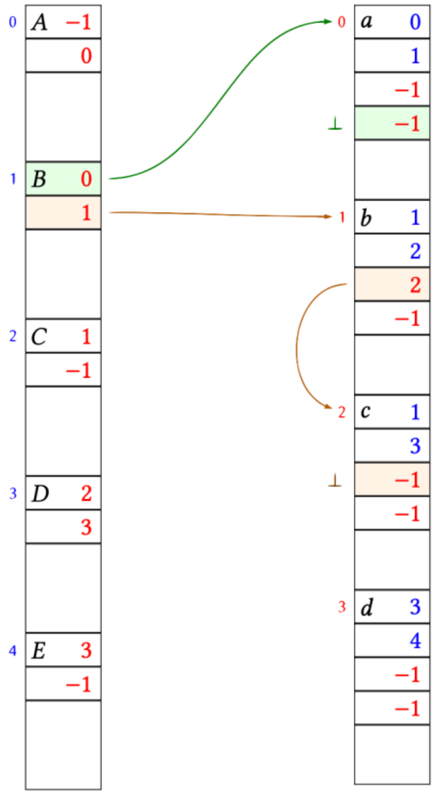

***Supp. Fig. 5***: *Memory layout for the example graph of Supp. Fig. 4. Left: vertex pool. Right: edge pool. The links between vertex B and its adjacent edges a, b, c have been highlighted.*

Let us look at that in more detail: Vertex *B* is stored at element index 1 in the vertex memory array. *B* has one incoming edge *a*. The edge *a* is stored at element index 0 in the edge memory array. Therefore the *FIRST_IN_EDGE* field of *B* is 0. Apart from *a*, the vertex *B* has no further incoming edges. Therefore, the *NEXT_TARGET_EDGE* field of a is -1, i.e., the list of edges entering *B* terminates here. *B* has two outgoing edges *b*, *c*. The edge *b* is stored at element index 1 in the edge memory array. Therefore the *FIRST_OUT_EDGE* field of *B* is 1. The next outgoing edge of *B* is *c* which is stored at element index 2. Therefore, the *NEXT_SOURCE_EDGE* field of b is 2. After *c*, the vertex *B* has no further outgoing edges. Therefore, the *NEXT_SOURCE_EDGE* field of c is -1, i.e., the list of edges leaving *B* terminates here.

Below (Supp. Fig. 6) is the same memory layout again, this time highlighting the references from edges *a*, *b*, *c* back to the vertex memory array. For example, edge *c* is leaving vertex *B* (index 1) and entering vertex *D* (index 3). Therefore the *SOURCE* field of *c* is 1, and the *TARGET* field of *c* is 3.

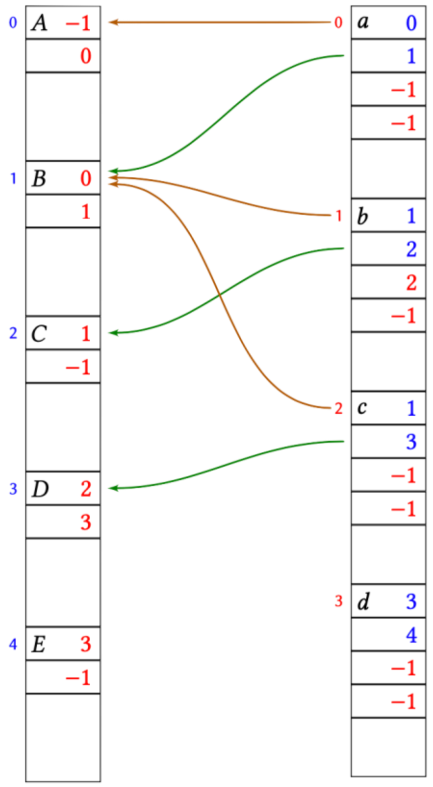


***Supp. Fig. 6**: Memory layout for the example graph of Supp. Fig. 4, this time highlighting the source vertices (ocher arrows) and target vertices (green arrows) for edges* a*,* b *and* c.

#### Free-list of unallocated elements.

The vertex and edge memory arrays can only ever grow. When elements are released, they are simply marked as free for reuse. Assume that in the above example vertex *D* is deleted, as well as its adjacent edges *c*, *d*, leaving the graph as follow (Supp. Fig. 7):

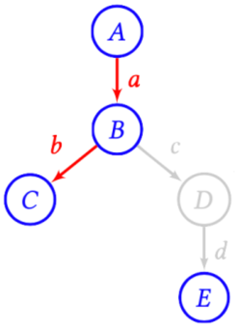


***Supp. Fig. 7****: Example graph after the deletion of vertex* D*. The edges* c *and* d *have been deleted in the process.*

After removing *c*, *d*, *D* the memory layout looks like this (Supp. Fig. 8):

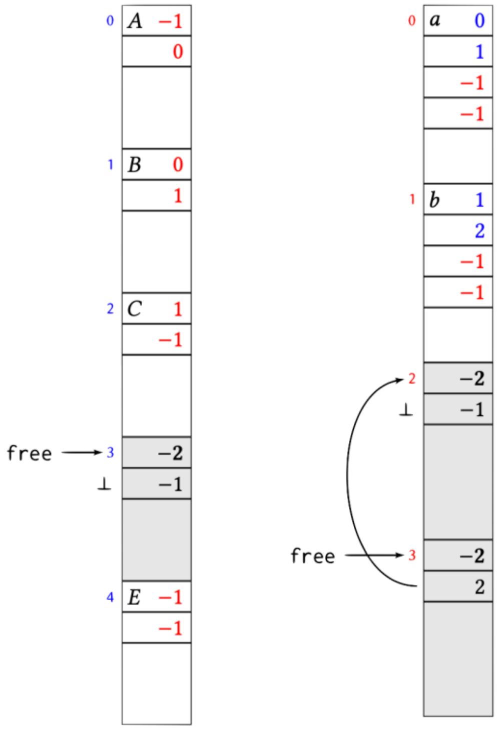


***Supp. Fig. 8****: Memory layout after the deletion of vertex* D*, corresponding to the graph of Supp. Fig. 7. Each pool stores the index* free *that is the head of the free list.*

The element 3 in the vertex memory array as well as elements 2 and 3 in the edge memory array have been marked as free. This is done by putting the magic number “-2” into the first 4 bytes of the element. In vertices and edges the first 4 bytes are always occupied by a (positive) index or a “-1” index list terminator. Therefore, occupied and free blocks can not be confused. The next 4 bytes of a freed element are the index of the next freed element in the (same) memory array, or -1 if there is no next freed element. Each memory array remembers the index *free* of the first freed element. Newly freed elements are enqueued at *free*, that is, at the head of the free-list. So in the above example, edge element 2 was freed first, followed by edge element 3.

The next edge element will be allocated at the head of the free-list and the *free* index moves to the next element. For example, assume that a new edge is created from B to E (Supp. Fig. 9):

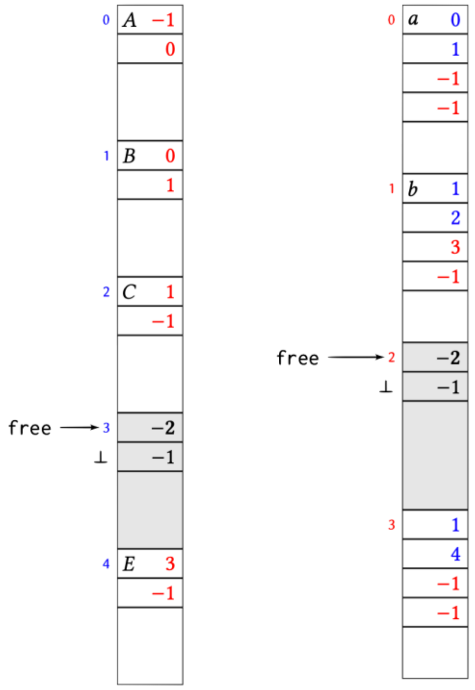


***Supp. Fig. 9****: Memory layout after creating an edge from* B *to* E.

If *free* is -1, then no more freed elements are available and the underlying memory array has to grow to fit newly added elements.

#### 6. Adaptive level of detail in the lineage display.

The display of a full lineage in a time-hierarchy view, as in the TrackScheme view (Supp. Fig. 10), can be problematic for large lineages. When the lineage is fully zoomed-out, it requires painting all the vertices and edges of the lineage. This would be painfully slow, and would break the application for large lineages. For instance, the MaMuT lineage display (also called TrackScheme) fails to draw a model made of 2.8 millions of spots and crashes. We therefore rewrote the TrackScheme view in Mastodon entirely, using a level-of-detail strategy to accommodate displaying a large lineage. The painting of lineage view works as follows.

First, a layout operation determines all the X and Y position of the nodes - the objects corresponding to spots in TrackScheme, in a global coordinate system. The layout Y coordinates of a node is simply given by its time-point. The nodes X coordinates are determined by iterating through tracks. We start from a list of roots (nodes without a parent node, the beginning of a track), and recursively descend to leaf nodes. The layout operation then assigns X coordinates such that:

- Leaf nodes are assigned a X layout position of 0, 1, 2, etc. based on which was reached first.
- Non-leaf nodes are centered between the first and last child's X layout position.

This yields the coordinates of all the nodes in a global coordinate system.

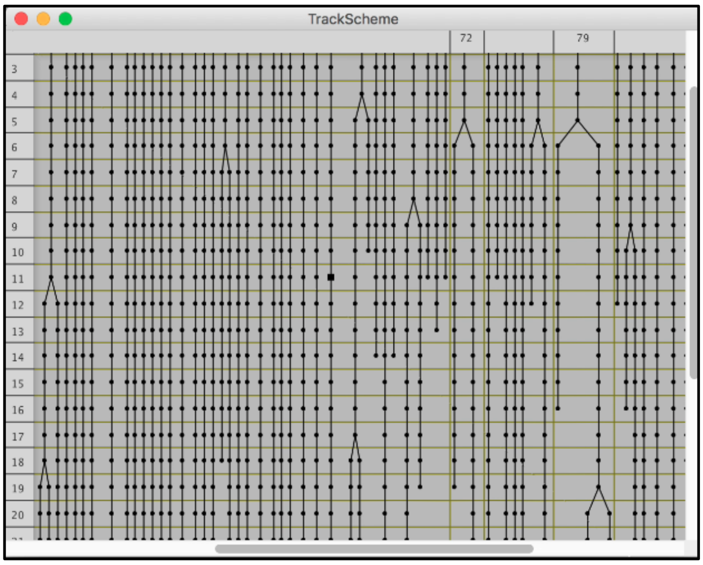


***Supp. Fig. 10****: Example lineage view in TrackScheme. The cells are represented by nodes, one per cell and per time-point. There is one row per time-point. The nodes are arranged vertically along time from top to bottom. A cell division is represented by a fork: one node representing the mother cell that connects to two nodes in the row below. One cell lineage is represented by a column in this view, and the columns are sorted from left to right by the name of the first cell in the lineage.*

In a second step, screen entities are created, based on the node layout coordinates and the portion of the lineage view to paint. The nodes that are present in the current view are iterated, and their coordinates on screen are computed. When they are spaced enough so that they can be individualized on screen, a screen entity is created for a node, with a level of detail that depends on the actual spacing between them. When the screen position of a node is closer to its neighbor by less than 1 pixel, separate screen entities are not created for them. Instead, they are merged into a less detailed entity representing a wide range of nodes. This yields the different node representations that can be observed when progressively zooming out or resizing the window, as exemplified in Supp. Fig. 11.

Finally, all the created screen-entities are actually painted in the TrackScheme window. Thanks to this level-of-detail approach, there can be considerably fewer screen entities to paint than there are nodes in the lineage. When they are individualized, their corresponding screen entity has more or less content to paint, depending on the proximity to their neighbor, hence their number on the screen. Finally, the drawing uses a buffer strategy to improve latency even more: screen entities are drawn (step 3) while the position of the nodes in the next iteration (e.g. after resizing the window) are computed (step 2).

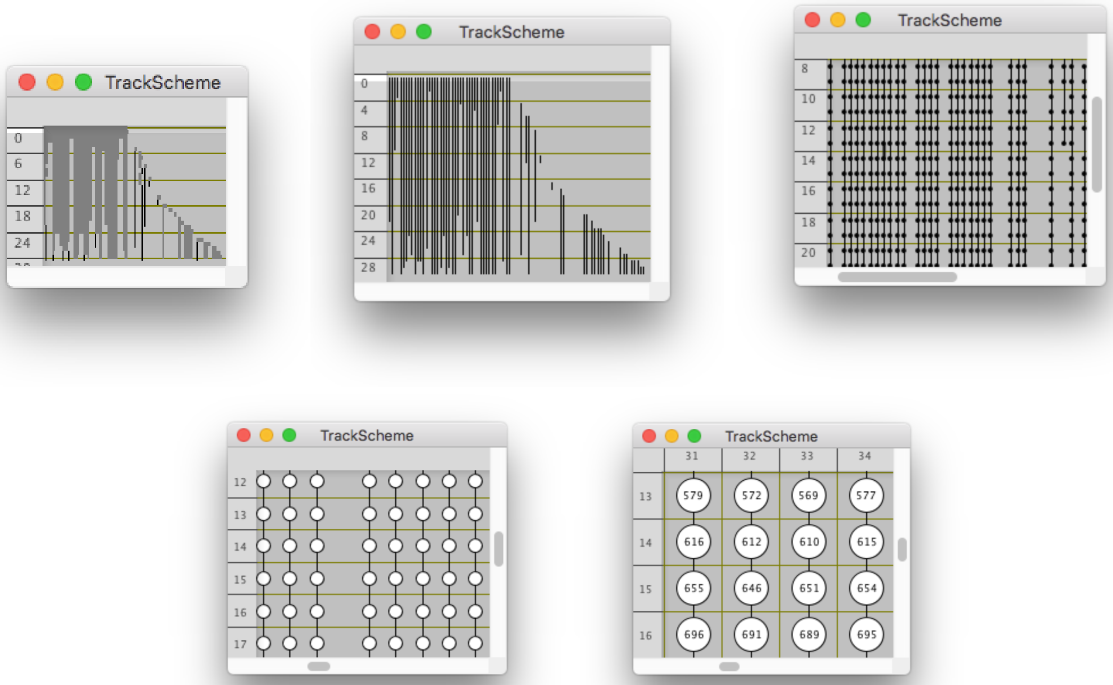


***Supp. Fig. 11***: *TrackScheme displays nodes differently depending on the zoom level. From left to right then top to bottom: 1. At low zoom level, nodes that coalesce are shown as gray boxes. 2. Zooming in, the individual tracks appear first as black lines. 3. When they are separated enough, nodes appear as black dots. 4. With an even higher zoom level they are shown as empty circles. 5. Until they grow big enough so that the node labels can be painted.*

#### 7. Efficient object retrieval with kD-tree search in convex polytopes.

Several important features of Mastodon rely on the fast retrieval of data items (spots and links) close to specific 3D positions. For instance we need to do so when the user moves the mouse close to the drawing of a spot in a view, to retrieve the spot in question. Or to determine what spots must be painted in an image view, depending on the zoom level, position, rotation and field of view. There exist several well established algorithms and techniques for fast object retrieval for spatial queries, where the space in which we need to retrieve data items is a 3D rectangle aligned with the X, Y, Z axes of the dataset. In our case it is not. In Mastodon, the image view is a BigdataViewer (BDV) window. This view can be rotated by an arbitrary angle around an arbitrary axis. The XY view plane does not match an orthogonal plane of the dataset. Also, the planes that make the bounds of the field of view are not aligned with these axes. In our case, the field of view is a convex polytope. It is a portion of 3D space delimited by a set of planes that are not necessarily orthogonal. Such a space is similar to an ideal diamond, in which each facet would be one of the bounding planes. The interior of this diamond is a convex polytope. Our goal is to know what are the points that are inside this volume so that we can paint them without losing time painting the ones not in the field of view.

At the time of the development of Mastodon, there was no published algorithm for the fast retrieval of points in a convex polytope. One of us (Tobias Pietzsch) derived such an algorithm in 2016, and it is detailed in the document linked below[1]. To the best of our knowledge this is unpublished.

[1] https://mastodon.readthedocs.io/en/latest/_downloads/9e43cd9b551c1991519dddbad738ed71/TPietzschConvexPolytopes.pdf

### Part II: Practical Implementation: Usage, Integration, and Performance

#### 1. Reference system hardware specifications

The reference notebook used for benchmarking Mastodon and generating Blender visualizations has the following hardware specifications:

| Notebook model | ThinkPad X1 Carbon Gen 9 |
|---|---|
| Processor | 11th Gen Intel(R) Core(TM) i7-1165G7 @ 2.80GHz 2.80 GHz |
| Installed RAM | 32,0 GB |
| System type | 64-bit operating system, x64-based processor |
| Operating System | Windows 11 Pro (Version 24H2) |

***Table 1:*** Hardware specifications of the reference notebook

#### 2. Flexible import/export architecture

Upon launching Mastodon, the launcher window offers options to either create a new project from BDV files (XML paired with N5, OME-Zarr, or HDF5), or to directly import tracks generated with Simi-BioCell, TGMM, TrackMate, or MaMuT. We also made a user-friendly CSV importer available that doesn't require columns in any particular order and is capable of taking already linked spots into account. Other import add-ons allow the import of spots from segmented label images and include the GraphML Importer. Mastodon also supports exporting a variety of data formats.

##### *Summary of available Importers in Mastodon*

| Importer | Input Files | Access/Location | Notes / References |
|---|---|---|---|
| Simi BioCell to MaMuT | BigDataViewer file + Simi-BioCell file | Mastodon launcher window | Published in Dev Genes Evol 229, 137–145 (2019). |
| TGMM | BigDataViewer file + TGMM folder | Mastodon launcher window | Mastodon online documentation, part C |
| TrackMate/MaMuT | XML file | Mastodon launcher window | — |
| CSV Importer | CSV file with columns (X, Y, Z, Frame); optional: Quality, Radius, Label, ID, Parent ID, Tag | Plugins > Imports > CSV Importer | Mastodon online documentation, part C |
| Import from Segmented Labels | Images/BigDataViewer files opened in Fiji | Plugins > Imports > Import Spots from Label Image | Mastodon online documentation, part C |

| | | | |
|---|---|---|---|
| GraphML Importer | GraphML file in a specific format | Mastodon online documentation, part C; File > Import > Import GraphML | Mastodon online documentation, part C; Github<br>Imports spot and track data from a properly formatted GraphML file into Mastodon |
| Cell Tracking Challenge Importer | CTC format | Requires Fiji update site:https://sites.imagej.net/Ulman/; Plugins > Import from CTC format | https://celltrackingchallenge.net/<br>Imports challenge-format tracking/ground-truth data into Mastodon<br>Maška et al., 2023 |
| OME-NGFF | — | Mastodon launcher window | Github, Moore J. et al., 2021 |

***Table 2 - List of available Importers in Mastodon***

#### *List of available Exporters in Mastodon*

| **Exporter** | **Output** | **Access/Location** | **Notes / References** |
|---|---|---|---|
| Label Image Export | Segmentation mask images | File > Export > Label image using ellipsoids | Mastodon online documentation, part C |
| GraphML Export | GraphML file representing track structure (branches) | File > Export > Export to GraphML (branches) | Mastodon online documentation, part C<br>Brandes et al., 2002 |
| Export Measurements | CSV file with spot counts and division counts per time point | File > Export > Export measurements | Mastodon online documentation, part C |
| PhyloXML Export (selected spots) | Subtree of the selected spot in phyloXML format | File > Export > Export phyloXML for selected spot | Mastodon online documentation, part C; Han and Zmasek 2009 |
| MaMuT Export | XML file for compatibility with the MaMuT tracking tool | File > Export > Export to MaMut file | — |
| CSV Export | Table-formatted CSV data (exact output not detailed) | File > Export > Export to CSV (from Table or Selection Table window) | — |
| Cell Tracking Challenge Exporter | Converts lineage into challenge-format tracking/ground-truth data, with spots shown as user-sized boxes or spheres in label masks. | Requires Fiji update site: https://sites.imagej.net/Ulman/; Plugins > Export to CTC format | https://celltrackingchallenge.net<br>Maška et al., 2023 |

***Table 3 - List of available Exporters in Mastodon***

#### 3. Integration with Blender via the Mastodon bridge

Blender is a free and open-source 3D graphics software (https://blender.org) with a wide variety of use cases: 3D modeling, texturing, data visualization, animation and rendering. It also has a history of scientific usage and offers numerous features, tutorials, and community resources. It exposes a Python API, which we utilize to import tracks from Mastodon, stored in a CSV file, and convert it into Blender's native 3D geometry to take advantage of its rendering capabilities.

##### Installation

We have implemented a user-friendly installation guide within Mastodon, enabling users to apply Blender's 3D capabilities with just a few clicks (see part A of our online documentation under Interactive 3D viewer for tracking data (Blender View)). The installer can be accessed from any Mastodon window via **Window > Blender Views > Setup Blender Addon** (Supp. Fig. 12).

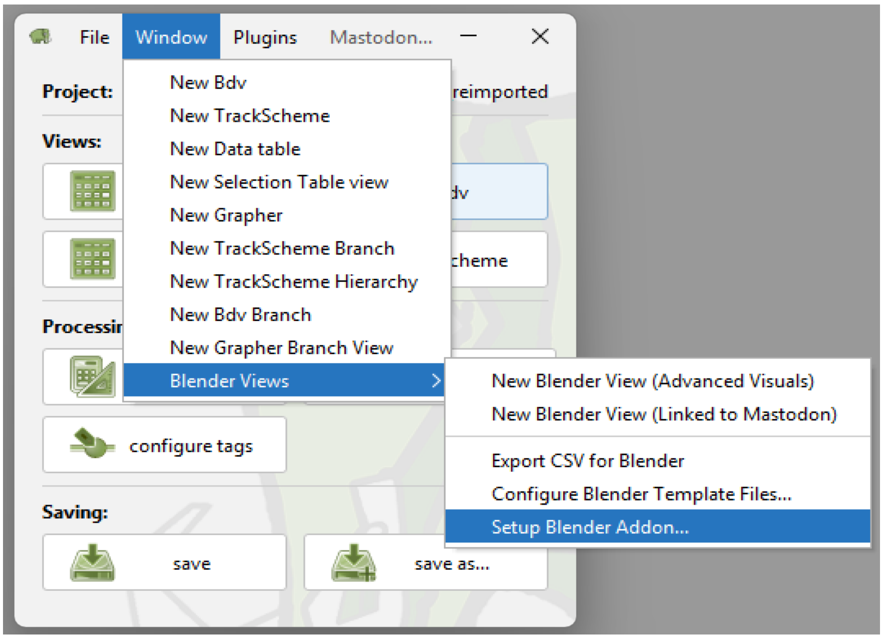


***Supp. Fig. 12:*** Accessing the 'Setup Blender Addon' dialog

##### Blender integration into Mastodon

In Blender each spot is represented by a 3D vertex, and linked spots are connected by mesh edges. Additional attributes like spot radii and tag colors are stored alongside the spot's position and time point. We apply a Geometry Nodes modifier to this 3D mesh to convert it into a visually appealing representation of spots as spheroids and links as tubular shapes with tapered tails. Geometry Nodes are a powerful node-based system for programmatically manipulating geometry in Blender by connecting individual function blocks into a node graph. The corresponding processing pipeline is illustrated in **Supp. Fig. 13**. Similarly, we rely on Blender's Shader Nodes system to procedurally generate surface properties, i.e. color and roughness, from the tag attributes stored in the original mesh.

Geometry Nodes allow us to conveniently define custom parameters and expose them to the node graph, letting the user tweak the look of the 3D representation in real time. These parameters include the overall cell scale, the range and thickness of the tracks, as well as performance-related parameters like the resolution of the mesh. Lowering the resolution is especially useful for visualizing large data sets, where fine cell details are of less importance than performance. The pipeline is divided into two processing branches, one for spheres and one for tracks. Each branch is encapsulated into a subgroup in the graph for a better node organization.

As a first step in creating the spheres, we convert the vertices of the original mesh into point objects. Blender differentiates between vertices and points, as the latter ones can also directly be rendered as point clouds on the GPU. After the conversion, we delete all points whose time point attributes do not match that of the current frame. Remember that Geometry Nodes act as a procedural modifier, so none of the underlying geometry is actually being deleted. We then instance icospheres on the remaining points. These types of spheres are geodesic polyhedrons and consist of regular triangles. Blender allows easy adjustment of the number of subdivisions as part of the icosphere primitive node. Each icosphere instance is then scaled according to the radius attribute stored in the underlying point. During the instancing process, the same mesh is being rendered onto the screen by the GPU multiple times, while only a single mesh resides in GPU memory. However, to achieve a more organic and inhomogeneous look of the cells, we need to be able to displace each mesh differently. We thus turn all icosphere instances into real geometry to deform every cell separately, using a 3D perlin noise texture for surface displacement. The resulting geometry is then merged with the result of the track representation.

To construct the tracks, we first discard all segments outside the user-defined visibility range, defined as a sliding window between the visible number of frames before and after the current time point. The remaining mesh is converted into a curve. Curves are spline types, and per default they are interpolated linearly between adjacent points. We provide an optional toggle to change the interpolation mode to NURBS (Non-Uniform Rational B-Splines), which smooths out the cell trajectories for a more natural look without jagged corners. Each anchor point along the curve possesses a radius attribute, which we set to follow a function that takes the normalized position on the curve as input, and returns the radius at this position to taper off the tail towards the end. We then use a curve-to-mesh operator that converts all the curves back into 3D geometry. The operator works by sliding a profile curve, in our case a circle of low resolution, along the curve to create a 3D surface.

After merging both the spheroids and the tracks together, we apply a procedural material that determines the surface properties of the geometry. The base color depends on the tag color stored in each spot. We mix in a perlin noise texture to create subtle differences in brightness. As an artistic choice, we also layer an ambient occlusion pass over the base color. This darkens the cavities between cells and emphasizes the 3D structure of the geometry.

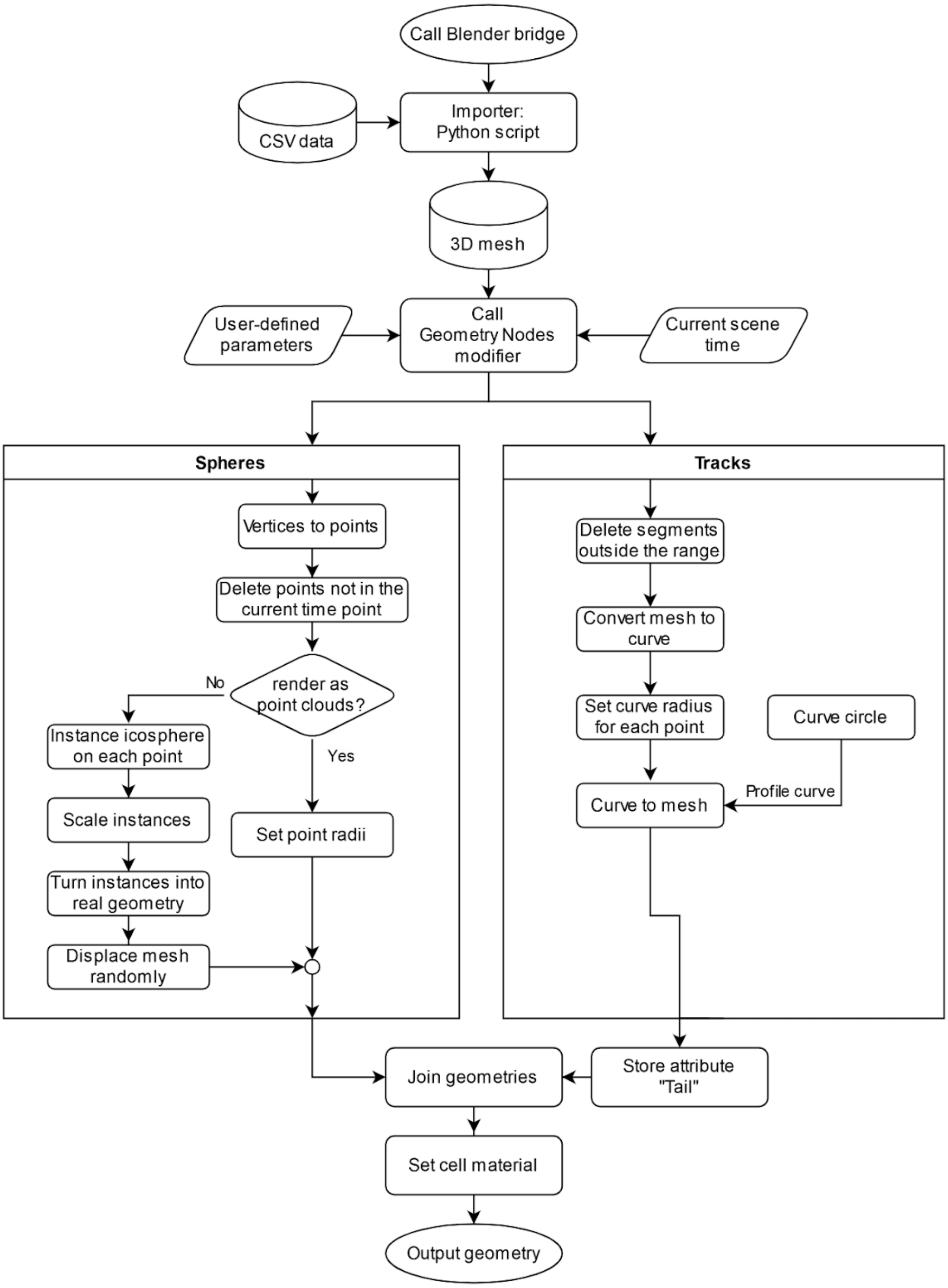


**Supp. Fig. 13:** Workflow for importing and rendering CSV Data in Blender via the Mastodon Blender Bridge. This flowchart illustrates the process of importing CSV data into Blender using a Python script and converting it into 3D geometry via the Geometry Nodes modifier. The workflow is split into two branches: Spheres and Tracks. In the Spheres branch, vertices are converted into points, and either point clouds or icosphere instances are generated, scaled, and turned into real geometry. In the Tracks branch, the mesh data are converted into curves, with radius adjustments and curve profiles applied. Both branches eventually merge, where geometries are joined, a cell material is assigned, and the final output geometry is produced.

Another subtle perlin noise creates fluctuations in the surface roughness of the cells, which can be seen around the specular highlights. The 3D scene is lit using several white area lights that create smooth shadows. One can animate the camera to slowly revolve around the sample, making it easy for the user to render out a complete animation without having to set up or manipulate the render scene.

We implemented two methods for exporting Mastodon projects into Blender that use the same CSV import-export approach but differ slightly in functionality:

##### *Blender View (Linked to Mastodon):*

This method automatically exports an open Mastodon project into Blender, with each tracking branch represented as an individual object. A major advantage is that individual spots can be selected at any time point within Blender, and selections are automatically synchronized with the TrackScheme/BDV windows, and vice versa. This is the recommended approach for quickly exploring and interacting with tracking data in 3D.

- Available under: **Window > Blender Views > New Blender View (Linked to Mastodon)**

Upon launching, Blender View includes an option to set a time scaling factor (Supp. Fig. 14), which interpolates between time steps to create a smoother flow of moving spots over time.

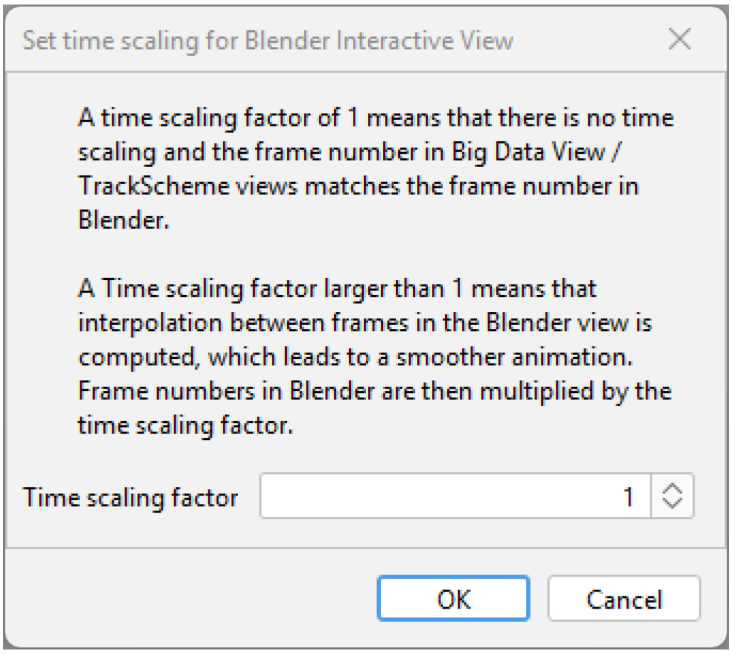


***Supp. Fig. 14 –*** Setting a time scaling factor for a new Blender View window (Linked to Mastodon).

Once the Blender window opens, Mastodon's track data still needs to be imported and converted into a 3D geometry with spots being represented as vertices. This requires some time before the Blender window becomes responsive, depending on the annotation size.

In the upper right corner of the 3D viewport area, below the ‘Options’ menu, a small extension arrow can be pressed (Supp. Fig. 15, encircled in pink), which expands the Mastodon 3D View Settings box (Supp. Fig. 15, framed in pink). Within this box, the appropriate Synchronization Group (Supp. Fig. 15, encircled in orange) corresponding to the actively linked Mastodon windows can be selected (Supp. Fig. 15, indicated by the small lock symbol shown in the BDV window inset at the bottom right of the figure). Additionally, the sphere size can be adjusted, and one of Mastodon’s available Tag Sets can be chosen (Supp. Fig. 15, orange arrows).

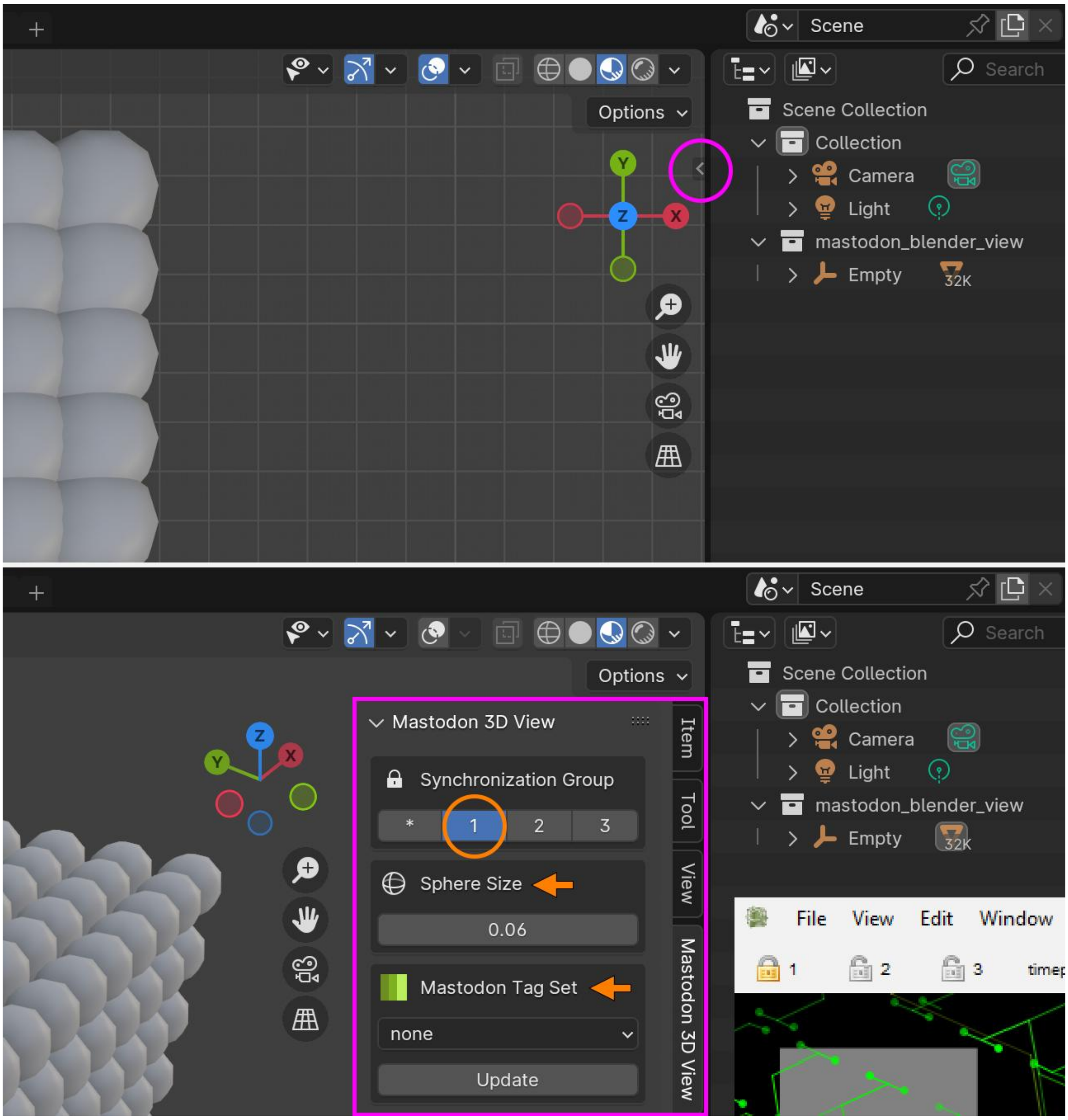


***Supp. Fig. 15:*** Accessing the Mastodon 3D View Settings. Using a small extension arrow in the upper right corner of the 3D viewport (below the ‘Options’ menu, encircled in pink) can be pressed to expand the Mastodon 3D View Settings box (framed in pink). Within this box, the appropriate Synchronization Group for the actively linked Mastodon windows can be selected (small lock symbol as seen in the BDV window inset at the bottom right). Sphere size adjustments and Tag Set selection (orange arrows) are also available. Changes to Tag Sets or tag colours require updating via the ‘Update’ button. Modifications to tracks (e.g., deletions or annotations) necessitate launching a new Blender View window, as real-time updates are not supported.

If a new Tag Set is created or tag colours are modified, synchronization with the changes can be achieved by pressing the 'Update' button. However, updates to edited tracks, such as small deletions or added annotations, cannot be performed on the fly. Such modifications to the TrackScheme structure always require the launching of a new Blender View window. For users with basic Blender experience, spheres (spots) can be modified by toggling from Object Mode to Edit Mode in the upper left corner of the 3D Viewport, followed by operations such as subdividing and smoothing.

A rendered image of the current 3D viewport view can be obtained using the '**View > Viewport Render Image**' command located in the upper left corner of the 3D viewport (Supp. Fig. 16, framed in pink). The resulting image can then be saved. To render multiple frames, Blender's Timeline editor can be used to set a Start and End frame (Supp. Fig. 16, framed in pink), and these frames can be recorded by executing the '**View > Viewport Render Animation'** command. Note that Blender's Output Properties (Supp. Fig. 16, encircled in pink) can be adjusted to change file save location (default: /tmp), resolution and output format (default: PNG), to name only a few options.

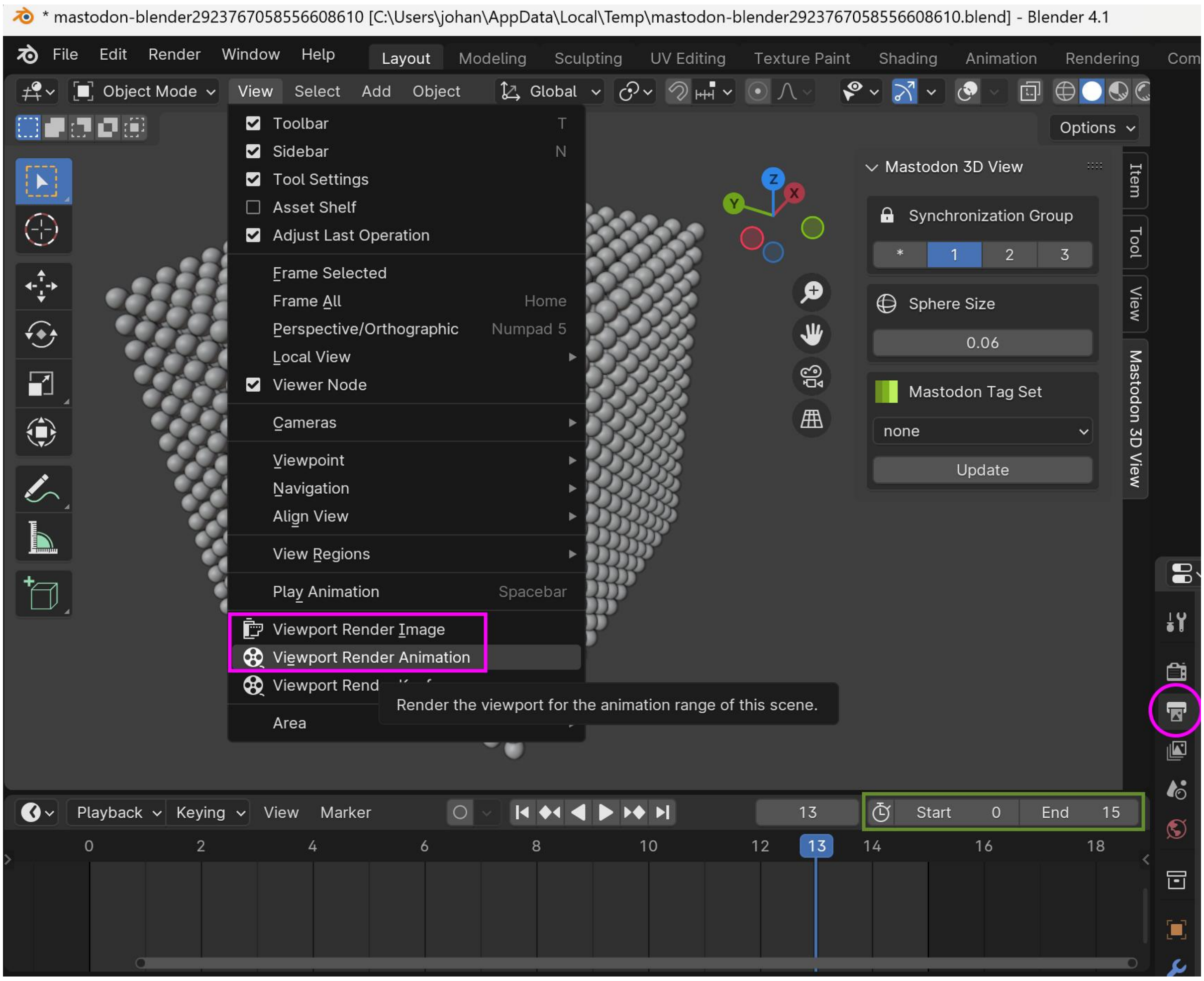

***Supp. Fig. 16:*** Setting up Blender's Timeline editor before capturing a series of frames from the 3D Viewport. Blender's default output properties (menu access encircled in pink) can be adjusted to change for example the file save location and the output format.

Blender View (Advanced Visuals):

Alternatively, an entire Mastodon project, including all tracking data, can be exported into Blender as a single object. While this limits direct interaction with individual spots, it significantly boosts performance, allowing the loading of millions of spots into Blender. Moreover, this method enables users to take full advantage of Blender's features, especially with our preset Geometry Nodes, to generate high-quality 3D models of their tracking data, with spots represented as spheroids and links as tapered tubes.

- Available under: **Window > Blender Views > New Blender View (Advanced Visuals)**

By default, a custom Geometry Nodes modifier is automatically applied to the exported data. This modifier visualizes spots and links as spheres and tracks, respectively, with adjustable sizes and lengths. The Modifier Properties panel (see Supp. Fig. 17) is located in the Properties Editor and highlighted with a pink circle and can be expanded by clicking it. This panel (pink frame in Supp. Fig. 17 allows users to quickly adjust spheres and tracks, which represent spots (vertices) and links (edges) in Mastodon, allowing users to change subdivisions of spheres, toggle track visibility, and modify track width as well as the number of frames shown before and after the current time point. Note that in this visualization, spot radii are also accurately represented through sphere sizes.

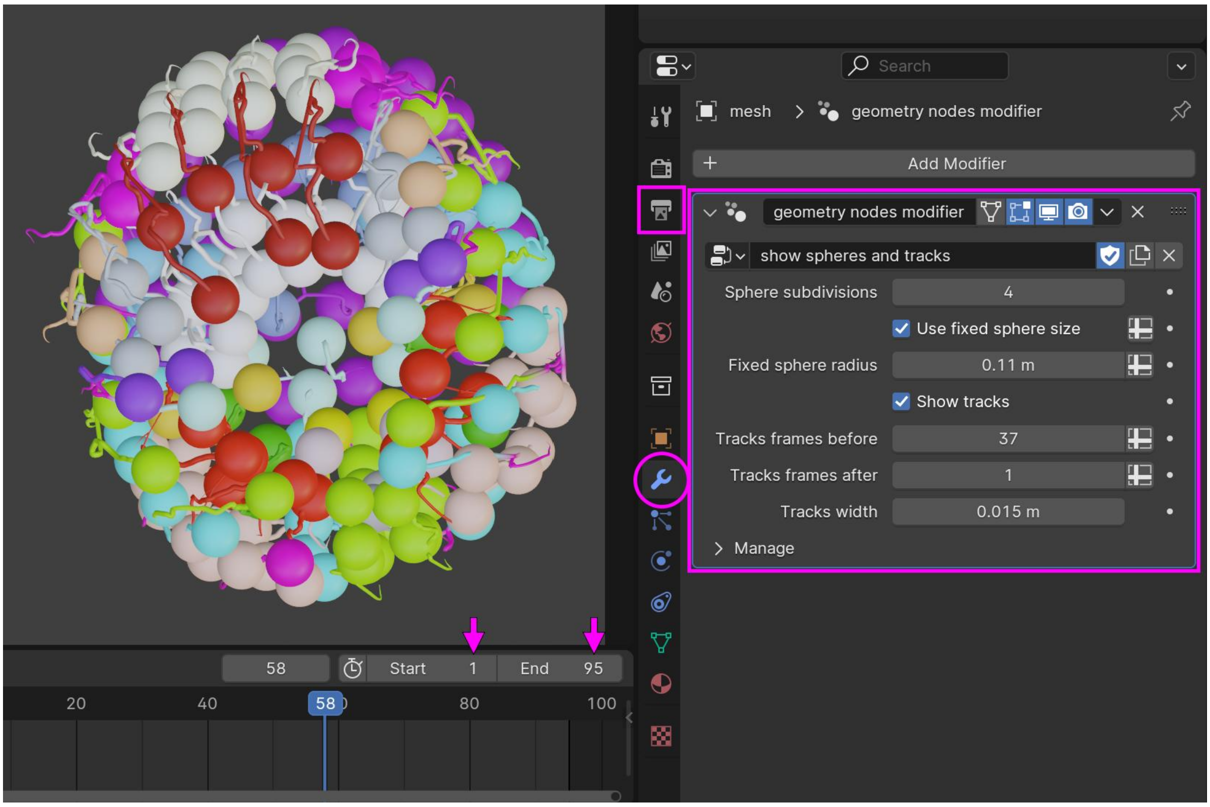


***Supp. Fig. 17:*** Accessing the Mastodon 3D View Settings. The Modifier Properties panel (located in the Properties Editor and highlighted with a pink circle) can be expanded by clicking it. This panel (see pink frame) allows adjusting the sphere size and subdivisions of spots, toggle track visibility, and the modification of track width as well as the number of frames shown before and after the current time point.

As mentioned previously, to take a snapshot of the tracking data displayed in Blender's active viewport (displaying the tracking data), users can navigate to **View > Viewport Render Image**. To capture an animation of the tracking data, they can go to **View > Viewport Render Animation (cf. Supp. Fig. 16)**. For the latter, we recommend defining the output file format, resolution, frame range (also highlighted with pink arrows in Supp. Fig. 17) and an output directory by clicking the Output Properties tab (marked with a pink square in Supp. Fig. 17).

##### Enhancing visualizations and enabling true rendering:

To further enhance visualizations and enable full 3D rendering via Blender's render engines, users typically need to configure camera settings, adjust lighting, and optionally modify the provided Geometry Nodes modifier (to manipulate textures, sphere shapes, etc.). While quick snapshots can be taken directly from the Viewport (as explained above), producing high-quality images or animations requires rendering through Blender's render engines Cycles (path tracing) or EEVEE (fast rasterized rendering). However, doing so requires some familiarity with Blender's more advanced tools. To simplify this process, we provide an additional file, *default_empty.blend* file, which we used for our advanced visualizations and high-quality renderings. It can be downloaded here: https://zenodo.org/records/15826991.

When using this file, multiple visualization options become available for spheres and tracks, respectively. These options can be accessed via the Modifier Properties panel (see Supp. Fig. 18), located in the Properties Editor and highlighted with a pink circle. Clicking the icon expands the panel (see pink frame in Supp. Fig. 18), allowing users to enable or disable spheres and tracks, adjusting Sphere Radius and Resolution, and modifying Sphere Displacement to create more deformed sphere shapes if desired. The Time Size Falloff option allows to gradually shrink spheres over time. Additional parameters such as track length (incoming and outgoing), track width, and interpolation resolution can also be defined. An easy to use cropping functionality can also be accessed at the bottom of the panel.

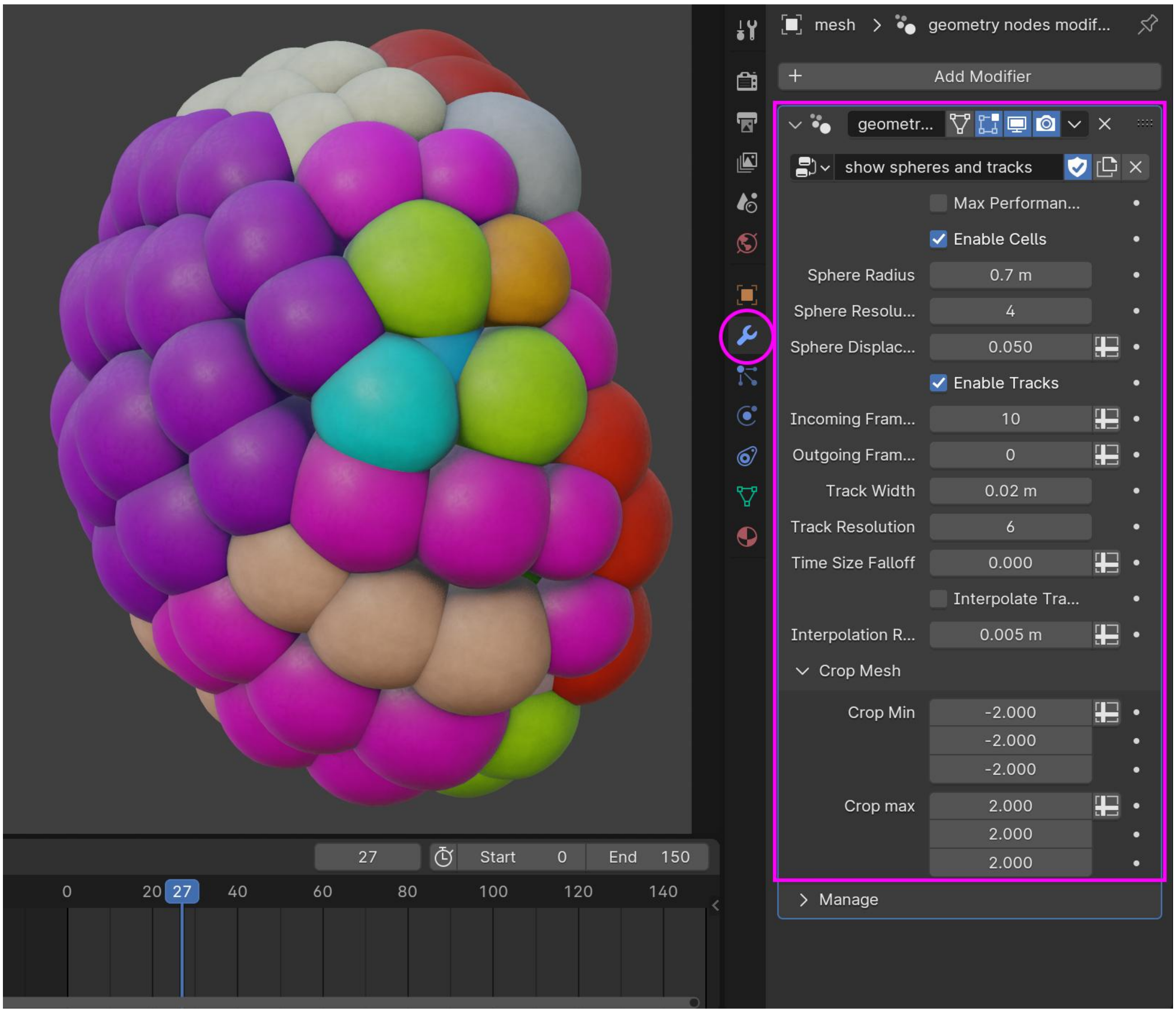


***Supp. Fig. 18:*** Visualization options provided by our custom 'default_empty.blend' file. The Modifier Properties panel (located in the Properties Editor and highlighted with a pink circle) can be expanded by clicking it. This panel (see pink frame) allows you to adjust the sphere size and subdivisions of spots, toggle track visibility, and modify track width as well as the number of frames shown before and after the current time point.

In the advanced visuals Blender view, any scene manually adjusted by the user can serve as a reusable default template for other Mastodon projects by simply saving it with Blender (**File > Save as**). We recommend deleting the initially displayed dataset in the Blender scene to create a clean default template that can be reused with other Mastodon datasets. To do this, **left-click** the mesh in the 3D viewport, press the '**X**' key, and confirm by selecting **Delete**. Alternatively, one can intentionally use a .blend file that includes a mesh from any Mastodon project (e.g., embryo A) as a starting point. In this way, additional datasets (e.g., embryo B) can be added to the same Blender scene and multiple Mastodon projects visualized and rendered together. **Supp. Fig. 19** depicts how a new .blender template can be selected.

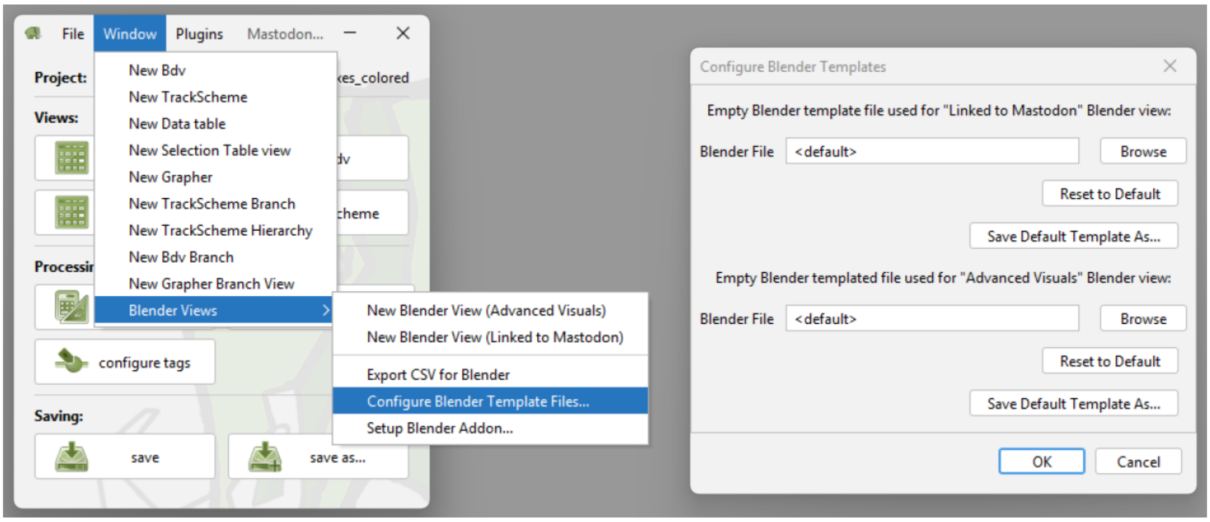

***Supp. Fig. 19:*** Accessing the 'Configure Blender Templates' window to change

#### Visualizing very large tracking datasets using point clouds:

Furthermore, the advanced visuals Blender view supports rendering of large-scale datasets by enabling the 'Max Performance' option. This mode takes advantage of Blender's point cloud rendering, which is optimized for large data. In this case, the Viewport Shading must be set to *Rendered* (Supp. Fig. 20, pink circle), and the render engine must be set to *Cycles* (Supp. Fig. 20, pink arrow and pink box), as the *Material Preview* shading uses the EEVEE engine, which cannot render points in Blender 4.1 (the version we tested thoroughly). However, starting from Blender 4.5, Eevee supports point clouds, rendered as diamond shapes. To ensure optimal performance, a system with an Nvidia graphics chip is recommended for GPU-based rendering.

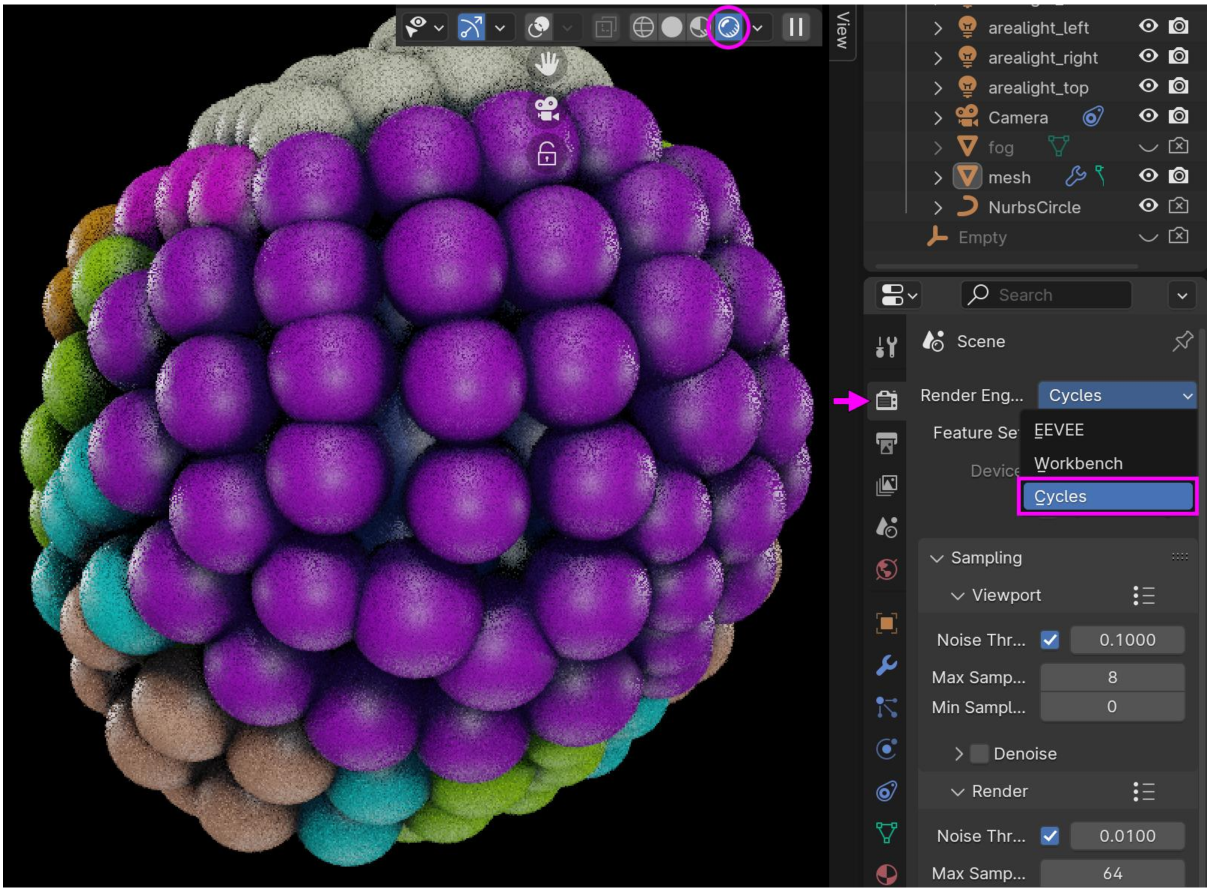


***Supp. Fig. 20:*** Point cloud rendering in Blender with 'Max Performance' enabled. This requires switching to the *Cycles* Render (pink circle) Engine and setting Viewport Shading to Rendered (pink arrow and pink box). Once enabled, millions of annotations can be visualized.

#### 4. Synthetic data generation via Mastodon's *Simulator*

Initially developed to benchmark Mastodon's data handling capabilities and challenge its theoretical limits, a simulator for a growing cell population was implemented directly within Mastodon. The simulator is technically an agent-based system in 3D Cartesian space, where agents are defined as rigid spheres with varying radii and fixed mass.

In each round of the simulation, agents change their positions as a result of contributing individual displacements, according to their own desired movement mode and in response to their immediate surroundings. For example, a randomly varying desired displacement is assigned to an agent in each round to simulate Brownian motion. To prevent "physical" overlap and maintain a small stay-away distance, each agent scans its immediate neighborhood for the presence of other agents, including those nearby. For each detected neighbor, a pair of repelling, distance-driven displacements (the closer the neighbor, the larger the displacement) is assigned to both the scanning and the detected agent.

A simulation round ends when all agents have calculated their resulting displacements and updated their positions. Then, a new simulation round begins. After a defined number of rounds, new Mastodon spots are created at the next time point, each linked to the previously inserted spot of the same agent, thereby constructing the lineage over time.

Agents implement a basic cell cycle and are capable of dividing. Each agent tracks its age, which increases with every new emission of Mastodon spots. Its lifespan is assigned at its birth by drawing from a normal distribution. Based on age progression, agents transition through different behavioral modes.

For example, newly created agents resulting from cell division (as opposed to those introduced at the start of the simulation) initially execute a straight, rapid "bulldozing" movement, which over a few time points decelerates to a normal pace. After this initial phase, agents switch to the default Brownian motion mode with constant size. During the "bulldozing" phase, agents respond only to those they are already colliding with and do not maintain any avoidance distance. But the surrounding (non-"bulldozing") agents maintain the distance. This setup allowed us to simulate the squeezing of daughter cells into dense environments and their gentle reintegration.

As agents approach the end of their lifespan, they attempt to divide (replacing themselves with two daughter agents), provided that their local neighborhood is sparsely populated and the daughters can be placed without overlapping existing agents. These division attempts continue even after the agents exceed their lifespan until the maximum allowed lifespan (which is a configurable simulation parameter and can be disabled entirely). If the agent reaches its lifespan limit, it is understood to undergo prolonged "starvation" and is removed from the simulation.

The simulator maintains an internal flat list of agents, along with their full states, fully autonomously; however, geometrical queries are outsourced to the Mastodon API. Since the complete state of an agent, regardless of behaviour mode, has a small memory footprint, sweeping through and updating a large number of agents is computationally feasible even without utilizing advanced data structures. This operation falls into a linear complexity class with respect to the number of agents. On the other hand, determining which agents fall within a given spatial range becomes easily computationally expensive if implemented straightforwardly, due to quadratic (worst-case) complexity with respect to the number of agents.

To address this, the intermediate spatial arrangements of agents are regularly exported to Mastodon, copied into its internal structures, as the primary purpose of this simulator is to generate content for Mastodon. This data flow design enables the simulator to utilize Mastodon's spot querying infrastructure and API without requiring any additional copy operations. Thus, using the API introduces no computational delay; on the contrary, it improves performance because Mastodon utilizes kd-trees for efficient spatial searches, which have a logarithmic complexity.

Note that the simulation itself actively examines Mastodon's ability to manage and query large-scale annotations in real-time (see '*Simulation of spherical cell populations and performance evaluation*' for an example with 22 million spots).

Interestingly, when agents scan their neighborhoods to determine potential repelling forces, they obtain lists of nearby spheres, derived from discovered neighboring spots, rather than lists of the simulator's own agent objects. This means that these spheres need not come only from spots created by the simulator itself, but they can also include spots that were manually added to Mastodon before the simulation began. Simulation agents are aware of all existing spots in the Mastodon space at each time point and actively avoid them.

This allows a unique interplay between pre-existing data and newly simulated lineages, enabling the creation of hybrid or mixed lineage trees. For example, simulations can be initialized from existing spots, allowing for the artificial extension of pre-existing lineage trees. Alternatively, a population can be simulated growing under one regime for a defined time period, then switched to a different regime in a subsequent simulation session, enabling controlled phase changes in the simulation dynamics. It is also possible to generate spatially denser lineages by simulating new trees alongside, or even within pre-existing ones. Moreover, spots that remain stationary in space but persist over several time points can act as spatial barriers (to be removed after the simulation is over), shaping or constraining the spread and growth of the simulated population as we demonstrate below in the section '*Generating a Gastrulation-like Simulation*'.

In summary, Mastodon's Simulator combines realistic agent-based modeling with seamless integration into the platform's scalable architecture. It supports dynamic lineage generation, efficient force-based interactions of agents, and direct interoperability with existing annotations. Its modularity and performance make it a robust tool for benchmarking, visualization, and synthetic lineage generation on a massive scale.

##### Creating artificial datasets:

We developed two distinct methods for generating large-scale synthetic datasets in Mastodon: agent-based simulations and a cubic-volume lineage generator.

The latter generates perfectly regular cubic spot arrangements, ideal for scalable, controlled benchmarking, with the option to link spots into lineages or leave them unconnected. Instead of agent-based simulation, the dataset is generated using simple Fiji commands.

###### *Generating cubic volumes with regular lineage topology*

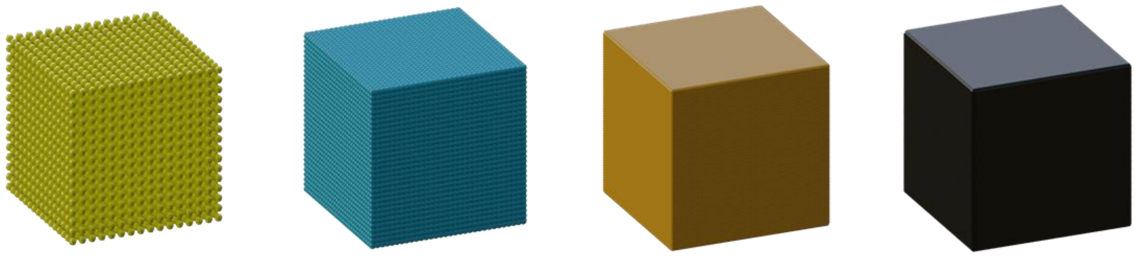

***Supp. Fig. 21:*** Cubic volume datasets generated without an agent-based simulation.

Our cubic volume datasets, designed specifically for benchmarking Mastodon, can be recreated in just a few simple steps using a Fiji installation that includes *Mastodon* and the Mastodon *Simulator*. To enable them, the following update sites have to be enabled in Fiji (**Help > Update > Manage Update Sites**):

- Mastodon (https://sites.imagej.net/Mastodon-jungle/)

- Mastodon-Simulator (https://sites.imagej.net/Mastodon-Simulator/)

After restarting Fiji, navigate to **Plugins > Tracking > Mastodon > Simulator** to download the Mastodon reference projects into a folder. These include empty .mastodon files on which artificial tracks can be generated using custom parameters. To generate the cubic volume datasets using a simple Python snippet in Fiji (Jython) one can then navigate to **Plugins > Tracking > Mastodon > Simulator > Generate regular cubes with lineage** and select an empty .mastodon reference project (e.g. reference_empty_dummydata.mastodon or reference_empty_pixeldata.mastodon). A triaxial growth cycle number of 7 results in 22 cubes distributed across 21 timepoints, with the final cube containing more than 2 million spots with a regular (perfect) lineage. After the dataset is generated, Mastodon's main window opens. Note that selecting higher values will generate massive datasets and significantly increase computation time.

##### *Generating a Gastrulation-like Simulation*

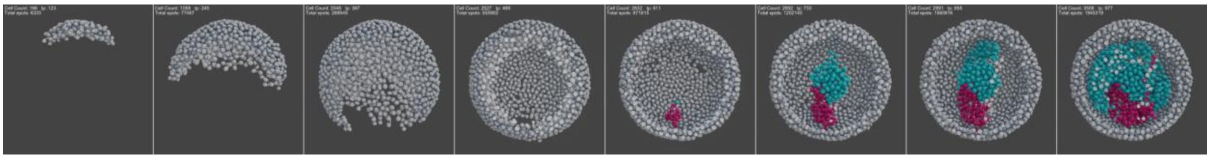

***Supp. Fig. 22:*** Example of a gastrulation-like simulation that requires scene setup prior to running an agent-based simulation with Mastodon's Simulator.

To simulate more dynamic cellular behaviors such as epiboly and gastrulation, we first added specific scene elements, consisting of smaller and larger spherical spots, into an initially empty mastodon project. Examples include layers of hollow spheres, walls or spots placed within these spheres to act as barriers, preventing the simulated dividing 'cells' (which we refer to as 'agents') from trespassing.

##### Creating a Scene for the Gastrulation-like Simulation:

The outer spherical hull, composed of 3,446 spots, was generated instantly inside the BDV using the '**Create Volume Spots**' command. Each spot was assigned a radius of 8 pixels with a 2-pixel overlap, based on an initial manually placed spot with an approximate radius of 200 pixels.

The mid hull, located beneath the outer hull, and the inner hull, forming the deepest layer, were created using the same method. A single hole was introduced at the bottom of the mid hull, and several holes were added to the inner hull by manually deleting selected spots (see **Supp. Fig. 23**).

These different 3D elements were initially created within a single frame, then immediately tagged and colored, as shown in **Supp. Fig. 23**. We used the Selection Creator Parser (SCP) to select all individual spots for each newly created scene element. For example, we first created a new TagSet named 'Scene1', then generated the outer hull spots as described above and tagged them as ‘outer_hull’. Subsequent elements were created and tagged within the same TagSet using distinct names and colors.

To duplicate, for instance, all outer hull spots across the entire 999-frame timeline, we selected only the spots from the original frame using the SCP with the following expression: vertexFeature('Spot frame') == 1 & tagSet('Scene1') == 'outer_hull'. We then duplicated the selected spots as linked spots across all frames using the ‘**Duplicate spots**’ command found under **Plugins > Spots Shuffling**.

The entire scene was created using the method described above, except for the addition of blocking spheres and walls. Blocking spheres were added manually by placing a single spot in the BDV window, renaming it ‘stay_out’, and duplicating it across all frames. Blocking walls were generated by briefly running the simulation on manually placed seed spots in the original frame, with the simulator restricted to 2D (along the x, y, or z axis). During the simulation, these seeds expanded into thin, 2D-like walls (**Supp. Fig. 23**).

##### Running the simulation:

After completing the scene, a single agent was seeded between the outer and inner hulls, with a radius of 6.345 pixels and the simulator was run for 999 frames using the following parameters:

| **Category** | **Parameter** | **Value** |
|---|---|---|
| Agent Spot Size | Initial agent radius | 6.345 |
| Agents Mobility Parameters | Look-around distance | 6.0 |
| Agents Mobility Parameters | Min distance to other agents | 5.0 |
| Agents Mobility Parameters | Usual step size (movement distance) | 4 |
| Agents Mobility Parameters | Max attempts to make a move | 4 |
| Agents Life-cycle Parameters | Average lifespan before division | 8 |
| Agents Life-cycle Parameters | Max lifespan (dies after) | 601 |
| Agents Life-cycle Parameters | Max local density to allow division | 2 |
| Agents Life-cycle Parameters | Max variability of division planes | 2.35 |
| Offspring Agents Settings | Initial distance after division | 0.4 |
| Offspring Agents Settings | Dozering (repulsion) distance | 1.5 |
| Offspring Agents Settings | Dozering time period | 1 |
| Simulation Debug Options | Each time point, add centre spot of agents | Unchecked |

***Table 4:*** Parameters for generating a gastrulation-like simulation

For the first 531 frames, only epiboly-like movements were observed, as the agent’s offspring divided and expanded within the confined space between the two hulls (see BDV in **Video 3**). This phase was followed by gastrulation-like dynamics, during which dividing agents slipped through the hole of the mid hull and began filling the hollow space within and around the inner hull (schematically indicated by pink arrows in **Supp. Fig. 23**).

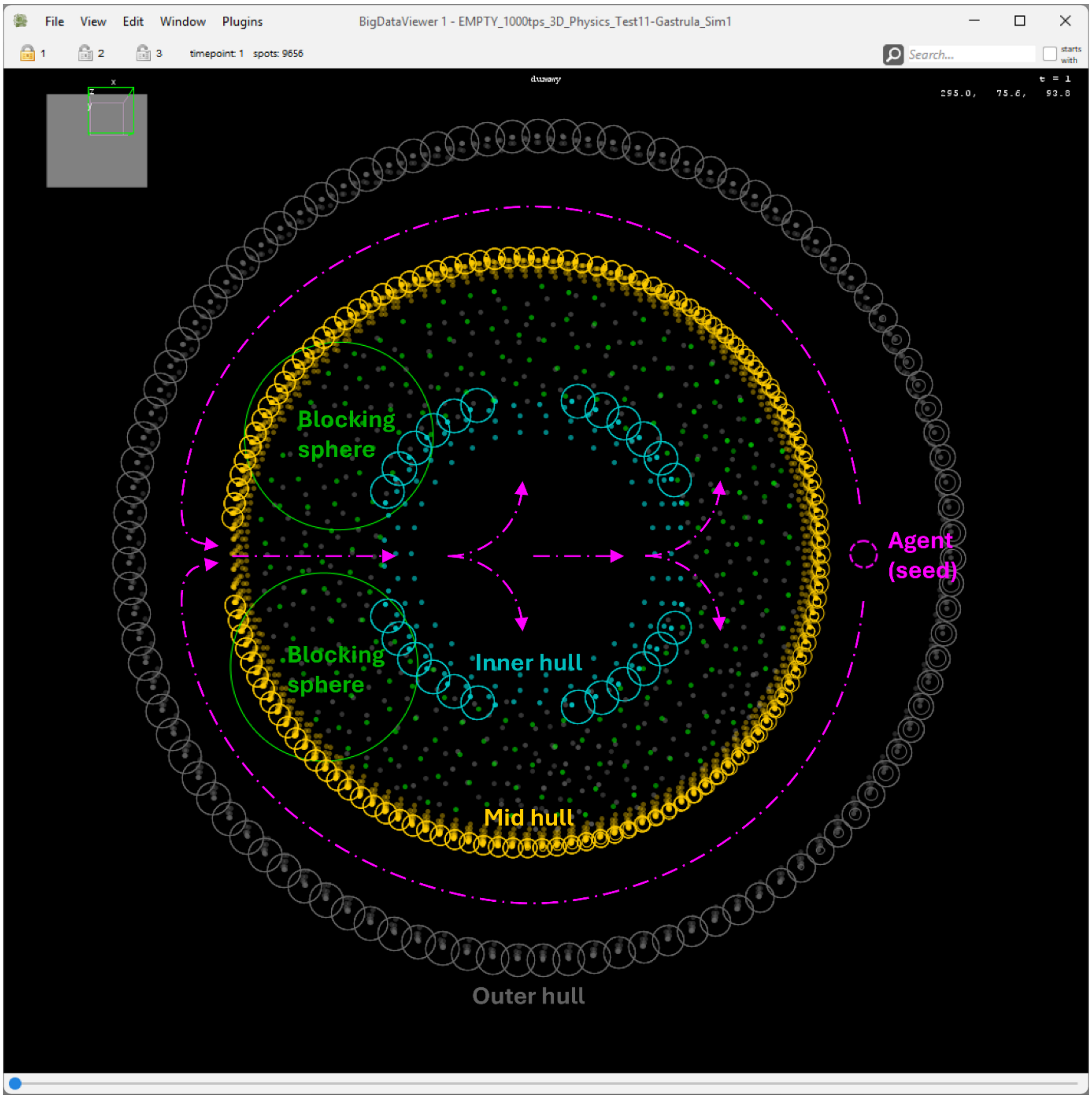


***Supp. Fig. 23:*** Scene setup prior to running an agent-based simulation with Mastodon's Simulator. Pink arrows indicate the path along which dividing agents will spread over 999 frames, confined by 3D scene elements such as the outer, mid, and inner hulls, as well as blocking spheres and barrier walls. The latter are out of focus but visible as green dots.

After the simulation completed, we used the SCP again to select and delete all spots that made up the scene, retaining only the agents (the simulated 'cells' or spots) whose dynamic behavior originated solely from the initially seeded agent. The result was saved as a new Mastodon project.

#### *Simulation of spherical cell populations and performance evaluation*

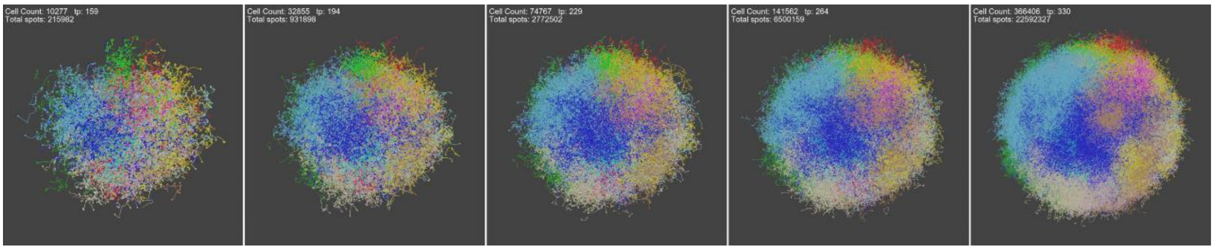


***Supp. Fig. 24:*** Spherical cell population simulated to grow to a total of 22.6 million spots over the entire frame span.

To recreate a lineage tree exceeding 22.6 million spots, we ran Mastodon's Simulator from scratch starting with a single agent (seed) on our reference notebook using the following parameters:

| **Category** | **Parameter** | **Value** |
|---|---|---|
| Agent Spot Size | Initial agent radius | 1.5 |
| Agents Mobility Parameters | Look-around distance | 4.0 |
| Agents Mobility Parameters | Min distance to other agents | 9.0 |
| Agents Mobility Parameters | Usual step size (movement distance) | 9 |
| Agents Mobility Parameters | Max attempts to make a move | 6 |
| Agents Life-cycle Parameters | Average lifespan before division | 10 |
| Agents Life-cycle Parameters | Max lifespan (dies after) | 30 |
| Agents Life-cycle Parameters | Max local density to allow division | 2 |
| Agents Life-cycle Parameters | Max variability of division planes | 2.4 |
| Offspring Agents Settings | Initial distance after division | 1.6 |
| Offspring Agents Settings | Dozering (repulsion) distance | 1.5 |
| Offspring Agents Settings | Dozering time period | 2 |
| Simulation Debug Options | Each time point, add centre spot of agents | Unchecked |

***Table 5:*** Parameters for the simulation of spherical cell populations

The generated dataset approaches Mastodon's current upper limit of 23,342,213 annotations, a constraint imposed by its use of a fixed-size array data structure (SingleArrayMemoryPool). Exceeding this limit without switching to an alternative implementation (such as an experimental MultiArrayMemoryPool version of Mastodon) will result in a simulator crash.

##### Simulator performance

To evaluate the performance of the simulator under near-maximum load, three independent runs were performed using identical parameters on the reference notebook (**Supp. Fig. 25**). Each simulation was manually stopped after surpassing 22 million agents (linked spots).
Across these highly consistent runs, the mean execution time was 5 minutes and 15.3 seconds (315.3 seconds), demonstrating the simulator's ability to efficiently generate large, artificially constructed lineages (see Table 6).

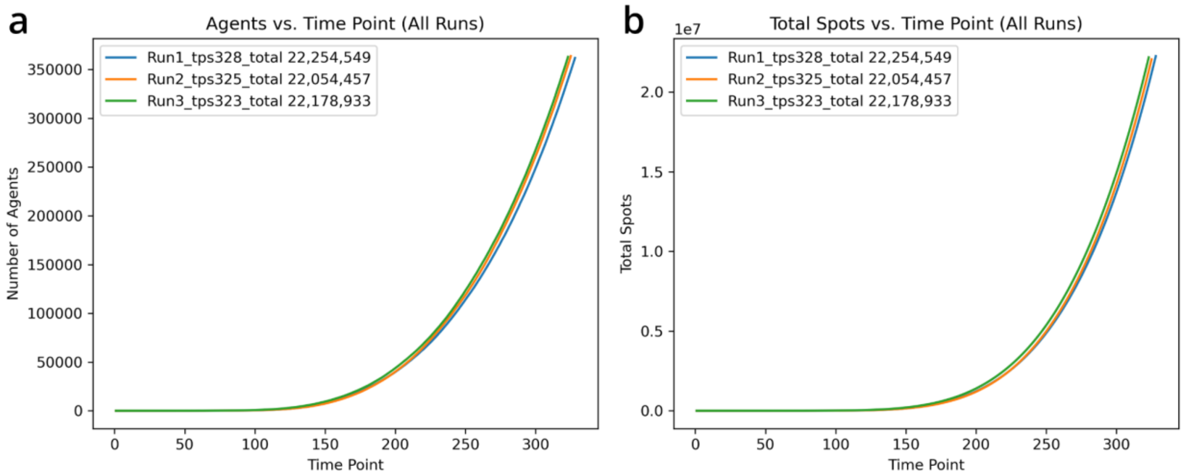


***Supp. Fig. 25:*** Three independent simulation runs (blue = Run1, orange = Run2, green = Run3), each generating over 22 million linked spots (agents). (a) Maximum number of artificially generated annotations (agents) per time point. (b) Cumulative total number of artificially generated annotations per time point.

| Run | Final Time Point (TP) | Number of Agents (final TP) | Agents (total) | Simulation Duration |
|---|---|---|---|---|
| Run 1 | 328 | 361816 | 22254549 | 5 min 15 sec |
| Run 2 | 325 | 363742 | 22054457 | 5 min 4 sec |
| Run 3 | 323 | 362985 | 22178933 | 5 min 27 sec |

***Table 6:*** Summary of outcomes from three simulation runs.

All three runs follow nearly the same curve, showing highly consistent simulation performance. The simulation time increases slowly at first (flat curve until ~ time point 120), then accelerates as the number of agents grows, indicating a rising computational load per time point (**Supp. Fig. 26**).

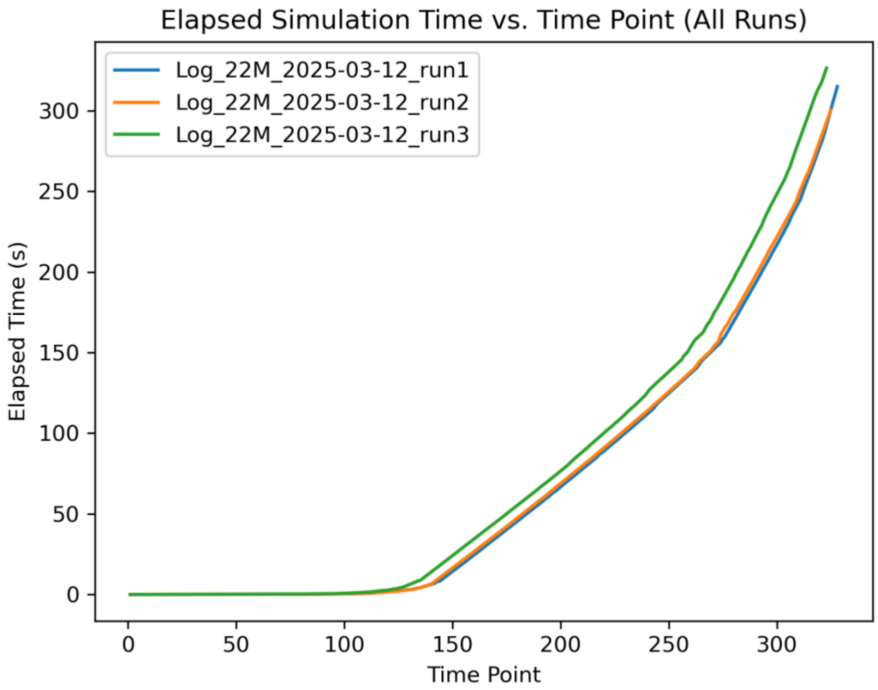

***Supp. Fig. 26:*** Time that Mastodon's Simulator takes to compute successive time points across three independent runs (blue = Run1, orange = Run2, green = Run3), each generating over 22 million linked spots (agents).

Notably, the simulator successfully handled over 22 million agents in approximately 315 seconds (around 5 minutes and 15 seconds). This demonstrates that, despite simulating millions of agents with independent state tracking, real-time interactions, and spatial awareness of all annotations, which together enable realistic proliferation and motion in large 3D volumes, the simulator can generate over 22 million linked annotation spots in under five and a half minutes on a standard laptop. This performance allows users to run large-scale, biologically inspired simulations without the need for clusters or GPUs, and to generate complex lineage scenarios or massive synthetic datasets for benchmarking and training.

##### *Simulating lymphocyte motility using MotilityLab*

Lymphocyte motility tracks were generated using the online tool MotilityLab Track Generator (http://www.motilitylab.net/beauchemin.php) with parameters listed in **Table 7**. The simulated data were then imported into Mastodon via the CSV importer.

| **Parameter** | **Value** |
|---|---|
| Free velocity (vfree) | 120 µm/min |
| Free run time (tfree) | 20.0 min |
| Pause time (ttpause) | 0.5 min |
| Time step (deltat) | 1 min |
| Total duration | 18 min |
| Image region (Z) | 100 µm |
| Simulation volume | 500 x 500 x 500 µm$^3$ |
| Motility coefficient | 46829.27 µm$^2$/min |
| Number of cells | 1000 |
| Parameter | Value |
| Free velocity (vfree) | 120 µm/min |
| Free run time (tfree) | 20.0 min |
| Pause time (ttpause) | 0.5 min |
| Time step (deltat) | 1 min |
| Total duration | 18 min |
| Image region (Z) | 100 µm |
| Simulation volume | 500 x 500 x 500 µm$^3$ |
| Motility coefficient | 46829.27 µm$^2$/min |
| Number of cells | 1000 |

***Table 7 - MotilityLab Simulation Parameters***

#### 5. Performance benchmark for rendering annotations

We benchmarked the rendering performance of lineage visualizations in both the BDV and TrackScheme views to quantitatively assess responsiveness and determine usability limits across different datasets and rendering configurations. Furthermore, because this benchmark is made available directly in Fiji, users can easily evaluate the performance of their own hardware and can determine the project sizes it can handle comfortably.

To install the Mastodon-Benchmark feature in a Fiji distribution, activate the following Fiji update site: Help > Update… > Manage Update Sites > Mastodon-Benchmark https://sites.imagej.net/Mastodon-Benchmark/.

While previous work has shown that large images can be rendered efficiently using the BDV [12], we aimed to evaluate how well Mastodon handles the rendering of large numbers of spot and track annotations. Rendering annotations without image data not only isolates the performance impact of overlays, but also reveals the upper limits of Mastodon's internal spot and track data structures. Specifically, we quantify how long it takes Mastodon to retrieve relevant spots, compute their graphical representation, and display them.

The Mastodon benchmark is implemented in a straightforward manner. A sampling timestamp is recorded immediately before a drawing request is issued to either BDV or TrackScheme, and again after the rendering is finished, essentially a stopwatch-based approach. The elapsed time between these two points represents the gross (or wall-clock) time needed to complete the request and is the value reported in the benchmark.

This measurement reflects the actual user experience, as it includes potential delays caused by external factors such as operating system multitasking. Importantly, in all benchmarks presented in this paper, each drawing request was issued to exactly one display window, either a single BDV or a single TrackScheme view. Due to the multithreading (parallel) nature of software components of Mastodon, when multiple display windows have been requested, we are not able to reliably report their individual measurement times. Instead, only a total time needed to complete is available, but this cannot be used to estimate time needed when, for example, a different number of BDV windows had been used.

Implementing the benchmark required modifications to both the Mastodon and BDV codebases. Consequently, the benchmark update site replaces the default BDV in the Fiji installation, which may affect other plugins that rely on it.

A known limitation of the benchmark is that it cannot reliably report wall times when additional repaint events occur. For example, when the operating system triggers a redraw after the user moves the mouse cursor over the BDV. For this reason, it is recommended that the computer remain untouched while the benchmark is running. Finally, TrackScheme view provided by the benchmark update site includes support for placing and retrieving view bookmarks, similar in concept to those available in BDV.

To ensure both flexibility and reproducibility, a dedicated benchmark command language was developed for Mastodon. This language allows users to define a sequence of commands, each instructing one or more display windows to render a specific view of the data.

Commands are executed sequentially, with each new command issued only after the previous rendering is complete. The rendering time for each command is recorded and reported. For consistency and performance measurement, standard Mastodon commands lack animation, ensuring that each requested view is established in a single rendering pass.

In short, a benchmark run is defined by providing the following components:

1. A .mastodon project file containing only lineage data (no image pixel data), typically of substantial size to place meaningful stress on the system.
2. A separate .txt file containing predefined TrackScheme bookmarks, since this information cannot be stored within standard .mastodon files.
3. The benchmark opens a single BigDataViewer and one TrackScheme view.
4. An initialization command (in the benchmark language) that sets all views to defined initial views.
5. A benchmark command sequence that drives the actual test.

This setup can be shared across different Mastodon installations and operating systems, with the resulting benchmark times remaining directly comparable.

##### Setting up and using the BDV window for benchmarking

###### *Rendering Options for BDV window and TrackScheme window when running the Benchmark*

In the 512x512-pixel BDV window used throughout the benchmark, we compared two rendering options: the **default (built-in)** settings and a customized version, which we refer to as "**Basic**." The default BDV render settings display ellipsoid projections and include tracking links within a defined time range. Importantly, a focus limit of 100 is applied, meaning that only the portion of the tracking annotations within this defined volume, visible in the BDV window's field of view, is rendered.
In contrast, our customized "Basic" BDV render settings effectively remove the focus limit by setting it to the maximum value (2000). However, in the Basic settings, neither tracking links nor ellipsoids are shown. Instead, each annotation vertex is represented by a single central point. The specific settings used throughout the benchmark are shown in **Table 8**.

| BDV Render Settings | Default (built-in) | Basic |
|---|---|---|
| Anti-aliasing | Enabled | Enabled |
| Draw links | Enabled | Disabled |
| Time range for links | 20 | 15 |
| Gradients for links | Disabled | Disabled |
| Arrow heads | Disabled | Disabled |
| Draw links ahead in time | Disabled | Disabled |
| Link stroke width | 1 | 1 |
| Draw spots | Enabled | Enabled |
| Ellipsoid intersection | Enabled | Disabled |
| Ellipsoid projection | Enabled | Disabled |
| Draw spot centers | Enabled | Enabled |
| Draw spot centers for ellipses | Disabled | Disabled |
| Draw spot labels | Disabled | Disabled |
| Spot stroke width | 1 | 4 |
| Fill spots | Disabled | Disabled |
| Focus limit (max dist to view plane) | 100 | 2000 |
| View relative focus limit | Enabled | Enabled |
| Ellipsoid fade depth | 0,2 | 1 |
| Center point fade depth | 0 | 1 |

***Table 8:*** Default BDV Render Settings - Default (built-in) and Basic

##### *Creating Bookmarks in the BDV Window*

Anything outside the field of view of the BDV window will not be rendered by the API, which has a significant impact on performance. This situation can arise, for example, when a tracking dataset rotates or expands over time, causing some trajectories to move outside the visible area. To ensure consistent benchmarking results, we created bookmarks that encompass the entire dataset, unless a specific subset of tracking data is intentionally targeted for the benchmark.

The ImageJ documentation provides detailed instructions on how to set bookmark locations and orientations with the BDV, as well as how to load and save these settings as an XML file (see https://imagej.net/plugins/bdv/).

In brief, a bookmark of the current view can be created by pressing Shift + B, followed by the desired shortcut key for that bookmark. Saving bookmarks as an XML file is then available under File > Save Settings.

Note that the settings file must be placed in the same directory as the dataset's XML file and must share the same filename, with the addition of the .settings.xml extension (e.g., mynewbookmarks.settings.xml). Its file path must also be specified in the Benchmark pop-up window under 'BigDataViewer display settings.xml file'.

##### *Using BDV Bookmarks During the Benchmark*

During a benchmark run, previously created BDV bookmarks can be recalled using the input line for benchmark commands:

- **Jump to a specific bookmark**:
    - BDV_Bn

  where n is the bookmark number.

- Multiple BDV views opened using the Benchmark launcher are each labelled as BenchBDV#n. Individual windows can be addressed using the following command:
    - BDVn

  where *n* corresponds to the BDV window number (#n). To address a specific BDV, *n* can be replaced with the appropriate number.

##### *Jumping to frames using the BDV during the Benchmark*

- **Jump to a specific frame (timepoint) using the following command.**
    - BDV_Tn

  Where n represents the frame/timepoint.
- **Rotate the tracking data within a specific frame (timepoint) using the following command.**
    - BDV_Rn

  Where *n* represents the total number of steps for a full 360° rotation.

To create a frame sequence, commands must be repeated for each frame in sequence. For example: `BDV_T1 BDV_R5 BDV_T2 BDV_R5.`
In this example sequence, an open BDV would jump to frame 1 (`BDV_T1`), perform a full 360° rotation in 5 steps (`BDV_R5`), then jump to frame 2 (`BDV_T2`) and perform another full rotations in 5 steps (`BDV_R5`).

#### Setting up and using the TrackScheme window for benchmarking

##### *Automated Zoom Operations in the TrackScheme Window*

To evaluate performance, automated zoom-in and zoom-out operations were performed within a 1024x512-pixel TrackScheme window.

##### *Creating Bookmarks in the TrackScheme Window*

Before creating bookmarks, the benchmark must be started to ensure that at least one TrackScheme window is opened. Note: TrackScheme windows launched independently (not from the benchmark starter window) are not supported for bookmarking.
Once the benchmark is running, it can be paused to generate bookmarks using the following commands:

- **Create a bookmark**: Hold **Shift** and press a number key (**n**) to save the current TrackScheme view as a bookmark.
- **Save all bookmarks**: Hold **Shift** and press **Q** to write the bookmarks to a file named 'bookmarks_trackscheme.txt' in the same folder where the BDV settings XML file is stored.

##### *Using TrackScheme Bookmarks During the Benchmark*

During a benchmark run, previously created TrackScheme bookmarks can be recalled using the input line for benchmark commands:

- **Jump to a specific bookmark**:
  TS_Bn

where *n* is the bookmark number.

- **Animate zoom between two bookmarks**:
  TS_Zn-m-f

where:

- *n* is the starting bookmark,
- *m* is the ending bookmark,
- *f* is the number of frames for the zoom animation

This command defines a zoom animation from one view to another. For example, a zoom-in effect can be achieved by starting with a wide view (e.g., showing all tracks) and ending with a close-up view (e.g., displaying only a few spots and links). To return to the original view, the zoom animation can be reversed by swapping the start and end bookmarks: `TS_Z1-3-10 TS_Z3-1-10`.

This example sequence above performs a 10-frame zoom-in followed by a 10-frame zoom-out, and returns the TrackScheme view to its initial state.

##### Creating complex command sequences using a Python script.

We created a Python script that automatically generates a sequence of commands to navigate and animate a series of time points in Mastodon's BDV and TrackScheme views. The script is shipped with the Mastodon-Benchmark Fiji update site: https://sites.imagej.net/Mastodon-Benchmark/.

It gives the following options:

- Go through selected timepoints (BDV_T*n* → BDV_T*n* → BDV_T*n* → *etc.*).
- Show timepoints with a certain frequency (e.g., every *n*th frame).
- Add rotation commands for the BDV (BDV_R*n*).
- Optionally skip some rotation steps for faster animations.
- Include optional bookmarks, zoom-ins, and zoom-outs (TS_Zn-m-f → TS_Zn-m-f).
- Combine all of this into a single output line of commands.

In short, the script builds complex sequences starting like this:
`BDV_T5 BDV_R5 TS_Z1-3-10 TS_Z3-1-10 BDV_T23 BDV_R5 TS_Z1-3-10 TS_Z3-1-10 ...`
Such a sequence can then be used to control the BDV and TrackScheme windows during a benchmark run.

### Acronyms used.

- **API** – Application Programming Interface
- **BDV** – BigDataViewer
- **CPU** – Central Processing Unit
- **CSV** – Comma-Separated Values
- **CTC** – Cell Tracking Challenge
- **GPU** – Graphics Processing Unit
- **GraphML** – Graph Markup Language
- **HDF5** – Hierarchical Data Format version 5

- **MaMuT** – Massive Multi-view Tracker (Fiji plugin)
- **N5** – A hierarchical data storage format used in ImageJ/Fiji ecosystem
- **NURBS** – Non-Uniform Rational B-Splines
- **OME-NGFF** – Open Microscopy Environment Next-Generation File Format
- **PhyloXML** – Phylogenetic eXtensible Markup Language
- **RAM** – Random Access Memory
- **SCP** – Secure Copy Protocol (mentioned in code/usage contexts; appears in Mastodon docs)
- **TGMM** – Tracking with Gaussian Mixture Models
- **TP** – Time Point (sometimes abbreviated in simulation tables)
- **XML** – eXtensible Markup Language

### References.

[7] J. L. Hennessy and D. A. Patterson, *Computer Architecture: A Quantitative Approach*, 5th ed. San Francisco, CA, USA: Morgan Kaufmann Publishers Inc., 2011.

[8] R. Nystrom, *Game Programming Patterns*. Genever Benning, 2014.

[9] T. Pietzsch, J. Y. Tinevez, and M. Aartz, "Mastodon-collection." 2014. Accessed: Apr. 19, 2024. [Online]. Available: https://github.com/mastodon-sc/mastodon-collection

[10] M. Hertz and E. D. Berger, "Quantifying the performance of garbage collection vs. explicit memory management," *ACM SIGPLAN Not.*, vol. 40, no. 10, pp. 313–326, Oct. 2005, doi: 10.1145/1103845.1094836.

[11] T. Pietzsch, J. Y. Tinevez, and M. Arzt, "Mastodon-graph." Mastodon Science, 2014. Accessed: Apr. 22, 2024. [Online]. Available: https://github.com/mastodon-sc/mastodon-graph

[12] Pietzsch, T., Saalfeld, S., Preibisch, S. et al. BigDataViewer: visualization and processing for large image data sets. Nat Methods 12, 481–483 (2015). https://doi.org/10.1038/nmeth.3392
